# Genome-scale metabolic models for a synthetic soil microbial community as a path for understanding community functioning

**DOI:** 10.64898/2026.10.16.759588

**Authors:** Omar S. Keshk, Evangelia Vayena, Senka Čaušević, Anna Sintsova, Davin Saviro Wijaya, Jan Roelof van der Meer, Vassily Hatzimanikatis

## Abstract

Microbial communities drive essential ecosystem processes, yet community-level behavior often deviates from predictions based on single-species studies. This mismatch reflects the context-dependent nature of nutrient dependencies, metabolic interactions, and resource availability in complex environments. Here, we present a mechanistic, species-resolved framework to dissect metabolism and nutrient exchange in a defined 21-member soil synthetic community spanning five bacterial phyla. For each isolate, we reconstructed and manually curated thermodynamically constrained genome-scale metabolic models, reconciling predictions with growth phenotypes across rich and minimal media. We mapped minimal nutrient requirements, cataloged environment-specific uptake and secretion profiles, and compared metabolic pathways to identify potential competitive and cooperative interactions. Using community-scale metabolomics, we assigned putative producers and consumers for detected metabolites and revealed medium-dependent shifts in niche overlap. This approach exposes metabolic strategies that govern community function and resilience, offering a predictive basis for understanding and manipulating microbiomes. Our findings highlight how species-level metabolic maps can clarify why microbial interventions succeed or fail and guide targeted manipulations in soil ecosystems.

## Introduction

Microorganisms inhabit every environmental niche on our planet. Typically, they form communities of species with different relative abundances that interact within a defined space, often with complex spatial structures. Microbial communities, also called microbiota, carry out complex biochemical transformations that strongly shape their habitats^1^. For example, soil microbiota contribute to soil health by fixing nitrogen, suppressing phytopathogens, and recycling carbon and nutrients^2,3^. Understanding how microbial communities function is therefore central to efforts aimed at improving bioremediation, wastewater treatment, and crop productivity^4^.

Yet most of our knowledge still comes from single-species studies that ignore local resource availability and cannot capture interactions such as competition, mutualism, and nutrient sharing; consequently, community-level behavior rarely aligns with pure-culture predictions. Furthermore, communities remain challenging to interrogate at single-species resolution^5^. This gap underscores the need for approaches that emphasize species interactions and resource exchange^6,7^.

Two lines of evidence motivate a mechanistic, metabolism-centered approach. First, nutrient dependencies and cross-feeding among microbial species are often context- and lineage-specific rather than strictly conserved. For example, “who depends on whom” for vitamins is often phylogenetically heterogeneous since vitamin auxotrophies can appear and disappear even among close relatives^8,9^. In soils specifically, the presence or absence of producers of cobalamin (vitamin B12), a type of corrinoid, can reverberate through the microbiome. Yet producer, remodeler, and consumer strains remain poorly characterized because corrinoids are scarce and analytically demanding to quantify in soil matrices^10^.

Second, attempts to enhance community functions by introducing specific strains (bioaugmentation) often fail in complex environments, such as oil-contaminated sites. Lab-designed communities often fail in situ because degraders may switch to alternative nutrients, essential partner microbes that supply required metabolites may be absent, and lab-selected strains may be outcompeted by native microbes^11^. Together, these observations underscore the need for frameworks that map species-specific metabolism and nutrient dependencies within defined environments and link these maps directly to community function.

Genome-scale metabolic models (GEMs) provide such a framework by linking genes, reactions, and biomass requirements into predictive networks^12,13^. Draft models reconstructed from annotated genomes with tools such as RAVEN using KEGG orthologies^14–16^ typically require curation, because ortholog-based inference can leave gaps or introduce spurious reactions^17^. Once curated, GEMs can be validated against growth phenotypes and probed with constraint-based simulation^18,19^ to delineate feasible nutrient uptake and byproduct secretion, identify minimal nutrient sets, and predict gene essentiality.

Recent theory suggests that even complex microbial communities exhibit predictable patterns^11^. By mapping the metabolism and interactions of individual species, we can often predict community behavior from composition, highlighting the value of species-resolved, mechanistic maps of metabolism and interactions.

Here, we integrate these approaches in a defined 21-member soil synthetic community (SynCom)^20^ spanning five phyla (Table 1). We reconstruct and manually curate high-quality, thermodynamically constrained GEMs for each isolate; reconcile model predictions with growth phenotypes on rich and minimal defined media, and systematically map each species’ metabolic traits along three axes directly tied to interaction potential: (i) identifying the minimal nutrients required for growth and highlighting biomass building blocks that cannot be synthesized, (ii) profiling compounds consumed and released under different environmental conditions, and (iii) comparing key metabolic pathways to reveal potential competition or complementarity. Coupling this with community-scale metabolomics, our pipeline translates phenotypes into precise constraints, assigns plausible producers and consumers for detected metabolites, and reveals medium-dependent shifts in niche overlap.

**Table 1:** Taxonomic composition of the 21-member soil synthetic community (SynCom).

| Model code | Assembly identifier | Submission code | GTDB assignment | GTDB phylum | GTDB class | Assembly accession |
| --- | --- | --- | --- | --- | --- | --- |
| bbet | PB02 | Bra | *Bradyrhizobium* | Pseudomonadota | Alphaproteobacteria | GCA_988226255 |
| buol | PB15 | Bur | *Paraburkholderia aspalathi* | Pseudomonadota | Gammaproteobacteria | GCA_988226305 |
| cauj | PB07 | Cau | *Caulobacter* | Pseudomonadota | Alphaproteobacteria | GCA_988226285 |
| cxy | PB01 | Cell | *Cellulomonas* | Actinomycetota | Actinomycetes | GCA_988226295 |
| cpi | PB16b | Chi | *Chitinophaga* | Bacteroidota | Bacteroidia | GCA_988226415 |
| cohm | PB10 | Coh | *Cohnella* | Bacillota | Bacilli | GCA_988226435 |
| cuvk | 018_S8 | Cur | *Curtobacterium*<br>*sp003234365* | Actinomycetota | Actinomycetes | GCA_988226495 |
| dri | PB09 | Dev | *Devosia* | Pseudomonadota | Alphaproteobacteria | GCA_988226505 |
| flk | 005_S3 | Fla | *Flavobacterium* | Bacteroidota | Bacteroidia | GCA_988226525 |
| lur | PB03a | Lut | *Luteibacter* | Pseudomonadota | Gammaproteobacteria | GCA_988226445 |
| lgu | PB06 | Lys | *Lysobacter* | Pseudomonadota | Gammaproteobacteria | GCA_988226385 |
| mlo | PB05 | Mes | *Mesorhizobium* | Pseudomonadota | Alphaproteobacteria | GCA_988226395 |
| mip | 012_S5 | Mic | *Microbacterium*<br>*sp001595495* | Actinomycetota | Actinomycetes | GCA_988226335 |
| mgk | 015_S7 | Muc | *Mucilaginibacter* | Bacteroidota | Bacteroidia | GCA_988226455 |
| pzu | PB04 | Phe | *Phenylobacterium* | Pseudomonadota | Alphaproteobacteria | GCA_988226365 |
| pmor | PB08 | Pse-1 | *Aquipseudomonas*<br>*sp003605735* | Pseudomonadota | Gammaproteobacteria | GCA_988226405 |
| pscf | PB14 | Pse-2 | *Aquipseudomonas paracarnis* | Pseudomonadota | Gammaproteobacteria | GCA_988226325 |
| racd | PB12 | Rah | *Rahnella victoriana* | Pseudomonadota | Gammaproteobacteria | GCA_988226485 |
| rfa | PB13 | Rho | *Rhodococcoides fascians* | Actinomycetota | Actinomycetes | GCA_988226545 |
| tro | PB11 | Tar | *Tardiphaga* *sp008329525* | Pseudomonadota | Alphaproteobacteria | GCA_988226465 |
| vpe | 004_S2 | Var | *Variovorax* | Pseudomonadota | Gammaproteobacteria | GCA_988226355 |

Collectively, these approaches provide a tractable, species-level view of metabolic strategies in soil, offering insights into when and why microbial interventions succeed and how nutrient dependencies shape interactions in soil ecosystems^10,11^.

## Results

### Growth phenotyping reveals metabolic diversity and auxotrophies

The soil SynCom comprised 21 bacterial strains isolated from forest soil, representing Proteobacteria, Actinobacteria, Bacteroidetes and Firmicutes (Table 1). In the original study, the strains were cultured individually and combined at approximately equal starting abundances before inoculation into sterile soil microcosms^20^. Here, we report genome sequencing, assembly, and annotation for all 21 strains, providing the genomic basis for strain-specific metabolic reconstruction (see Methods). To characterize their metabolic capabilities experimentally, we profiled growth on 29 single carbon substrates in defined media and on the rich, undefined medium peptone-tryptone-yeast extract-glucose (PTYG; Fig. 1; Supplementary Figs. S1 and S2). Several Beta- and Gammaproteobacteria, including buol, vpe, pmor and racd, used a broad range of substrates, whereas mlot and mgk used fewer (Fig. 1; strain abbreviations in Table 1). Two strains, mip and cohm, grew measurably only on PTYG, suggesting dependence on biomass precursors that they could not synthesize (auxotrophies).

**Figure 1.**
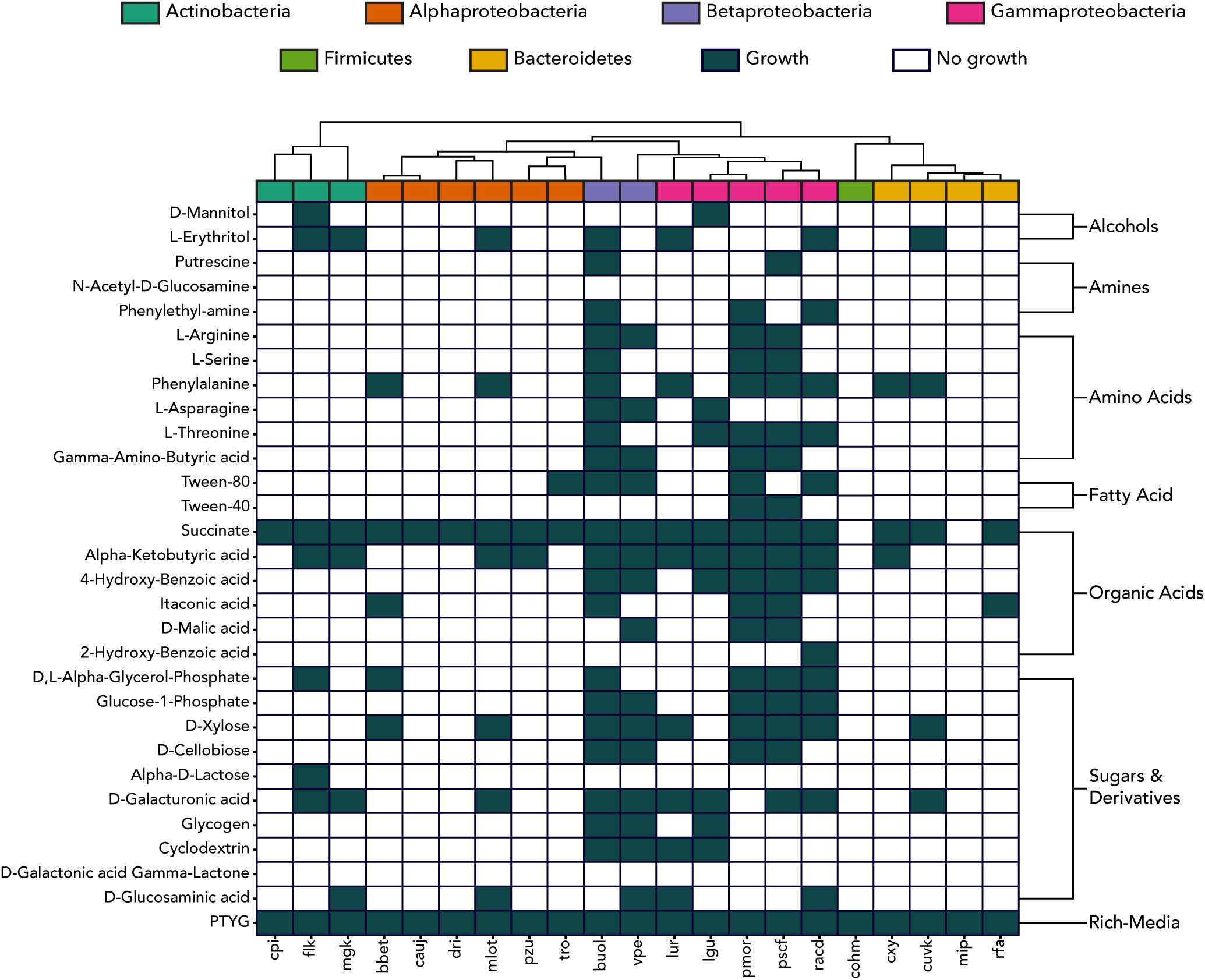
Experimental growth phenotypes across the community. Binary growth matrix across 29 carbon sources (rows) and species (columns). Green denotes growth; white denotes no growth. Species are grouped by phylum (top color strip; acronyms as in Table 1), and substrates are grouped by chemical class on the left. The profiles reveal substantial metabolic diversity among the 21 SynCom strains, ranging from broad generalists capable of utilizing many substrates to auxotrophs restricted to rich medium.

### Genome-scale metabolic models capture species-specific metabolism

To investigate the metabolic basis of these growth phenotypes, we used the genome annotations to reconstruct and manually curate GEMs for all 21 soil SynCom strains (Fig. 2; see Methods). The curated models contained 2,849-3,651 reactions and 2,598-3,414 metabolites per strain. Reaction and metabolic gene counts varied among strains (Fig. 3A) and were positively correlated with genome size (Pearson’s r = 0.74, P = 1.11 × 10⁻⁴; Supplementary Fig. S3A-B).

**Figure 2.**
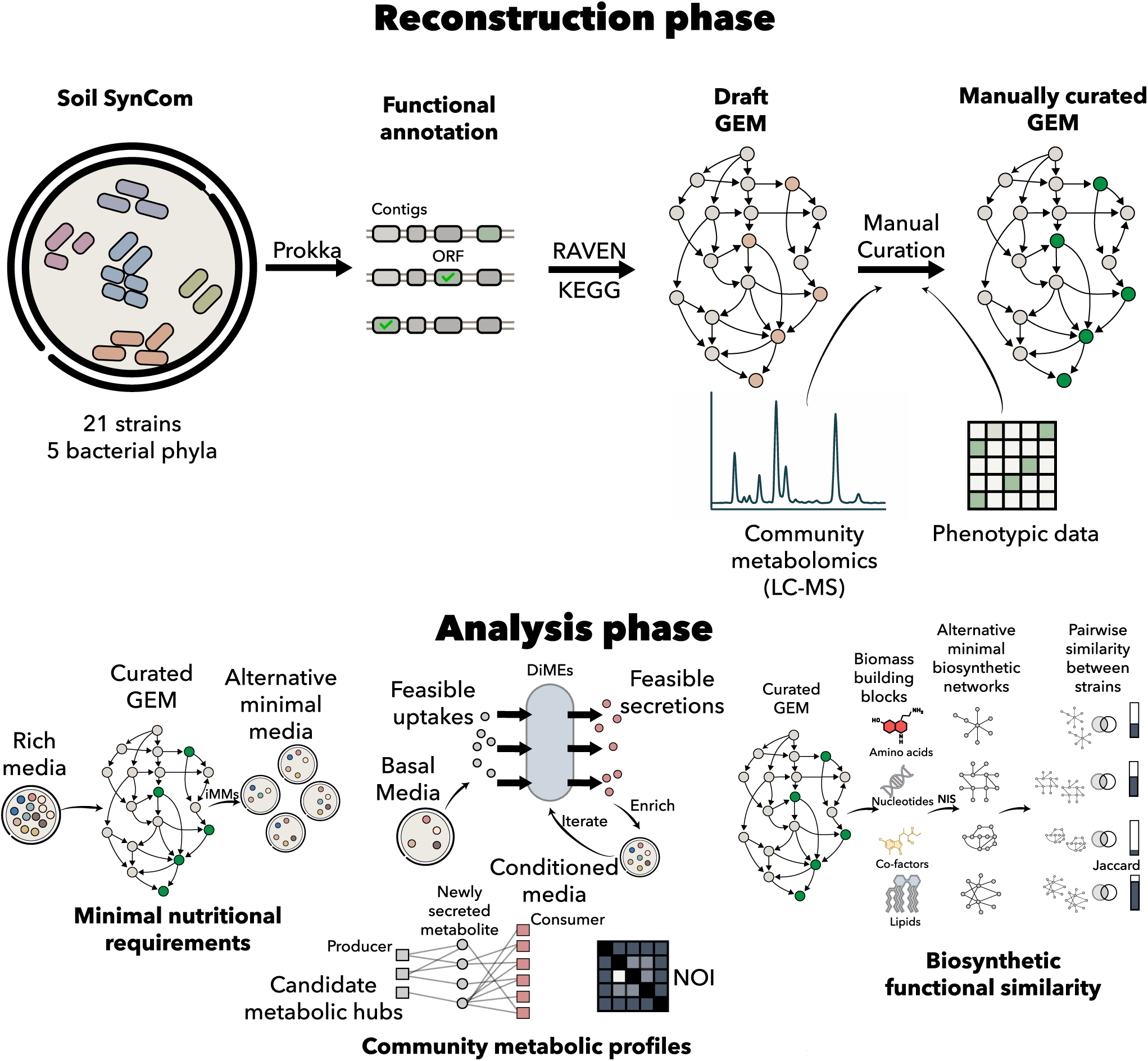
Overview of the genome-scale metabolic modeling workflow for the soil synthetic community. Overview of the experimental and computational pipeline used to reconstruct and analyze genome-scale metabolic models (GEMs) for 21 soil bacterial isolates. The workflow begins with strain isolation, genome sequencing, assembly, and annotation, followed by draft GEM reconstruction. Models are then iteratively curated and gap-filled using NICEgame, guided by experimental growth data. Curated GEMs are subsequently used in three downstream analyses: (i) prediction of minimal nutrient requirements through in silico minimal media simulations, (ii) characterization of uptake and secretion profiles using DiMEs, and (iii) comparison of biosynthetic pathway repertoires using NIS. Together, this integrated pipeline translates genomic information into curated, predictive GEMs that enable mechanistic and species-resolved analysis of metabolic capabilities and interactions within the soil community.

**Figure 3.**
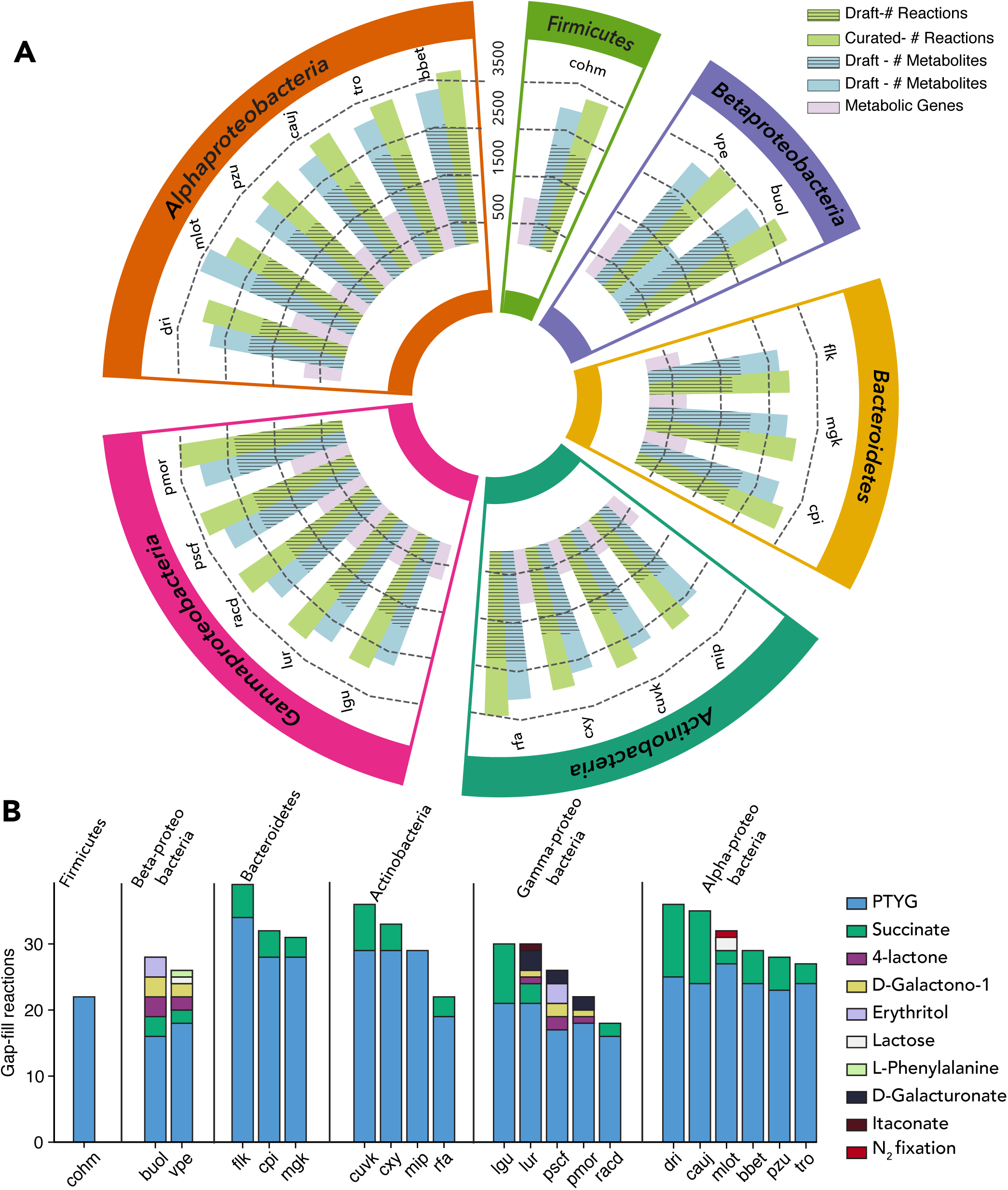
Genome-scale metabolic models of the 21 soil SynCom isolates. (A) Circular summary of genome-scale metabolic model (GEM) content grouped by phylum (outer colored sectors). Within each phylum, isolates are ordered by total number of reactions in the curated model. For each isolate, a single radial bar reports the total number of reactions in the final curated GEM. The dark inner segment corresponds to reactions present in the draft reconstruction, while the lighter outer segment represents reactions added during curation, including exchange reactions, transporters, and gap-filling reactions. Together, the dark and light segments indicate the total reaction content of each curated model. Concentric reference circles indicate reaction count scale. (B) Gap-filling requirements across isolates. Stacked bars show the number of reactions added to restore growth feasibility, partitioned by growth medium. Models are grouped by phylum, ordered from phyla with fewer represented species to those with more species; within each phylum, isolates are ordered from highest to lowest total number of gap-filling reactions. The total bar height represents the cumulative number of reactions added for each isolate.

To reconcile the draft GEM models with the observed growth data, we gap-filled pathways using the NICEgame workflow^21^, first on PTYG and then on defined media with individual substrates if further required (see Methods). For all models, this required gap-filling at least 16 reactions to allow *in silico* growth on PTYG (Fig. 3B). The number of added reactions further varied by strain and substrate, with flk requiring the highest overall number of additions (38 reactions), primarily for PTYG and succinate (Fig. 3B). Other strains, such as lur, required additions across multiple single substrates, reflecting substrate-specific gaps (Fig. 3B). The number of added gap-filling reactions varied both within and across each phylum, without any clear phylum-dependent tendency (Fig. 3B).

Curation and gap-filling resulted in model predictions closely matching observed growth phenotypes, with F1 scores^22^ and Matthews correlation coefficients (MCC)^23^ ≥ 0.8 for all strains, indicating good predictive accuracy (Table 2; Supplementary Fig. S3C-E).

**Table 2:** Genome properties and confusion-matrix metrics of draft and curated GEMs for the 21 soil SynCom isolates.

| Models | Class | Total Length (Mb) | Total Genes | Metabolic Genes | F1 Draft | F1 Final | MCC Draft | MCC Final |
| --- | --- | --- | --- | --- | --- | --- | --- | --- |
| bbet | Alphaproteobacteria | 8.39 | 7843 | 1463 | 0.429 | 1 | 0.302 | 1 |
| buol | Betaproteobacteria | 8.86 | 7864 | 1559 | 0.723 | 0.95 | -0.27 | 0.85 |
| cauj | Alphaproteobacteria | 5.69 | 5159 | 918 | 0.167 | 1 | 0.161 | 1 |
| cohm | Firmicutes | 8.01 | 6970 | 972 | 0.095 | 1 | 0.131 | 1 |
| cpi | Bacteroidetes | 7.63 | 5870 | 878 | 0.5 | 1 | 0.535 | 1 |
| cuvk | Actinobacteria | 4.09 | 3877 | 708 | 0.462 | 1 | 0.354 | 1 |
| cxy | Actinobacteria | 4.64 | 4265 | 714 | 0.167 | 0.8 | -0.024 | 0.8 |
| dri | Alphaproteobacteria | 4.47 | 4340 | 772 | 0.182 | 1 | 0.189 | 1 |
| flk | Bacteroidetes | 5.53 | 5455 | 689 | 0.483 | 0.93 | 0.23 | 0.91 |
| lgu | Gammaproteobacteria | 5.95 | 4793 | 745 | 0.5 | 0.89 | 0.144 | 0.85 |
| lur | Gammaproteobacteria | 4.88 | 4402 | 806 | 0.222 | 0.93 | 0.308 | 0.91 |
| mgk | Bacteroidetes | 5.92 | 5158 | 767 | 0.345 | 0.91 | 0.079 | 0.89 |
| mip | Actinobacteria | 3.45 | 3355 | 677 | 1 | 1 | 1 | 1 |
| mlo | Alphaproteobacteria | 6.5 | 6264 | 1244 | 0.414 | 0.93 | 0.066 | 0.91 |
| pmor | Gammaproteobacteria | 6.78 | 6044 | 1222 | 0.683 | 0.94 | 0.032 | 0.87 |
| pscf | Gammaproteobacteria | 6.2 | 5518 | 1064 | 0.667 | 0.94 | 0.059 | 0.87 |
| pzu | Alphaproteobacteria | 4.59 | 4463 | 834 | 0.182 | 0.8 | 0.023 | 0.8 |
| racd | Gammaproteobacteria | 5.37 | 4724 | 1049 | 0.667 | 0.93 | 0.167 | 0.87 |
| rfa | Actinobacteria | 5.39 | 5032 | 1120 | 0.261 | 1 | 0.236 | 1 |
| tro | Alphaproteobacteria | 5.93 | 5603 | 1149 | 0.174 | 1 | 0 | 1 |
| vpe | Betaproteobacteria | 7.06 | 6467 | 1229 | 0.486 | 0.9 | -0.321 | 0.8 |

### In silico minimal media analysis reveals strain-specific metabolic flexibility but auxotrophic limitations

Each curated GEM was subjected to in silico minimal-medium (iMM) analysis^24–26^ to identify alternative minimal nutrient sets that can support growth (Fig. 4; Supplementary Fig. S4). These sets were drawn from a defined pool of 69 metabolites representing PTYG, based on the medium recipe and published small-molecule compositions of yeast extract and trypticase^27^ (Supplementary Table S1; see Methods). The iMM analysis consistently identified O₂ and orthophosphate as essential in all solutions. Biotin and thiamine were also required in every alternative minimal-medium solution across strains, although iMM size varied by strain. (Fig. 4). Requirements for sulfate, nucleotides, amino acids, other cofactors, and other compounds varied among the strains, revealing species-specific signatures of nutritional dependence (Fig. 4). Strains differed in both the minimum number of nutrients required for modeled growth (iMM size) and the number of distinct nutrient combinations that could meet this minimum. For a given strain, these alternative sets contained the same number of nutrients but differed in composition. A larger iMM therefore indicated a requirement for more supplied nutrients, whereas more alternatives indicated greater flexibility in nutrient combinations within the PTYG-derived pool.

**Figure 4.**
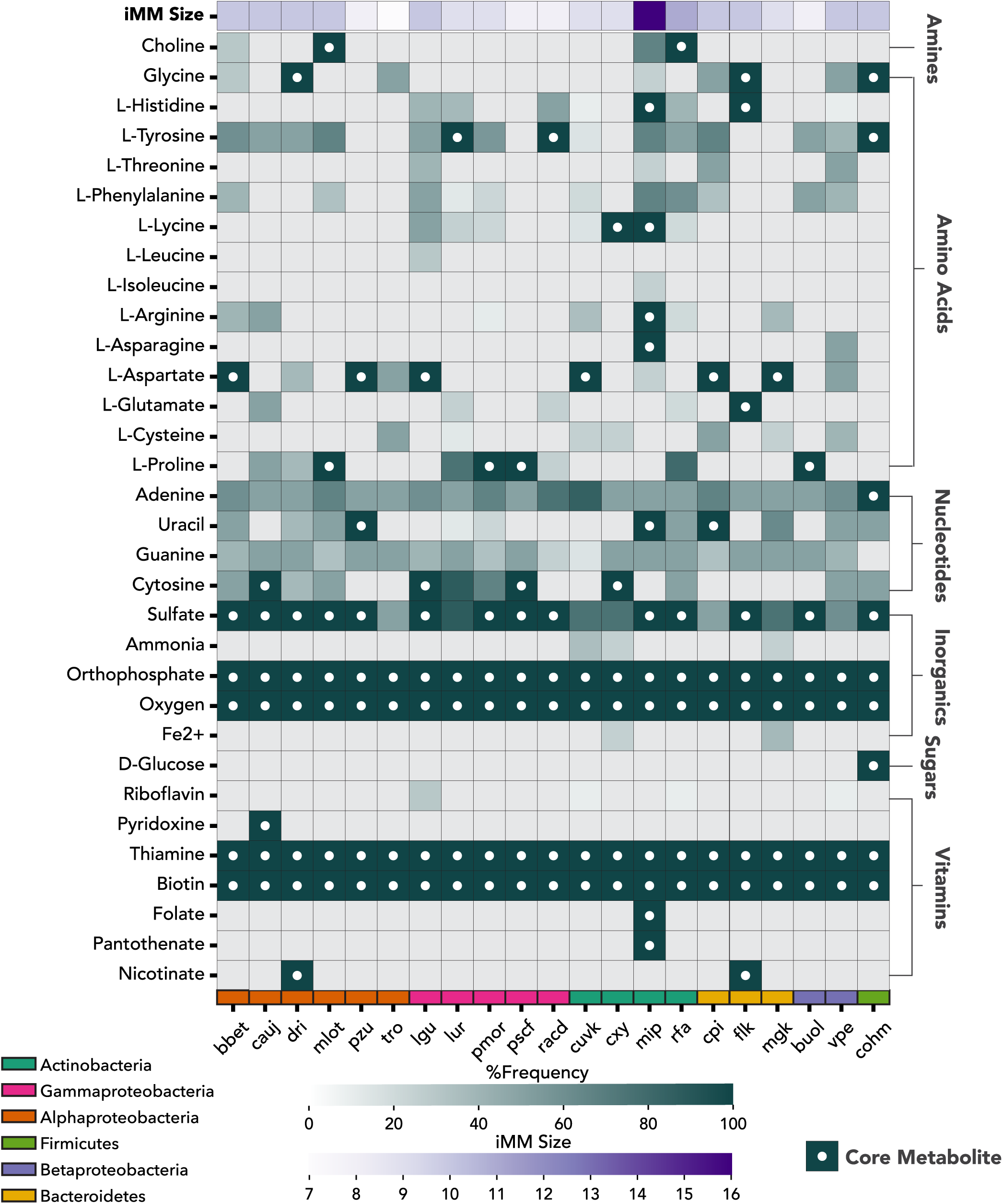
In silico minimal-media analysis identifies shared core requirements and strain-specific nutrient dependencies. The heatmap summarizes in silico minimal media (iMM) alternatives predicted for each SynCom strain. Columns represent strains grouped by phylum, and rows represent metabolites grouped by chemical class. Cell color indicates the frequency with which each metabolite appeared across the minimal-media alternatives for a given strain, from absent to present in all alternatives. White dots mark core metabolites required in 100% of alternatives. The top row shows iMM size, reflecting the number of media compounds required to support growth. Oxygen, orthophosphate, thiamine, and biotin were shared requirements across the community, whereas amino acids, nucleotides, sulfate, and other cofactors varied among strains. These patterns separate community-wide nutritional constraints from strain-specific auxotrophic requirements and metabolic flexibility.

The iMM analysis also helped pinpoint potential nutrient dependencies underlying the PTYG-restricted behavior of mip and cohm observed in the growth assays (Fig. 1). For mip, all 24 minimal iMM alternatives shared a fixed core consisting of four vitamins and cofactors (thiamine, biotin, folate and pantothenate), four amino acids (L-asparagine, L-arginine, L-lysine and L-histidine), uracil, sulfate, oxygen and orthophosphate. Additional compounds, including choline, glycine, adenine, guanine, L-aspartate, L-isoleucine, L-threonine, L-phenylalanine or L-tyrosine, appeared in subsets of alternatives, indicating flexibility in auxiliary nutrient acquisition. A smaller minimal medium was predicted for cohm: both minimal iMM alternatives contained D-glucose, glycine, L-tyrosine, adenine, sulfate, biotin, thiamine, oxygen, and orthophosphate, while the pyrimidine requirement could be met by either uracil or cytosine. Thus, both PTYG-restricted strains were predicted to depend on externally supplied biomass precursors, but mip required a broader amino-acid and cofactor core, whereas cohm required fewer compounds and showed only limited substitutability among nucleobases. In the initial growth assays, mip showed no measurable growth on 21C/M9 with succinate (Fig. 1, Supplementary Figure S1). To test its predicted nutrient requirements, we supplemented this medium with a mixture of the core nutrients shared by all 24 iMM alternatives. In the supplemented medium, mip showed an increase in OD₆₀₀ of 0.0280 ± 0.0034 after 150 h (mean ± SD; n = 4; Supplementary Fig. S5).

### Community context reshapes uptake-secretion profiles and niche overlap among strains

To capture potential metabolite exchange that may influence strain growth in a community context, we used Decomposed in silico Minimal Exchanges^28^ (DiMEs; see Methods). DiMEs enumerate, for each strain, minimal sets of metabolites that can be taken up or secreted while supporting growth under thermodynamic constraints. We defined the starting environment using water-extractable compounds detected by LC–MS in sterile soil (see Methods). After mapping these compounds to model metabolites, this pool included 145 exchangeable metabolites and served as the basal environment for DiMEs simulations (Supplementary Table S2). We then constructed a conditioned environment by adding the union of metabolites predicted to be secreted by SynCom strains in the basal environment and repeating this procedure until no additional secreted metabolites were identified.

We summarized the exchange profiles in basal and conditioned environments using four complementary metrics (Fig. 5A-D). At the strain level, strain fertility describes the number of unique substrates a strain can take up, whereas strain versatility describes the number of unique products a strain can secrete. At the metabolite-class level, substrate fertility describes how many strains can take up at least one metabolite from a given substrate class, while product versatility describes how many strains can secrete at least one metabolite from a given product class. Thus, strain versatility and strain fertility describe the exchange breadth of individual strains, whereas substrate fertility and product versatility describe how broadly each metabolite class is represented across the SynCom.

**Figure 5.**
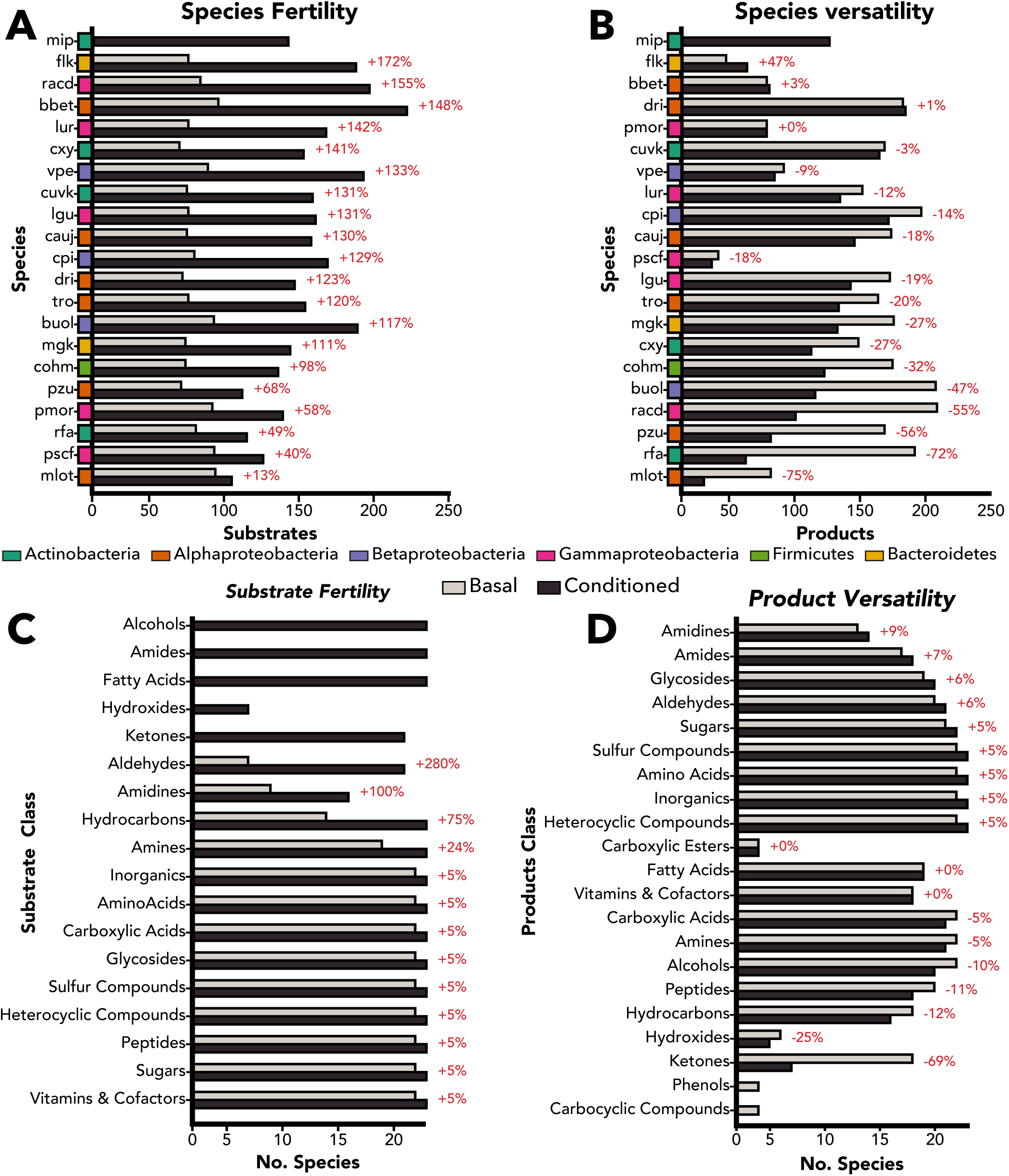
Community conditioning expands feasible uptake while reshaping secretion potential across the SynCom. Decomposed in silico Minimal Exchanges (DiMEs) were used to compare model-feasible exchange profiles in the basal environment and after iterative conditioning with predicted community secretions. (A) Strain fertility, defined as the number of unique feasible substrates per strain, increased after conditioning for all feasible strains, with mip becoming feasible only in the conditioned environment. (B) Strain versatility, defined as the number of unique feasible products per strain, changed more heterogeneously, with some strains gaining products and others losing secretion breadth. (C) Substrate fertility shows how many strains can access at least one compound from each substrate class. (D) Product versatility shows how many strains can secrete at least one compound from each product class. Percentages indicate change from basal to conditioned profiles; coloured side bars denote phylum.

Conditioning substantially expanded feasible resource access. Across all strains, the number of metabolites appearing as feasible substrates in at least one DiMEs solution increased from 94 in the basal environment to 243 after conditioning (Supplementary Fig. S6C, D). At the strain level, all strains that were growth-feasible in both environments showed greater predicted uptake breadth, although the magnitude of this expansion varied strongly (Fig. 5A). The largest gains were observed for bbet, racd, flk and vpe, which gained 126, 113, 112 and 104 additional feasible substrates, respectively. By contrast, mlot and pscf showed smaller increases, gaining 11 and 33 additional feasible substrates, respectively. Under the modeled basal environment, mip was infeasible but became feasible after conditioning, consistent with adding community-derived compounds to the simulated resource pool.

Secretion breadth followed a different pattern. Most strains had fewer feasible secreted products in the conditioned environment, whereas only a few strains, including flk, bbet, and dri, gained predicted products (Fig. 5B). For example, rfa decreased from 178 to 49 predicted products, racd from 195 to 87, and buol from 194 to 102, while flk increased from 34 to 50. pscf illustrates that uptake and secretion breadth capture different aspects of metabolic potential: it had one of the broadest feasible uptake profiles in the basal environment, but the narrowest predicted secretion profile under the minimal-exchange criterion. Thus, conditioning broadened feasible resource access, while byproduct secretion was constrained or redirected in a strain-dependent manner.

The predicted products were not all terminal byproducts. Of the 254 predicted basal secretions, 233 appeared as feasible substrates in the conditioned profiles, whereas 21 did not. Thus, most predicted products could in principle re-enter the community resource pool. At the same time, most predicted basal products were shared among multiple potential producers: 238 of 254 predicted secretions were associated with at least two producing strains, while 16 were unique to one predicted producer. These patterns indicate that the modeled conditioned environment comprised a mixture of broadly shared secretion capacities and a smaller set of more producer-specialized metabolites.

Because changes in resource access can alter potential competition^29^, we quantified pairwise nutritional niche overlap using the Niche Overlap Index (NOI) computed from uptake sets (Fig. 6A, B). NOI is directional: for a focal strain A, it measures how much of A’s feasible uptake set is also feasible for strain B. To compare the same strain set across environments, we evaluated 380 directed focal-strain comparisons among the 20 strains retained for the basal-conditioned comparison. Strong overlap (NOI ≥ 0.75) occurred in 343 comparisons in the basal environment but in only 192 comparisons after conditioning. Sixteen comparisons that initially showed very high overlap (NOI ≥ 0.9) fell below 0.5 after conditioning. This reduction is consistent with community-derived metabolites reducing predicted overlap in feasible resource use and shifting the community toward greater resource differentiation.

**Figure 6.**
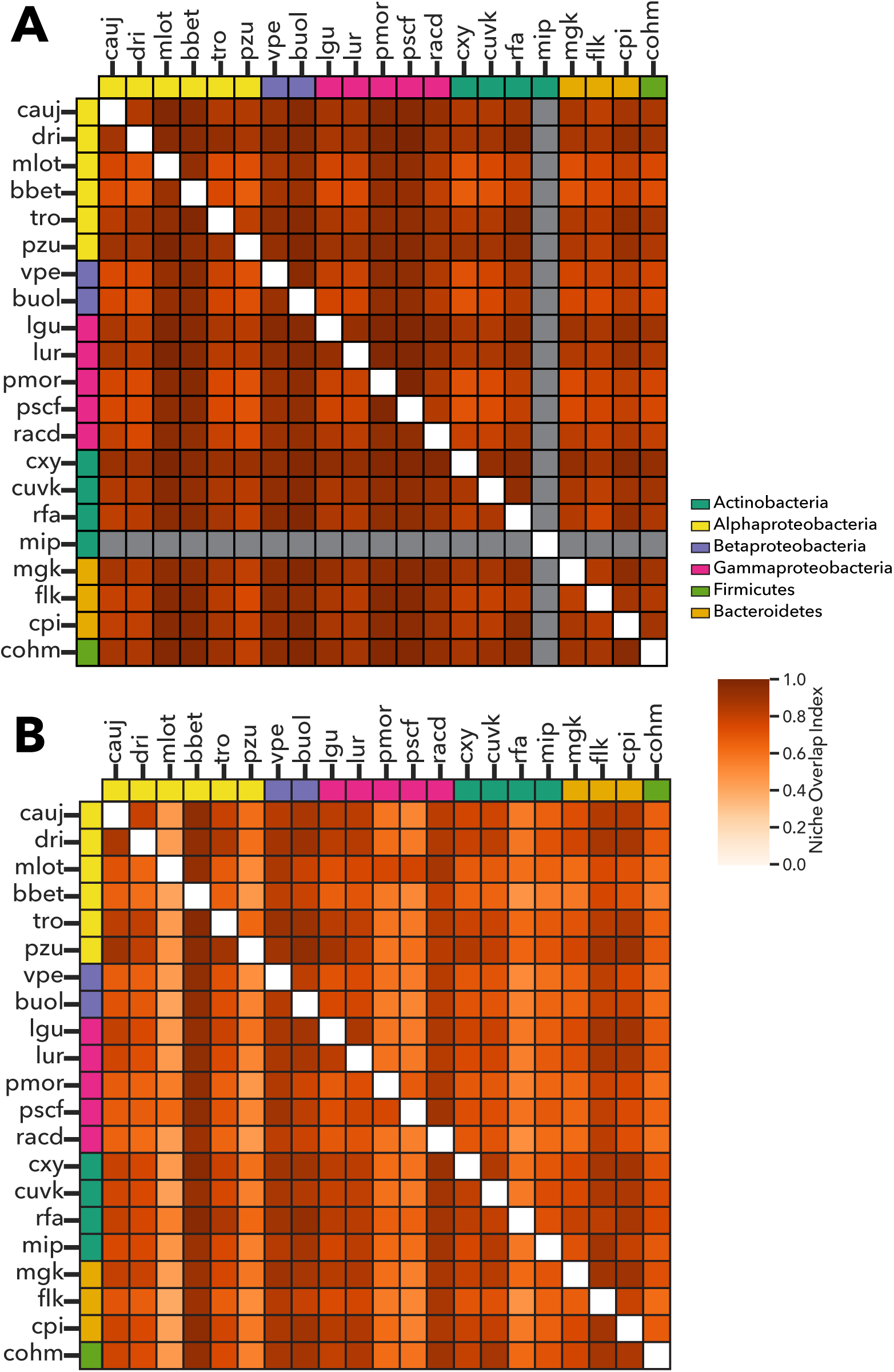
Community conditioning reduces predicted nutritional niche overlap across the SynCom. Niche Overlap Index (NOI) was computed from DiMEs-predicted feasible uptake sets. Rows represent focal strains, columns represent comparison strains, and each cell indicates the fraction of the focal strain’s feasible substrates that are also feasible for the comparison strain. (A) Basal environment. (B) Conditioned environment generated by iterative addition of predicted community secretions. Darker colors indicate higher overlap; white diagonal cells indicate self-comparisons, and grey cells correspond to comparisons involving mip, which was infeasible and therefore lacked a defined uptake set. Coloured bars denote phylum. In the basal environment, most defined strain pairs showed high overlap. After conditioning, overlap became more heterogeneous and lower for many pairs, with strong overlap (NOI ≥ 0.75) decreasing from 343/380 to 192/380 directed comparisons.

Finally, we visualized candidate producer-metabolite-consumer links induced by conditioning using class-resolved Sankey diagrams (Fig. 7). In each panel, strains on the left are predicted producers, central nodes are the newly secreted metabolites, and strains on the right are predicted consumers. Links denote metabolites that were predicted to be secreted in the basal environment and to become feasible uptake substrates only after conditioning (self-links were excluded). We show three representative classes in the main figure, sugars, carboxylic acids, and vitamins and cofactors, while additional classes are provided in Supplementary Figure S7.

**Figure 7.**
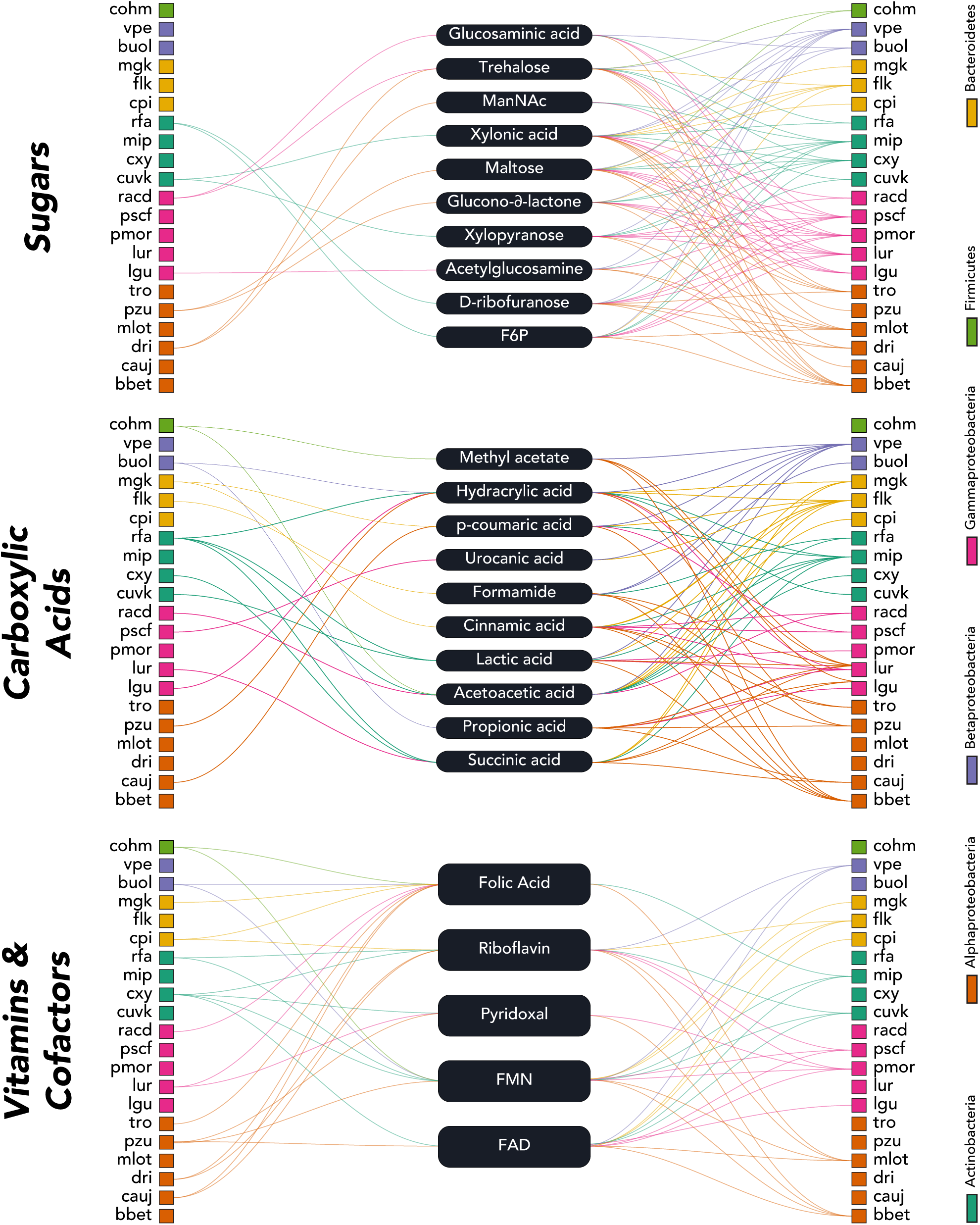
Community conditioning nominates candidate shared metabolites across chemical classes. Producer-metabolite-consumer maps show candidate exchange links inferred from DiMEs profiles after community conditioning. For each class, strains on the left are predicted producers, central nodes are metabolites, and strains on the right are predicted consumers. Links indicate metabolites predicted to be secreted in the basal environment and to become feasible uptake substrates after conditioning; self-links were excluded. Three representative classes are shown: sugars, carboxylic acids, and vitamins and cofactors. Several metabolites showed asymmetric connectivity, with one or a few predicted producers linked to many potential consumers, including xylonic acid, maltose, trehalose, propionic acid, succinic acid, flavin adenine dinucleotide (FAD), and folic acid. Strain colors indicate phylum. These maps nominate candidate community currencies and testable cross-feeding hypotheses, rather than directly measured metabolite transfer.

The representative maps reveal that a subset of metabolites had an asymmetric producer-consumer structure, with few predicted producers and many predicted consumers. Among sugars, xylonic acid connected one predicted producer, cuvk, to 16 predicted consumers, while maltose connected one predicted producer, pzu, to 12 predicted consumers (Fig. 7). Trehalose showed a similar pattern, with two predicted producers, dri and racd, and 10 predicted consumers. Among carboxylic acids, propionic acid connected one predicted producer, buol, to 14 predicted consumers, while succinic acid connected three predicted producers to six predicted consumers (Fig. 7). Vitamins and cofactors showed both asymmetric and more distributed patterns: flavin adenine dinucleotide connected two predicted producers to 10 predicted consumers, whereas folic acid had a broader producer set and fewer predicted consumers (Fig. 7). Together, these maps nominate candidate shared metabolites that may act as community currencies under the modeled constraints. These maps therefore provide testable hypotheses rather than direct evidence of realized exchange.

### Biosynthetic network similarity reveals functional clusters independent of phylogeny

We used the NIS^30^ biosynthesis module to compare strains by the reaction subnetworks that synthesize biomass building blocks. We focused the main network analysis on 21C/M9 minimal medium with succinate, because this medium provides a defined nutritional context for comparing biosynthetic routes across strains. The analysis included 19 strains for which the comparison was feasible under this condition. For each biomass-building-block class, amino acids, lipids, cofactors, and nucleotides, we calculated pairwise Jaccard similarity from the reactions used in the predicted biosynthetic routes. We also repeated the analysis under PTYG and show the corresponding clustered similarity heatmaps in Supplementary Fig. S8.

We visualized the 21C/M9 similarities as thresholded networks (Fig. 8). In these networks, each node is a strain, and each edge connects two strains whose biosynthetic routes are similar above a given Jaccard threshold. At low thresholds, many weak similarities are shown. At higher thresholds, only the strongest similarities remain. Therefore, networks that retain many edges at stringent thresholds indicate more conserved biosynthetic routes, whereas networks that lose edges quickly indicate more divergent routes.

**Figure 8.**
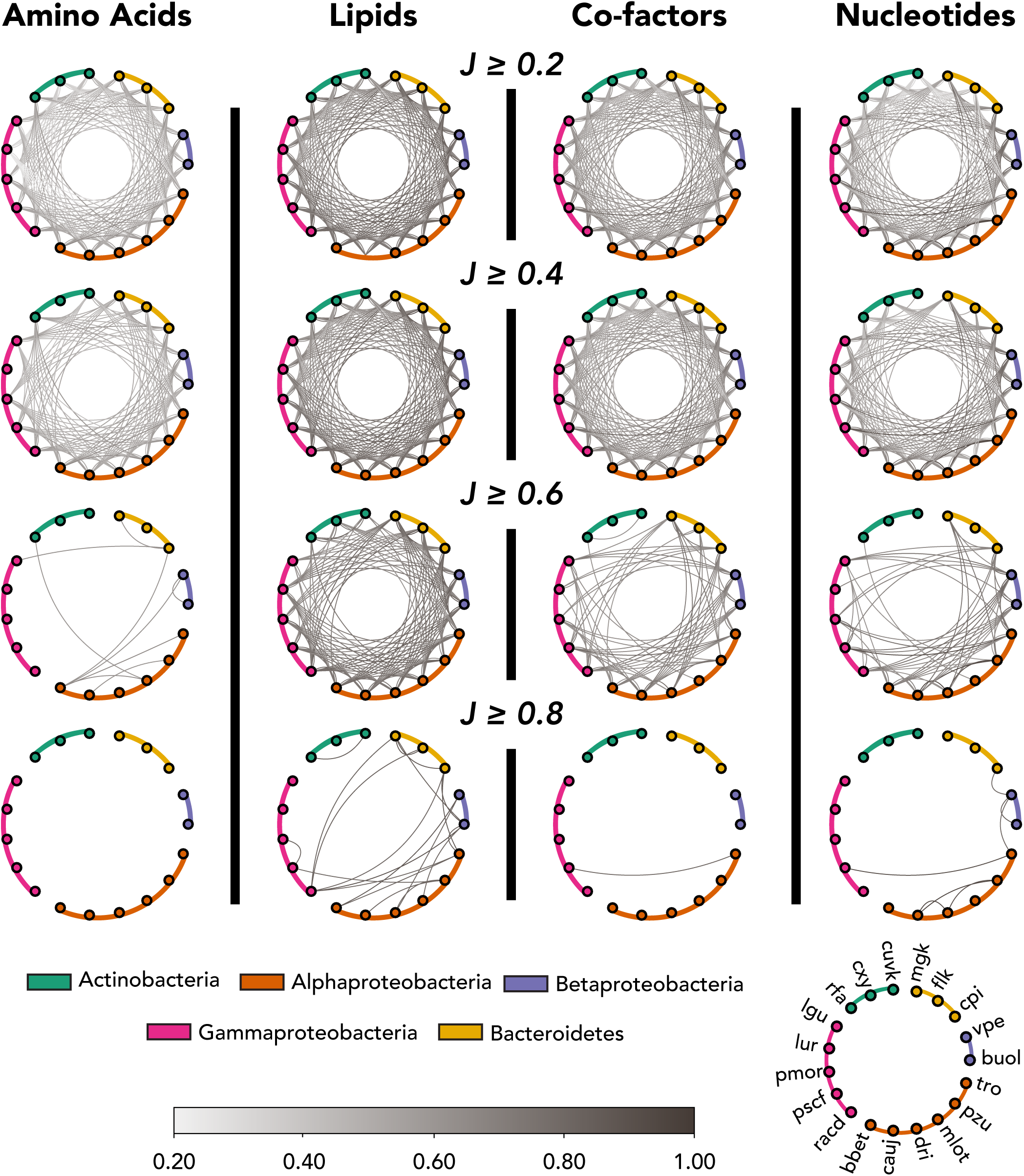
Biosynthetic similarity is pathway-specific across SynCom strains. NIS was used to compare biosynthetic reaction subnetworks for biomass-building-block classes under 21C/M9 minimal medium with succinate. Each Column shows one biosynthetic class, amino acids, lipids, cofactors or nucleotides, and each row applies an increasing Jaccard similarity threshold (J ≥ 0.2, 0.4, 0.6 or 0.8). Nodes represent the 19 strains feasible in this condition, colored by phylum. Edges connect strain pairs with biosynthetic route similarity above the indicated threshold; darker edges indicate higher similarity. Amino-acid biosynthesis lost most connections at stringent thresholds, indicating divergent routes among strains. Lipid biosynthesis retained many high-similarity links, whereas cofactor and nucleotide biosynthesis showed intermediate structure. These patterns show that functional similarity depends on the biosynthetic pathway considered rather than taxonomy alone.

This comparison showed that biosynthetic similarity was pathway-dependent. Amino-acid biosynthesis was the most divergent: although all 171 strain pairs were connected at J ≥ 0.2, only 9 pairs remained connected at J ≥ 0.6, and none remained connected at J ≥ 0.8. This means that few strain pairs shared highly similar amino acid biosynthetic routes under stringent similarity cutoffs.

Lipid biosynthesis showed the opposite pattern. All strain pairs remained connected at J ≥ 0.4, 138 pairs remained connected at J ≥ 0.6, and 20 pairs remained connected at J ≥ 0.8. This indicates that lipid-building routes were more conserved across the SynCom than amino-acid biosynthesis. Cofactor and nucleotide biosynthesis showed intermediate structure: cofactors retained 75 edges at J ≥ 0.6 but only one edge at J ≥ 0.8, whereas nucleotides retained 60 edges at J ≥ 0.6 and 7 edges at J ≥ 0.8.

The PTYG heatmaps provided a complementary medium-context comparison (Supplementary Fig. S8). In this rich-medium setting, lipid biosynthesis again showed high similarity across many strain pairs, with 192 of 210 pairs retained at J ≥ 0.6 and 70 retained at J ≥ 0.8. Cofactor and nucleotide biosynthesis showed intermediate structure, retaining 68 and 64 pairs at J ≥ 0.6, respectively. By contrast, the amino-acid matrix showed no pairwise similarity signal in the PTYG output, consistent with the rich medium changing the biosynthetic routes NIS recovered for this class. Thus, the PTYG analysis is best interpreted as a medium-context check rather than a direct repetition of the 21C/M9 network pattern.

These similarity patterns were not uniformly explained by taxonomy. Some high-similarity pairs occurred within the same class, such as flk and mgk for lipid biosynthesis or cauj and pzu for nucleotide biosynthesis. However, several strong similarities also connected strains from different phyla, including buol and racd for lipid biosynthesis, and pscf and tro for nucleotide and cofactor biosynthesis. In addition, the same strain pair could be highly similar in one biosynthetic class but much less similar in another. For example, buol and cpi were highly similar in lipid biosynthesis but only moderately similar in amino-acid biosynthesis. Thus, functional proximity depended on the biosynthetic pathway considered rather than taxonomy alone.

Because the NIS comparison suggested that biosynthetic similarity depends on pathway and medium context, we next asked whether predicted metabolic bottlenecks also depended on the nutritional environment. Gene essentiality analysis supported this context-dependent view of biosynthetic function (Supplementary Fig. S9B & S10). Under 21C/M9 minimal medium with succinate, the 19 feasible strains had 67–96 unique essential genes per strain (median, 85), whereas PTYG supported smaller essentiality sets across all 21 strains, with 48–65 unique essential genes per strain (median, 58). Cofactor-related bottlenecks were the largest class in both media, but amino-acid-related essentiality was prominent under 21C/M9 with succinate and not recovered in the PTYG-classified output. Thus, predicted metabolic bottlenecks depended on both strain identity and medium composition, supporting the view that biosynthetic function is context dependent.

## Discussion

Microbial communities assemble and persist through interactions between species-level traits and a chemically dynamic environment^31,32^. In soil, this coupling is especially strong because resource availability is heterogeneous, temporally variable, and shaped by the community itself^33^. Here, we used a defined soil SynCom to build a strain-level, mechanistic account of how biosynthetic capacity and metabolite exchange jointly structure ecological opportunity. By combining curated metabolic reconstructions with community metabolomics, we link what strains can make, what they can access, and how those possibilities change as the community conditions the environment.

A central message is that nutrient dependence is context-dependent rather than intrinsic. Minimal-media analyses reveal that strains differ not only in what they require, but in how many alternative nutrient solutions can satisfy those requirements. Some nutrients formed an invariant core, whereas others could substitute for one another across alternative solutions, indicating flexibility in acquisition routes. This distinction matters ecologically: core requirements define hard constraints that restrict where a strain can establish, while optional requirements define degrees of freedom that multiple environmental configurations can satisfy. In soils, where nutrient availability fluctuates and local microenvironments differ, such flexibility likely shapes both persistence and competitive outcomes^34,35^.

Community conditioning extends this principle by changing the resource landscape itself. In the DiMEs analysis, conditioning expanded feasible uptake breadth across strains, indicating that community-derived metabolites broadened the set of resources that strains could potentially access. Secretion breadth followed a different pattern: most strains had fewer feasible secreted products after conditioning, while a smaller subset gained products. This asymmetry suggests that community metabolism does not simply enrich the environment. Instead, it expands access to resources while redirecting the byproducts predicted under basal-exchange constraints. In ecological terms, the community reshapes the mapping between metabolic traits and feasible niches.

This chemical reshaping altered predicted niche relationships. Conditioning reduced pairwise nutritional niche overlap among the strains represented in both NOI matrices, consistent with lower overlap in feasible resource use after adding community-derived metabolites. This pattern supports the interpretation that conditioning can open more differentiated resource opportunities, although it should not be read as direct evidence of realized resource partitioning. The result instead shows that the potential competitive landscape depends on the chemical context that constrains each strain’s feasible uptake set. These patterns align with a view of microbial coexistence in which competition is not fixed by taxonomy or baseline physiology but varies with the metabolite landscape that the community collectively produces^36^.

Integrating LC-MS-derived chemical information with DiMEs^28^ predictions provided a way to move from environmental chemistry to strain-level exchange hypotheses. We defined the modeled basal environment from LC-MS-detected water-extractable compounds in sterile soil, after mapping these compounds to exchangeable metabolites in the GEMs. DiMEs then enumerated feasible uptake and secretion profiles for each strain under thermodynamic constraints and used predicted secretions to construct a conditioned environment. This design separates the chemical context used to constrain the simulations from the model-inferred strain-level roles. In turn, the resulting producer-consumer assignments can indicate which strains may contribute to the uptake or secretion patterns detected by LC-MS.

The candidate producer-consumer maps highlight a smaller subset of metabolites with asymmetric connectivity. These metabolites were predicted to be secreted by one or a few strains but to become feasible substrates for many others after conditioning. This structure nominates candidate community currencies, in the limited sense that production by a restricted set of strains could broaden resource access for a larger set of predicted consumers. Such metabolites are useful hypotheses for targeted testing because they connect network structure to experimentally tractable perturbations: adding or withholding them should alter feasible consumer sets if the predicted links are biologically relevant.

The pathway-level biosynthesis comparisons add a second layer to this trait-environment view. Under 21C/M9 minimal medium with succinate, amino-acid biosynthetic routes were comparatively divergent, lipid routes were more conserved, and cofactor and nucleotide routes showed intermediate structure. PTYG provided a complementary medium context, with lipid biosynthesis again showing high similarity, but amino-acid route similarity was not recovered in the PTYG output. These patterns indicate that functional similarity is not a single organism-level property. It depends on both the biosynthetic pathway considered and the nutritional environment used to constrain the analysis. Gene essentiality supported the same conclusion: predicted essential-gene sets were larger under 21C/M9 with succinate than under PTYG, and bottleneck classes shifted with medium composition. Together, these results show that metabolic fragilities are strain- and context-dependent, rather than uniformly explained by taxonomy.

Together, these results support a general ecological principle: communities partly construct their own niche space. By releasing metabolites that become available to others, community members can change both resource accessibility and predicted overlap in resource use. In the SynCom, conditioning expanded feasible uptake opportunities while reducing predicted nutritional niche overlap, suggesting that community-generated chemistry can simultaneously increase connectivity and differentiate resource use. Which effect dominates will likely depend on the balance between metabolites that diversify accessible resources and metabolites that create shared resource axes. This trade-off provides a mechanistic lens for understanding why microbiome interventions can be context dependent: an introduced strain may fail not because it lacks genes for a focal function, but because it requires specific precursors, depends on a particular conditioned state, or enters a resource landscape already reshaped by resident strains.

Several directions would strengthen the link from feasible exchange to realized exchange. The interaction maps here describe metabolic feasibility under model constraints, not direct strain-resolved fluxes in the mixed community. The modeled basal environment also represents the detected and mapped compound set, rather than the full chemical complexity of soil or soil extract. Stable-isotope tracing, temporal metabolomics and perturbations that selectively add or remove candidate currency metabolites would help test whether the predicted links operate in mixed culture and whether specific metabolites modulate niche overlap. Extending the framework to spatial structure will also be important for soils, where diffusion limitation and microscale gradients can localize exchange and alter which metabolites become effectively shared^37^.

In conclusion, our study connects species-level metabolic traits to community-level ecological organization in a defined soil microbiome. By showing how conditioning changes feasible exchange potential, reduces predicted niche overlap, and reveals pathway-specific functional similarity not fully explained by taxonomy, we provide a mechanistic basis for asking when strains compete, when they complement one another, and which metabolites may shape community connectivity. This trait-environment perspective offers a route to more predictive soil microbiome ecology and more rational strategies for testing how defined communities respond to nutrient and metabolite perturbations.

## Methods

### Strains, genomes and annotations

The 21 strains comprising the soil SynCom have been described previously^20^ and are listed in Table 1, Supplementary Table S3 and Supplementary Fig. S11. For genome sequencing, isolates were cultured individually in liquid PTYG medium, and Genomic DNA was extracted as described previously^20^. Sequencing was performed at the Lausanne Genomic Technologies Facility. We sequenced five isolates using both Illumina and Oxford Nanopore Technologies (ONT): vpe (004_S2), flk (005_S3), mip (012_S5), mgk (015_S7), and cuvk (018_S8). Illumina sequencing generated 150-bp paired-end reads, and ONT sequencing generated long reads. For the remaining 16 isolates, the genome assemblies used in this study were generated from Pacific Biosciences (PacBio) long reads and were assembled using the Microbial Assembly application in SMRT Link v9.0.0.92188.

### Read processing and hybrid assembly

Illumina reads were processed with Trimmomatic v0.39^38^ to remove adapters using mismatch, palindrome, and simple clipping thresholds of 3, 25, and 6, respectively. Low-quality leading and trailing bases were removed using LEADING:9 and TRAILING:9, and reads were trimmed using a four-base sliding window with a minimum mean quality of 15 (SLIDINGWINDOW:4:15). Reads shorter than 60 bases after trimming were discarded (MINLEN:60). Read quality before and after processing was assessed with FastQC v0.11.7^39^.

ONT reads were filtered with Filtlong v0.2.0 using a minimum read length of 7,000 bases (-- min_length 7000), a length weight of 10 (--length_weight 10) and a minimum mean quality of 10 (--min_mean_q 10). The target number of retained bases was set to approximately 100 times the expected assembly length (--target_bases). Filtered reads were assembled de novo with Flye v2.7.1^40^, using the --nano-raw option, eight threads and five internal polishing iterations, with an estimated genome size supplied for each input. Assembly graphs were inspected with Bandage^41^.

During the principal assembly-polishing stage, assemblies underwent two rounds of polishing with ONT reads, using GraphMap v0.5.2^42^ for mapping and Racon v1.0.1^43^ for polishing. These were followed by three rounds of polishing with trimmed Illumina reads, using Bowtie2 v2.3.4.1^44^ with the --local --very-sensitive-local options for mapping and Pilon v1.22^45^ for polishing. SAMtools v1.8 was used for BAM-file handling during these polishing steps. Assembly quality was assessed after each polishing round with QUAST v4.6.0^46^. For ONT inputs containing reads from two isolates, contigs were assigned to individual isolates using isolate-specific Illumina read mappings. Sequencing depth was estimated by mapping reads to the final assemblies with BWA v0.7.17^47^ and calculating coverage with SAMtools v1.11^48^.

### Annotation and taxonomic assignment

All assemblies were annotated with Prokka v1.1.3^49^ using GenBank/ENA/DDBJ-compliant output (--compliant) and an e-value cutoff of 0.001 (--evalue 0.001). Annotated assemblies were then reoriented to begin at *dnaA*. Taxonomic classification of the genome assemblies was performed with GTDB-Tk v2.7.2 (classify_wf) against GTDB release R232^50,51^. The genome sequences have been submitted to ENA and are retrievable under accession number PRJEB124582.

### In vitro media compositions

PTYG medium contained 0.5 g L⁻¹ glucose, 0.5 g L⁻¹ yeast extract, 0.25 g L⁻¹ peptone, 0.25 g L⁻¹ trypticase, 0.6 g L⁻¹ MgSO₄·7H₂O, and 0.07 g L⁻¹ CaCl₂·2H₂O. The small-molecule composition of yeast extract and trypticase was taken from the recent characterisation by Marinos et al. (2020)^27^, which lists 62 metabolites spanning inorganics, amino acids, vitamins and sugars^27^. For in silico use, we mapped PTYG components to 69 metabolites in the models (Supplementary Table S1). The 21C/M9 minimal medium followed Jones^52^: Hutner’s mineral base, 8.6 mM NaCl, 47.8 mM Na₂HPO₄, 22.1 mM K₂HPO₄, 18.7 mM NH₄Cl, and a vitamin solution (2 mL L⁻¹) containing biotin (0.5 mg L⁻¹), nicotinic acid (50 mg L⁻¹), and thiamine (25 mg L⁻¹). Succinate (10 mM) was supplied as the sole carbon source.

### Growth phenotype assays

Each strain was grown in four biological replicates in 200 µl liquid 21C minimal medium with 10 mM succinate or in PTYG, filled in a sterile 96-well plate. Cultures were incubated at room temperature with continuous shaking in a BioTek Synergy H1 plate reader, and the increase in culture turbidity at 600 nm was recorded during 72 h. These data provided the phenotype reference for reconstruction, curation, and subsequent gap-filling decisions.

Raw OD₆₀₀ time series were background-corrected against medium controls. A binary growth call per well was made using a pre-specified threshold on either (i) ΔOD (max OD - initial OD) or (ii) a fitted logistic/sigmoid parameter (e.g., asymptote or growth rate). Replicates were aggregated by majority vote to a single call per strain-medium combination; ties/ambiguous wells were flagged and excluded from scoring.

In addition, substrate utilization was profiled using BIOLOG plates. Each strain was inoculated individually in biological triplicates on the prefilled 30 different carbon substrates provided by BIOLOG and again incubated in a BioTek plate reader. Growth was then assessed from conversion of the resazurin salt detected at 590 nm.

### Metabolomics

For the community metabolomics assay we used a defined basal medium containing a fixed panel of supplied metabolites (composition in Supplementary Table S2). This medium was used only for the metabolomics experiment. Because the panel is fixed, observations are limited to those compounds: depletion relative to the starting concentration indicates net uptake, and accumulation indicates net secretion over the assay window.

### GEM reconstruction and curation

Draft genome-scale metabolic models (GEMs) were reconstructed for all SynCom members from their Prokka annotations^49^ using RAVEN v2.0^14^ and the KEGG 2022 prokaryotic reaction set^16^. We used a stringent homology threshold (E-value = 1×10⁻⁵⁰) to minimize spurious assignments and a more relaxed threshold (E-value = 1×10⁻^10^) to support the gap-filling process. Unbalanced and incomplete reactions were removed from the draft reconstructions. Transport reactions following the phosphotransferase system (PTS) mechanism were curated to include the phosphoenolpyruvate to pyruvate transformation. Each GEM was structured with two cellular compartments, cytoplasm and periplasm, and the extracellular space.

The simulated extracellular space of each model includes curated metabolites from the extracellular space of the *E. coli* iML1515 GEM^53^, all PTYG constituents, succinate and salts for 21C/M9, the 29 single-carbon substrates used in BioLog growth assays, and compounds detected in community metabolomics. Transport mechanisms were adapted from the *P. veronii* iPsvr GEM^54^.

The biomass reaction was adapted from the iPsvr GEM by substituting lipoyl-ACP with octanoyl-ACP, and vaccenic acid with linolenic acid. Additionally, we curated reaction directionalities using thermodynamic information to prevent CO₂ fixation and growth on CO₂ alone in non-autotrophs, avoid oxygen generation, and ensure that non-growth-associated ATP requirements could not be met without a carbon source.

Draft reconstructions lacked the reactions required to synthesize all BBBs and to reproduce observed growth on PTYG and 21C/M9 with succinate. To add essential reactions, we systematically gap-filled the draft GEMs using the NICEgame workflow^21^ and the prokaryotic subset KEGG 2022 as a reaction database. NICEgame identifies alternative minimal reaction sets that enable the production of each of the BBBs that cannot be produced prior to gap-filling. Gap-filling proceeded in a media-aware manner: first on PTYG, then on 21C/M9 with succinate, and finally on single-substrate conditions from the BIOLOG growth tests, if needed.

To prioritize biologically plausible solutions, we introduced an evidence-filtering step. First, we generated draft reconstructions of each GEM but this time with a less stringent homology similarity threshold (E-value = 1×10⁻¹⁰) and preferentially retained gap-fill alternatives containing reactions for which the respective genome encoded genes under the relaxed model. Second, remaining alternatives were scored using an EC-evidence rubric (Table 3), gathered from NCBI, BioCyc, UniProt, and BRENDA^16,55–58^. The highest-scoring minimal alternative(s) per BBB were then integrated (Supplementary Figure S12).

**Table 3:** Evidence rubric used to rank NICEgame gap-filling alternatives. Candidate minimal reaction sets proposed by NICEgame to restore production of missing biomass building blocks were scored based on the strength of external evidence supporting the enzyme commission (EC) number associated with reactions in the alternative. Full EC matches receive higher scores when supported in the target species, a closely related species, or the genus (scores 5–3), whereas partial EC matches (first three digits) receive lower scores (2–1). A +1 bonus is added when the corresponding gene is also present in a draft model reconstructed under a less stringent homology threshold, and the highest-scoring alternative(s) were integrated during gap-filling.

| Evidence type | Score |
| --- | --- |
| E.C. exists in species | 5 |
| E.C. exists in a closely related species | 4 |
| E.C. exists in the genus | 3 |
| 3 letters E.C. exist in the species/closely related species | 2 |
| 3 letters of E.C. exist in the genus | 1 |
| The gene exists in the less stringent homology similarity threshold model | + 1 |

The models were thermodynamically curated by incorporating estimated Gibbs free energies of formation and their associated uncertainties^59^ (Supplementary Fig. S13B, C). Maximal substrate uptake rates were estimated by constraining the models to experimentally measured growth rates using thermodynamics-based flux balance analysis^60^.

### Model evaluation

At each stage of the reconstruction process, we simulated models on the same media and compared growth/no-growth predictions to experimental data. From the resulting confusion matrix, we computed the F1 scores, the Matthews Correlation Coefficients (MCC), accuracy, and precision (Table 2; Supplementary Fig. S3C–E). We additionally benchmarked models with MEMOTE^61^, which evaluates annotation completeness, basic consistency, biomass reaction properties, and stoichiometric balance. The SynCom models achieved an overall score ≈ 81% (Supplementary Figure S13A), close to well-curated references (e.g., iML1515), indicating strong structural quality.

### In silico minimal media (iMM) analysis

We identified the smallest nutrient sets that enable growth for each GEM using the in silico minimal media (iMM) analysis workflow that casts nutrient selection as a cardinality minimization problem and enumerates all minimal alternatives^24–26^. We ran iMM separately on (a) the PTYG-derived pool of components (69 mapped small molecules spanning inorganics, vitamins, amino acids, and sugars) and (b) a 21C/M9 pool (minerals/vitamins plus succinate) to assess medium dependence. For each strain, we reported (i) the number of minimal alternatives, and (ii) the fractional occurrence of each metabolite across alternatives (0-100%). Metabolites with 100% occurrence are denoted essential. Metabolites with intermediate occurrence are optional and reflect flexibility in acquisition routes.

### In silico identification and analysis of uptake–secretion profiles

We characterized the feasible uptake and secretion profiles of each GEM using the DiMEs (Decomposed in silico Minimal Exchanges) method^28^. Metabolomics data from the water-extractable compounds in sterile soil guided the simulation of metabolites available in the extracellular space, defining a basal medium of 145 metabolites (Supplementary Table S2). For each GEM, we ran DiMEs under basal configuration to identify secreted metabolites that can appear in any minimal exchange solution achieving experimentally observed growth rates. We then augmented the medium with the union of these secretions across species and repeated the procedure until we found no new secretions. The resulting augmented pool defines the conditioned environment used throughout the paper.

Subsequently, for each species and medium (basal or conditioned), we computed DiMEs solutions/profiles at optimal and suboptimal biomass yields (100%, 90%, …, 10% of the single-species optimum; Supplementary Fig. S14). Formally, this consists of solving a mixed-integer problem that minimizes the number of active exchange reactions (uptakes and secretions) required to achieve a target growth rate, under steady-state and TFA constraints.

Finally, for each species, the union of minimal solutions was collapsed into profile sets of (a) metabolites that can be taken up and (b) metabolites that can be secreted. Profiles were reported at the metabolite-class level (curated via MeSH^62^, ChEBI^63^, and PubChem^64^) as class coverage (Supplementary Fig. S15):

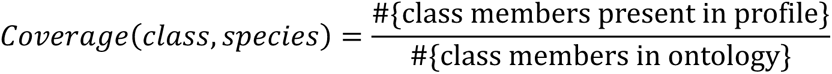

Species were ordered by “versatility” (breadth across classes; mean coverage), while classes were ordered by “fertility” (fraction of species for which the class is present).

Using the DiMEs results, we quantified the pairwise nutritional niche overlap^29^ for any focal species A and a competitor strain B:

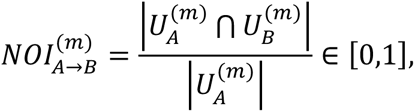

where *U_s_^(m)^* is the set of metabolites that species *s* can uptake in medium *m* (we computed *m* ∈{basal, conditioned} from our in-silico uptake profiles). Values near 1 indicate that most of species *A*’s usable resources are also usable by species *B*. Strain-medium combinations with no uptake set ∣*U_s_^(m)^=0*∣) are undefined and plotted as NA/grey in Figure 6 (e.g., basal non-growers).

### Comparison of the biosynthetic capabilities of the members of the soil SynCom

We used the NIS^30^ biosynthesis module to compare strains by the biosynthetic routes that produce each biomass-building block (BBB). For every strain and BBB in the biomass reaction, NIS enumerated minimal, elementally balanced subnetworks that can synthesize the BBB from a defined core network under our modeling constraints (Supplementary Fig. S16). For each strain, we took the union of reactions across its alternative route for a given BBB and computed pairwise pathway similarity between strains with the Jaccard index (J)^65^ on reaction IDs. We then averaged J within each BBB class (amino acids, nucleotides, cofactors, lipids) to obtain one similarity matrix per class (Supplementary Figs. S9A and S8). This matrix was hierarchically clustered on 1−J with average linkage. All runs used the curated GEMs with TFA constraints under a common minimal medium (21C/M9 + succinate), or with PTYG medium.

### Gene and reaction essentiality

We performed gene essentiality analysis under thermodynamic constraints on PTYG and 21C/M9 with succinate^60,66^. A gene was considered essential if the mutant growth rate was below 10% of the growth rate of the wild type. For each essential deletion, we attributed its effect to the BBB class whose production was impaired (amino acids, nucleotides, lipids, cofactors, etc.).

### Identifier harmonisation and annotation sources

Reactions, metabolites, and genes identifiers were mapped across databases, including KEGG^16^, ModelSEED^67^, BiGG^68^, MetaNetX^69^, and NCBI^58^, so that every entity has (i) a primary namespace and (ii) traceable cross-references. KEGG compound IDs and reaction IDs served as the primary metabolite and reaction identifiers throughout, with MetaNetX used to reconcile synonyms and legacy IDs and to populate cross-reference tables. For each metabolite, we compiled: name, formula, charge, compartment, KEGG ID, InChIKey and SMILES (when available). Structures were sourced from KEGG and verified/filled via ChEBI and PubChem. Gene identifiers were anchored to the Prokka locus tags (genome-native) and mapped to NCBI accessions for provenance.

Metabolites were assigned to high-level classes (e.g., amino acids, fatty acids, nucleotides, cofactors, inorganics, phenols, polyamines, carboxylic esters) using a curated merge of MeSH^62^ headings and ChEBI^63^ ontology. These labels underpin the “class coverage” and “class fertility” statistics reported for DiMEs profiles.

### Calculation of metabolite Gibbs free energies of formation

The Gibbs free energy of formation is a fundamental thermodynamic property used to predict the directionality of chemical reactions. In general, a reaction will operate in the direction where the Gibbs free energy of formation of the products is lower than that of the reactants^59^. To calculate the Gibbs free energy of formation for the compounds, we first used Marvin (version 18.1, 2018, ChemAxon http://www.chemaxon.com) to transform the compound structures (canonical SMILES) into their major protonation states at pH=7 and generate their MDL molfiles (a file format used to store molecular structures and associated data). SMILES is a chemical notation that uses a string of characters to represent the compound’s molecular structure in machine-readable form^70^. The standard Gibbs free energy of formation, along with the estimation error, was then estimated using the Group Contribution Method (GCM). The group contribution method is a technique used in thermodynamics to predict the properties of complex chemical compounds based on the thermodynamic properties of their constituent functional groups. The contribution of each functional group to the molecule’s overall Gibbs free energy is then determined based on its chemical structure and bonding. Once the contribution of each functional group to the Gibbs free energy has been determined, the overall Gibbs free energy of the molecule can then be calculated by summing the contributions of all the functional groups^71^.

### Software, solvers, versions, and computing environment

All analyses were performed on Ubuntu 22.04.x Linux workstations. Reconstruction, curation, and simulations ran in MATLAB R2020b with COBRA Toolbox 3.0^72^ (general FBA utilities, model I/O), RAVEN 2.0^14^ (draft reconstruction from KEGG Orthology; task checks, model quality control), matTFA^60^ (thermodynamics-based flux analysis and variability analysis), phenomapping^66^ (thermo-aware single gene/reaction deletions). We used NICEgame^21^ for gap-filling with the KEGG 2022 prokaryotic reaction pool. We used MEMOTE^61^ for independent model quality control (annotation, basic tests, biomass, stoichiometric consistency). Summary plots and heat maps were produced in Python (data handling/plotting), with export of publication SVG/PDFs. Linear and mixed-integer programs were solved using GUROBI 9.5.2 or IBM ILOG CPLEX 12.10. Unless stated otherwise, we used default tolerances (optimality gap ≤ 1e-6 for MILPs) and reported an infeasible solve as zero growth^16,61,73,74^.

## Author Contributions

J.R.v.d.M. and V.H. conceived the study. O.K., E.V., J.R.v.d.M. and V.H. developed the methodology. O.K. reconstructed and curated the metabolic models, developed the software, curated the data, performed the formal analyses, and prepared the visualizations. S.Č. conducted the laboratory experiments. D.S.W. processed the genome sequencing data and annotated the genomes. A.S. managed sequence data handling and submission. E.V. provided direct supervision, and J.R.v.d.M. and V.H. supervised the project. O.K. wrote the original manuscript draft, and E.V., J.R.v.d.M. and V.H. reviewed the work and the manuscript. J.R.v.d.M. and S.Č. provided resources, and V.H. secured funding.

## Materials & Correspondence

Further information and requests for resources should be directed to and will be fulfilled by the corresponding author, Vassily Hatzimanikatis. The workflow and the models used in this study are available at the LCSB GitHub: https://github.com/EPFL-LCSB/SoiliSynCom. The raw data is deposited on Zenodo under accession number 10.5281/zenodo.23256682.

Genomes of the soil community members can be accessed from NCBI/ENA Bioproject number: PRJEB124582/https://www.ebi.ac.uk/ena/browser/view/PRJEB124582?show=related-records

## Declaration of generative AI and AI-assisted technologies in the writing process

During the preparation of this work, the authors used ChatGPT to improve the manuscript’s readability and language. After using this tool, the authors reviewed and edited the content as needed and take full responsibility for the content of the published article.

## References

1. Antoniewicz, M. R. A guide to deciphering microbial interactions and metabolic fluxes in microbiome communities. Current Opinion in Biotechnology 64, 230–237 (2020).

2. Jacoby, R., Peukert, M., Succurro, A., Koprivova, A. & Kopriva, S. The role of soil microorganisms in plant mineral nutrition—current knowledge and future directions. Frontiers in plant science 8, 1617 (2017).

3. Ray, P., Lakshmanan, V., Labbé, J. L. & Craven, K. D. Microbe to Microbiome: A Paradigm Shift in the Application of Microorganisms for Sustainable Agriculture. Frontiers in Microbiology 11, (2020).

4. Lawson, C. E. et al. Common principles and best practices for engineering microbiomes. Nature Reviews Microbiology 17, 725–741 (2019).

5. Stubbendieck, R. M., Vargas-Bautista, C. & Straight, P. D. Bacterial Communities: Interactions to Scale. Frontiers in Microbiology 7, (2016).

6. Gralka, M., Szabo, R., Stocker, R. & Cordero, O. X. Trophic Interactions and the Drivers of Microbial Community Assembly. Current Biology 30, R1176–R1188 (2020).

7. Machado, D. et al. Polarization of microbial communities between competitive and cooperative metabolism. Nature ecology & evolution 5, 195–203 (2021).

8. Helliwell, K. E. et al. Cyanobacteria and Eukaryotic Algae Use Different Chemical Variants of Vitamin B12. Current Biology 26, 999–1008 (2016).

9. Bunbury, F. et al. Exploring the onset of B12-based mutualisms using a recently evolved Chlamydomonas auxotroph and B12-producing bacteria. Environmental Microbiology 24, 3134– 3147 (2022).

10. Jörgensen, A. M. & Joergensen, R. G. Soil contribution to the cobalamin (vitamin B12) supply of terrestrial organisms. Biol Fertil Soils 60, 613–625 (2024).

11. Diaz-Colunga, J., Skwara, A., Vila, J. C. C., Bajic, D. & Sanchez, A. Global epistasis and the emergence of function in microbial consortia. Cell 187, 3108–3119.e30 (2024).

12. Feist, A. M., Herrgård, M. J., Thiele, I., Reed, J. L. & Palsson, B. Ø. Reconstruction of biochemical networks in microorganisms. Nat Rev Microbiol 7, 129–143 (2009).

13. Ankrah, N. Y. D. et al. Enhancing Microbiome Research through Genome-Scale Metabolic Modeling. mSystems 6, e00599–21 (2021).

14. Wang, H. et al. RAVEN 2.0: A versatile toolbox for metabolic network reconstruction and a case study on Streptomyces coelicolor. PLOS Computational Biology 14, e1006541 (2018).

15. Kanehisa, M. & Goto, S. KEGG: Kyoto Encyclopedia of Genes and Genomes. Nucleic Acids Research 28, 27–30 (2000).

16. Kanehisa, M., Furumichi, M., Sato, Y., Ishiguro-Watanabe, M. & Tanabe, M. KEGG: integrating viruses and cellular organisms. Nucleic Acids Research 49, D545–D551 (2021).

17. Thiele, I. & Palsson, B. Ø. A protocol for generating a high-quality genome-scale metabolic reconstruction. Nat Protoc 5, 93–121 (2010).

18. Orth, J. D., Thiele, I. & Palsson, B. Ø. What is flux balance analysis? Nat Biotechnol 28, 245– 248 (2010).

19. Gu, C., Kim, G. B., Kim, W. J., Kim, H. U. & Lee, S. Y. Current status and applications of genome-scale metabolic models. Genome Biology 20, 121 (2019).

20. Čaušević, S., Tackmann, J., Sentchilo, V., von Mering, C. & van der Meer, J. R. Reproducible Propagation of Species-Rich Soil Bacterial Communities Suggests Robust Underlying Deterministic Principles of Community Formation. mSystems 7, e00160–22 (2022).

21. Vayena, E. et al. A workflow for annotating the knowledge gaps in metabolic reconstructions using known and hypothetical reactions. Proceedings of the National Academy of Sciences 119, e2211197119 (2022).

22. Chinchor, N. & Sundheim, B. MUC-5 Evaluation Metrics. in Fifth Message Understanding Conference (MUC-5): Proceedings of a Conference Held in Baltimore, Maryland, August 25-27, 1993 (1993).

23. Matthews, B. W. Comparison of the predicted and observed secondary structure of T4 phage lysozyme. Biochimica et Biophysica Acta (BBA) - Protein Structure 405, 442–451 (1975).

24. Tymoshenko, S. et al. Metabolic Needs and Capabilities of Toxoplasma gondii through Combined Computational and Experimental Analysis. PLoS Comput Biol 11, e1004261 (2015).

25. Chiappino-Pepe, A., Tymoshenko, S., Ataman, M., Soldati-Favre, D. & Hatzimanikatis, V. Bioenergetics-based modeling of Plasmodium falciparum metabolism reveals its essential genes, nutritional requirements, and thermodynamic bottlenecks. PLOS Computational Biology 13, e1005397 (2017).

26. Masid, M., Ataman, M. & Hatzimanikatis, V. Analysis of human metabolism by reducing the complexity of the genome-scale models using redHUMAN. Nat Commun 11, 2821 (2020).

27. Marinos, G., Kaleta, C. & Waschina, S. Defining the nutritional input for genome-scale metabolic models: A roadmap. PLOS ONE 15, e0236890 (2020).

28. Oftadeh, O., Sahin, A., Vayena, E. & Hatzimanikatis, V. A constraint-based framework to reconstruct interaction networks in microbial communities. 2024.01.30.577913 Preprint at 10.1101/2024.01.30.577913 (2024).

29. Schäfer, M. et al. Metabolic interaction models recapitulate leaf microbiota ecology. Science 381, eadf5121 (2023).

30. Vayena, E., Ataman, M. & Hatzimanikatis, V. In silico analysis and comparison of the metabolic capabilities of different organisms by reducing metabolic complexity. Microbiome 14, 82 (2026).

31. Ratzke, C. & Gore, J. Modifying and reacting to the environmental pH can drive bacterial interactions. PLOS Biology 16, e2004248 (2018).

32. Zhalnina, K. et al. Dynamic root exudate chemistry and microbial substrate preferences drive patterns in rhizosphere microbial community assembly. Nature Microbiology 3, 470–480 (2018).

33. Wiesenbauer, J. et al. A pulse of simulated root exudation alters the composition and temporal dynamics of microbial metabolites in its immediate vicinity. Soil Biology and Biochemistry 189, 109259 (2024).

34. Wang, Y. et al. Substrate Utilization and Competitive Interactions Among Soil Bacteria Vary With Life-History Strategies. Front. Microbiol. 13, (2022).

35. Borer, B., Tecon, R. & Or, D. Spatial organization of bacterial populations in response to oxygen and carbon counter-gradients in pore networks. Nat Commun 9, 769 (2018).

36. Hammarlund, S. P., Gedeon, T., Carlson, R. P. & Harcombe, W. R. Limitation by a shared mutualist promotes coexistence of multiple competing partners. Nat Commun 12, 619 (2021).

37. Tecon, R. & Or, D. Biophysical processes supporting the diversity of microbial life in soil. FEMS Microbiology Reviews 41, 599–623 (2017).

38. Bolger, A. M., Lohse, M. & Usadel, B. Trimmomatic: a flexible trimmer for Illumina sequence data. Bioinformatics 30, 2114–2120 (2014).

39. Andrews, S. (2010) FastQC A Quality Control Tool for High Throughput Sequence Data. - References - Scientific Research Publishing. https://www.scirp.org/reference/referencespapers.

40. Kolmogorov, M., Yuan, J., Lin, Y. & Pevzner, P. A. Assembly of long, error-prone reads using repeat graphs. Nat Biotechnol 37, 540–546 (2019).

41. Wick, R. R., Schultz, M. B., Zobel, J. & Holt, K. E. Bandage: interactive visualization of de novo genome assemblies. Bioinformatics 31, 3350–3352 (2015).

42. Sović, I. et al. Fast and sensitive mapping of nanopore sequencing reads with GraphMap. Nat Commun 7, 11307 (2016).

43. Vaser, R., Sović, I., Nagarajan, N. & Šikić, M. Fast and accurate de novo genome assembly from long uncorrected reads. Genome Res 27, 737–746 (2017).

44. Langmead, B. & Salzberg, S. L. Fast gapped-read alignment with Bowtie 2. Nat Methods 9, 357–359 (2012).

45. Walker, B. J. et al. Pilon: an integrated tool for comprehensive microbial variant detection and genome assembly improvement. PLoS One 9, e112963 (2014).

46. Gurevich, A., Saveliev, V., Vyahhi, N. & Tesler, G. QUAST: quality assessment tool for genome assemblies. Bioinformatics 29, 1072–1075 (2013).

47. Li, H. & Durbin, R. Fast and accurate short read alignment with Burrows–Wheeler transform. Bioinformatics 25, 1754–1760 (2009).

48. Twelve years of SAMtools and BCFtools | GigaScience | Oxford Academic. https://academic.oup.com/gigascience/article/10/2/giab008/6137722.

49. Seemann, T. Prokka: rapid prokaryotic genome annotation. Bioinformatics 30, 2068–2069 (2014).

50. Parks, D. H. et al. A complete domain-to-species taxonomy for Bacteria and Archaea. Nat Biotechnol 38, 1079–1086 (2020).

51. Chaumeil, P.-A., Mussig, A. J., Hugenholtz, P. & Parks, D. H. GTDB-Tk v2: memory friendly classification with the genome taxonomy database. Bioinformatics 38, 5315–5316 (2022).

52. Jones, D. M. Manual of Methods for General Bacteriology. Journal of Clinical Pathology 34, 1069–1069 (1981).

53. Monk, J. M. et al. iML1515, a knowledgebase that computes Escherichia coli traits. Nat Biotechnol 35, 904–908 (2017).

54. Hadadi, N. et al. Mechanistic insights into bacterial metabolic reprogramming from omics-integrated genome-scale models. npj Syst Biol Appl 6, 1–11 (2020).

55. Karp, P. D. et al. The BioCyc collection of microbial genomes and metabolic pathways. Brief Bioinform 20, 1085–1093 (2019).

56. Chang, A. et al. BRENDA, the ELIXIR core data resource in 2021: new developments and updates. Nucleic Acids Res 49, D498–D508 (2021).

57. The UniProt Consortium. UniProt: the universal protein knowledgebase in 2021. Nucleic Acids Research 49, D480–D489 (2021).

58. Sayers, E. W. et al. Database resources of the National Center for Biotechnology Information in 2025. Nucleic Acids Res 53, D20–D29 (2025).

59. Soh, K. C. & Hatzimanikatis, V. Constraining the Flux Space Using Thermodynamics and Integration of Metabolomics Data. in Metabolic Flux Analysis: Methods and Protocols (eds Krömer, J. O., Nielsen, L. K. & Blank, L. M.) 49–63 (Springer, New York, NY, 2014). doi:10.1007/978-1-4939-1170-7_3.

60. Salvy, P. et al. pyTFA and matTFA: a Python package and a Matlab toolbox for Thermodynamics-based Flux Analysis. Bioinformatics 35, 167–169 (2019).

61. Lieven, C. et al. MEMOTE for standardized genome-scale metabolic model testing. Nat Biotechnol 38, 272–276 (2020).

62. Lipscomb, C. E. Medical Subject Headings (MeSH). Bull Med Libr Assoc 88, 265–266 (2000).

63. Malik, A. et al. ChEBI: re-engineered for a sustainable future. Nucleic Acids Res 54, D1768– D1778 (2026).

64. Kim, S. et al. PubChem 2025 update. Nucleic Acids Res 53, D1516–D1525 (2025).

65. Jaccard, P. THE DISTRIBUTION OF THE FLORA IN THE ALPINE ZONE.^1^. New Phytologist 11, 37–50 (1912).

66. Chiappino-Pepe, A. & Hatzimanikatis, V. PhenoMapping: a protocol to map cellular phenotypes to metabolic bottlenecks, identify conditional essentiality, and curate metabolic models. STAR Protoc 2, 100280 (2021).

67. Seaver, S. M. D. et al. The ModelSEED Biochemistry Database for the integration of metabolic annotations and the reconstruction, comparison and analysis of metabolic models for plants, fungi and microbes. Nucleic Acids Res 49, D575–D588 (2021).

68. King, Z. A. et al. BiGG Models: A platform for integrating, standardizing and sharing genome-scale models. Nucleic Acids Research 44, D515–D522 (2016).

69. Moretti, S. et al. MetaNetX/MNXref – reconciliation of metabolites and biochemical reactions to bring together genome-scale metabolic networks. Nucleic Acids Research 44, D523–D526 (2016).

70. Weininger, D. SMILES. 3. DEPICT. Graphical depiction of chemical structures. J. Chem. Inf. Comput. Sci. 30, 237–243 (1990).

71. Jankowski, M. D., Henry, C. S., Broadbelt, L. J. & Hatzimanikatis, V. Group contribution method for thermodynamic analysis of complex metabolic networks. Biophys J 95, 1487–1499 (2008).

72. Heirendt, L. et al. Creation and analysis of biochemical constraint-based models using the COBRA Toolbox v.3.0. Nat Protoc 14, 639–702 (2019).

73. Gurobi Optimization, LLC. Gurobi Optimizer Reference Manual. (2024).

74. I. I. CPLEX (2009) V12.1 User’s Manual for CPLEX. Vol. 46, International Business Machines Corporation, Armonk, 157. - References - Scientific Research Publishing. https://www.scirp.org/reference/referencespapers.

