## SupplementaryInformation for "Genome-scale metabolic models for a synthetic soil microbial community as a path for understanding community functioning"

### Genome-scale metabolic models and analyses of 21 soil isolates

Supplementary Figures S1-S18 present experimental growth trajectories, genome assembly statistics, model content and curation, nutrient requirements, feasible metabolite exchanges, and biosynthetic analyses. Supplementary Tables S1-S3 list the genome assembly statistics, the biochemical composition of the DiMEs environments and the PTYG medium composition.

Strain codes are defined in the manuscript Table 1. Genus names and taxonomic color groups retain the labels used in the figures, whereas Table 1 reports the updated Genome Taxonomy Database (GTDB) assignments. Unless a different key is shown, taxonomic colours are:

Actinobacteria   Alphaproteobacteria   Betaproteobacteria
Gammaproteobacteria   Bacteroidetes   Firmicutes

| Item | Content |
|---|---|
| Figure S1 | Growth trajectories of the soil isolates on succinate |
| Figure S2 | Growth trajectories of the soil isolates in PTYG |
| Figure S3 | Genome content and confusion metrics |
| Figure S4 | Nutrient demand and core requirements across alternative in silico minimal media |
| Figure S5 | Growth of *Microbacterium* (mip) in iMM-predicted medium |
| Figure S6 | Distribution of feasible metabolites across strains |
| Figure S7 | Candidate metabolite exchange links across 15 metabolite classes |
| Figure S8 | Biosynthetic pathway similarity in PTYG |
| Figure S9 | Biosynthetic pathway similarity and gene requirements in minimal medium |
| Figure S10 | Medium-dependent gene requirements for biomass-building-block production |
| Figure S11 | Contiguity and composition of the genome assemblies used for model reconstruction |
| Figure S12 | Evidence supporting model gap filling |
| Figure S13 | Model quality scores and thermodynamic information |
| Figure S14 | Effect of biomass-yield aggregation on predicted exchange and niche overlap |
| Figure S15 | Predicted exchange profiles |
| Figure S16 | Reaction content and similarity of central metabolic networks |
| Figure S17 | Genome and metabolic-model characteristics of the 21 strains |
| Figure S18 | Model-feasible exchange of metabolites listed in the experimental analysis |
| Table S1 | PTYG medium composition |
| Table S2 | Biochemical composition of the basal and conditioned environments |
| Table S3 | Statistics of the genome assemblies used for model reconstruction |

**Supplementary Figure S1 Sheet 1 of 2**

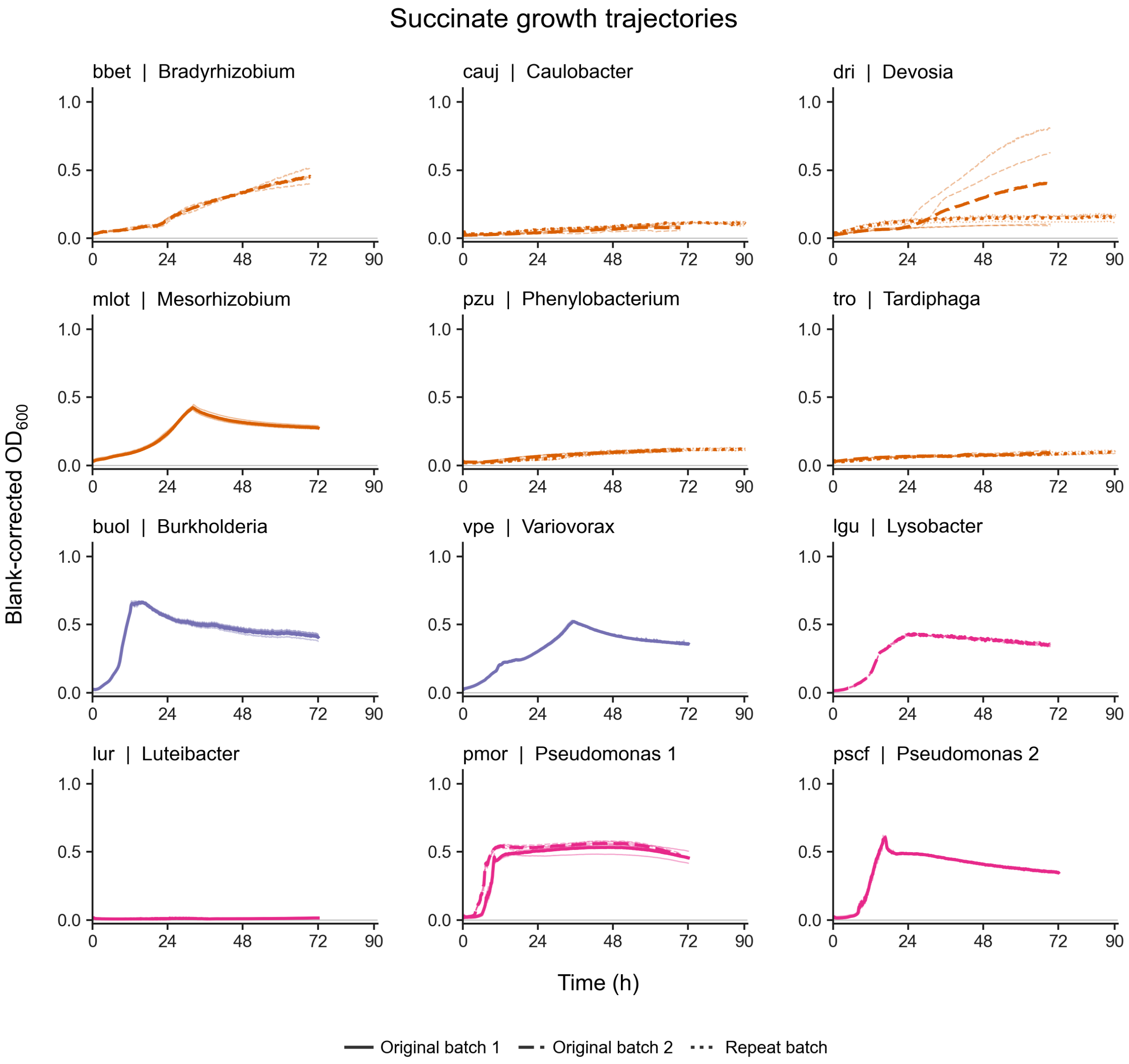


**Growth trajectories of the soil isolates on succinate**. Optical density at 600 nm ($OD_{600}$) was measured for 21 isolates in 21C minimal medium containing 10 mM succinate. Thin lines show individual wells and thick lines show the mean of four wells per strain and batch, after subtraction of the matching uninoculated controls. Solid, dashed and dotted lines denote original batch 1, original batch 2 and the repeat batch, respectively, where available. Batches are not pooled, and each curve ends at its last measurement. All wells are shown without smoothing. Axis limits are shared across strains. Colors identify taxonomic plotting groups, and genus labels follow the culture records. The remaining nine isolates are shown on the next sheet.

**Supplementary Figure S1 Sheet 2 of 2**

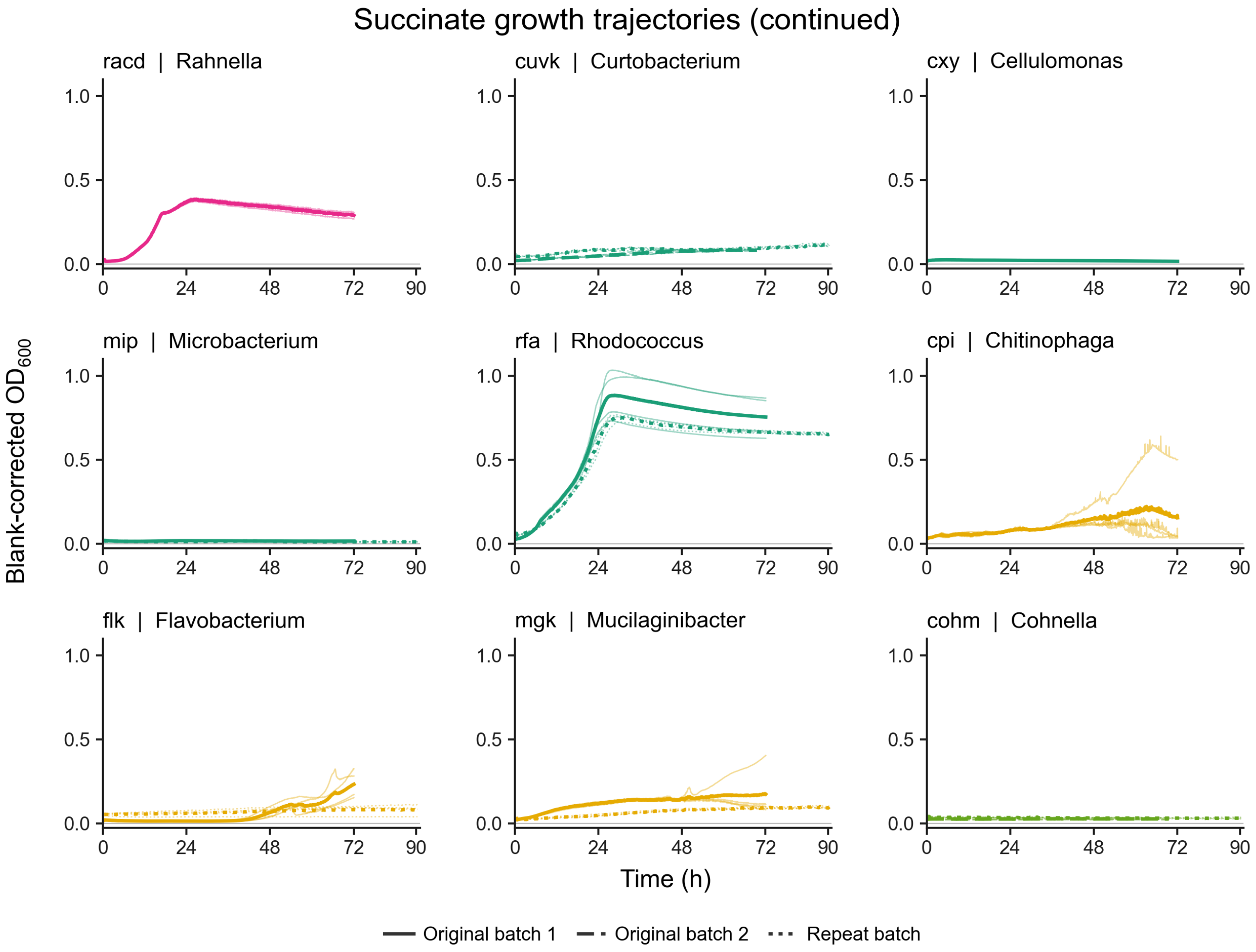


**Growth trajectories of the soil isolates on succinate (continued)**. The remaining nine isolates from the experiment are shown on the preceding sheet. Thin lines represent individual wells and thick lines represent four-well means after correction against matching uninoculated controls. Solid, dashed and dotted lines denote original batch 1, original batch 2 and the repeat batch, respectively. The two sheets together show all 21 isolates with common axis limits; colors identify taxonomic plotting groups.

**Supplementary Figure S2 Sheet 1 of 2**

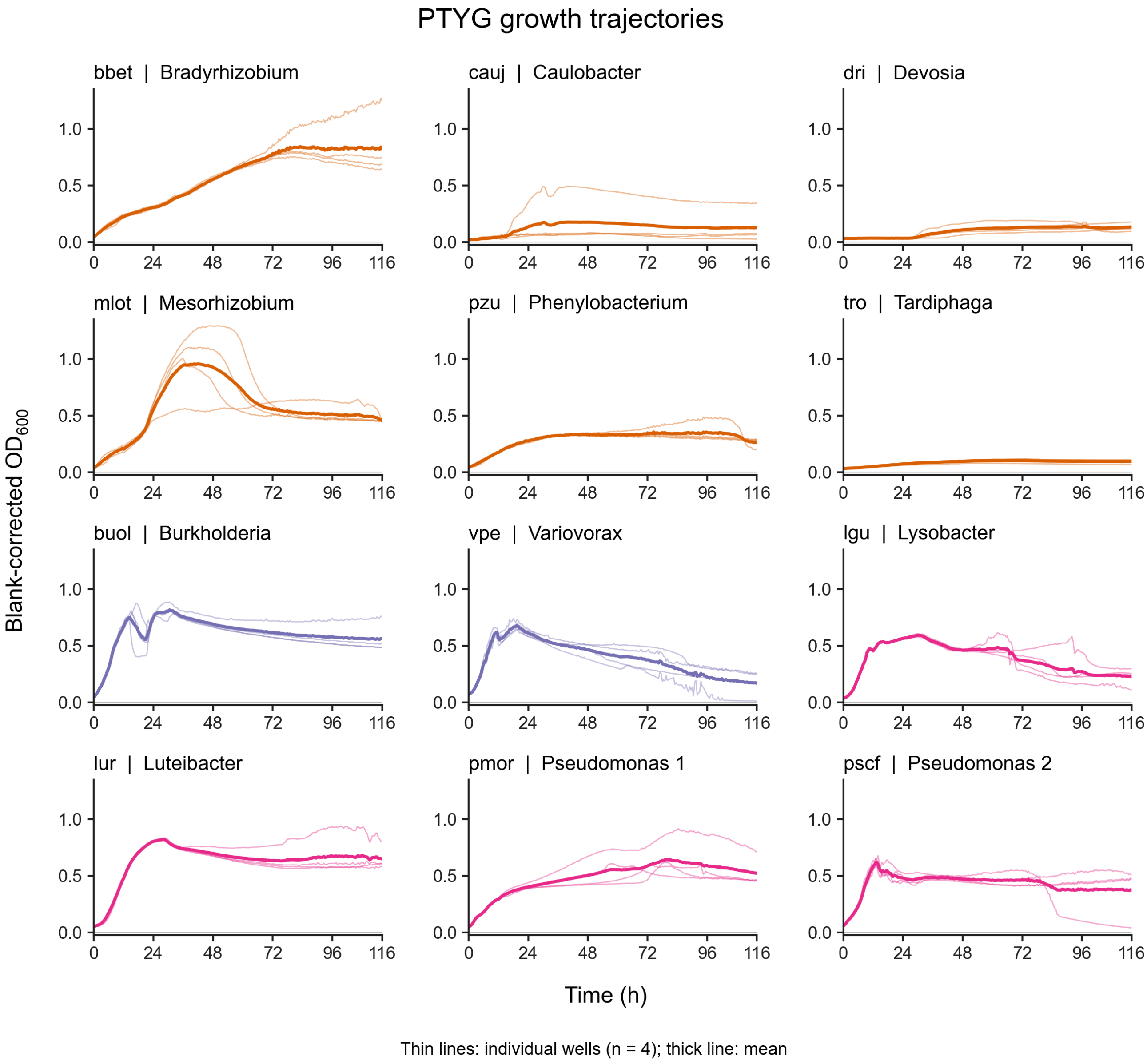


**Growth trajectories of the soil isolates in PTYG**. Optical density at 600 nm ($OD_{600}$) was measured for 21 isolates in peptone-tryptone-yeast extract-glucose (PTYG) medium. Thin lines show four individual wells per strain and thick lines show their mean after correction against matching uninoculated medium controls. All wells, including plate-edge wells, are retained without smoothing or outlier removal. Measurements span approximately 0.5-116 h, with common axis limits across strains. Colors identify taxonomic plotting groups, and genus labels follow the culture records. The remaining nine isolates are shown on the next sheet.

**Supplementary Figure S2 Sheet 2 of 2**

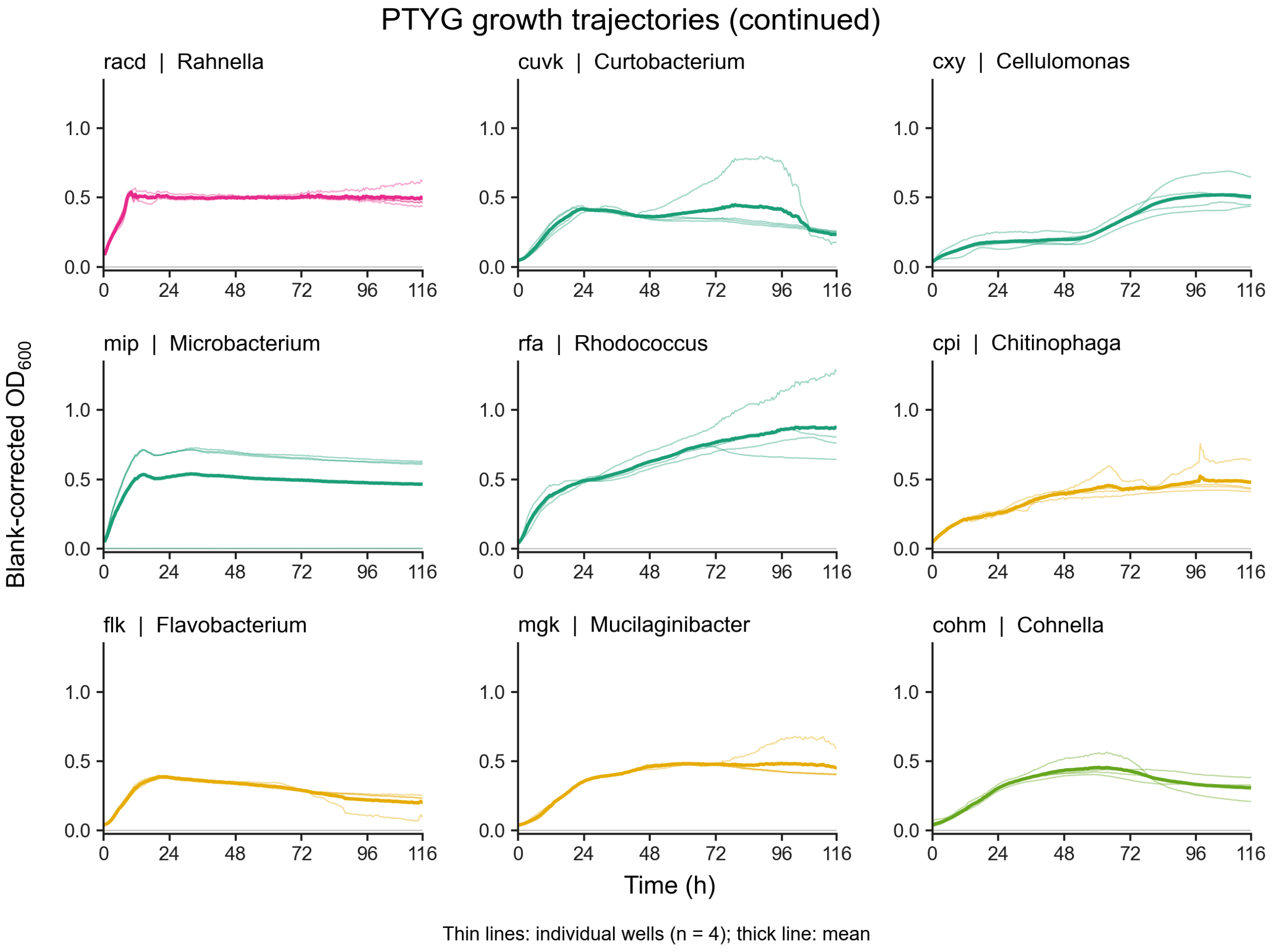


**Growth trajectories of the soil isolates in PTYG (continued)**. The remaining nine isolates from the peptone-tryptone-yeast extract-glucose (PTYG) experiment shown on the preceding sheet. Thin lines represent individual wells and thick lines represent four-well means after correction against matching uninoculated medium controls. All wells are retained. The two sheets together show all 21 isolates with common axis limits; colors identify taxonomic plotting groups.

### Supplementary Figure S3 Sheet 1 of 2

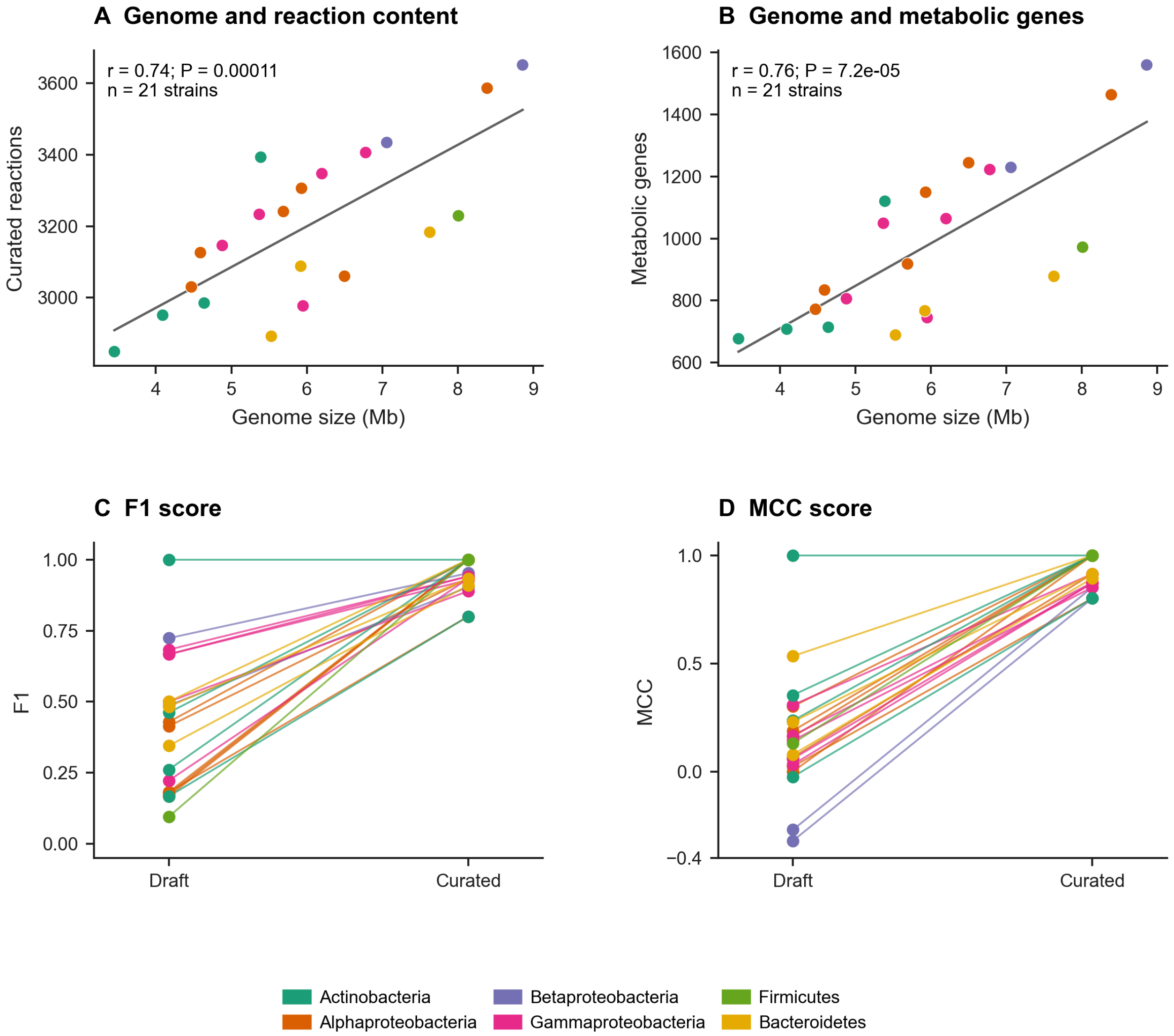


**Genome content and confusion metrics**. Genome assembly length is plotted against curated-model reaction counts (A) and model-associated metabolic gene counts (B) for 21 strains. Points represent strains; colors identify the taxonomic groups in the legend. Lines show linear fits, with Pearson correlation coefficients (r) and two-sided P values. (C, D) F1 scores and Matthews correlation coefficients (MCC), respectively, for draft and curated models. Lines connect the same strain, and a score of 1 indicates perfect agreement with the growth records used for curation. Panel E on the next sheet shows the corresponding substrate-level classifications. Two reference entries involving cxy and lur differ between the scoring records and the phenotype matrix displayed in the main manuscript.

**Supplementary Figure S3 Sheet 2 of 2**

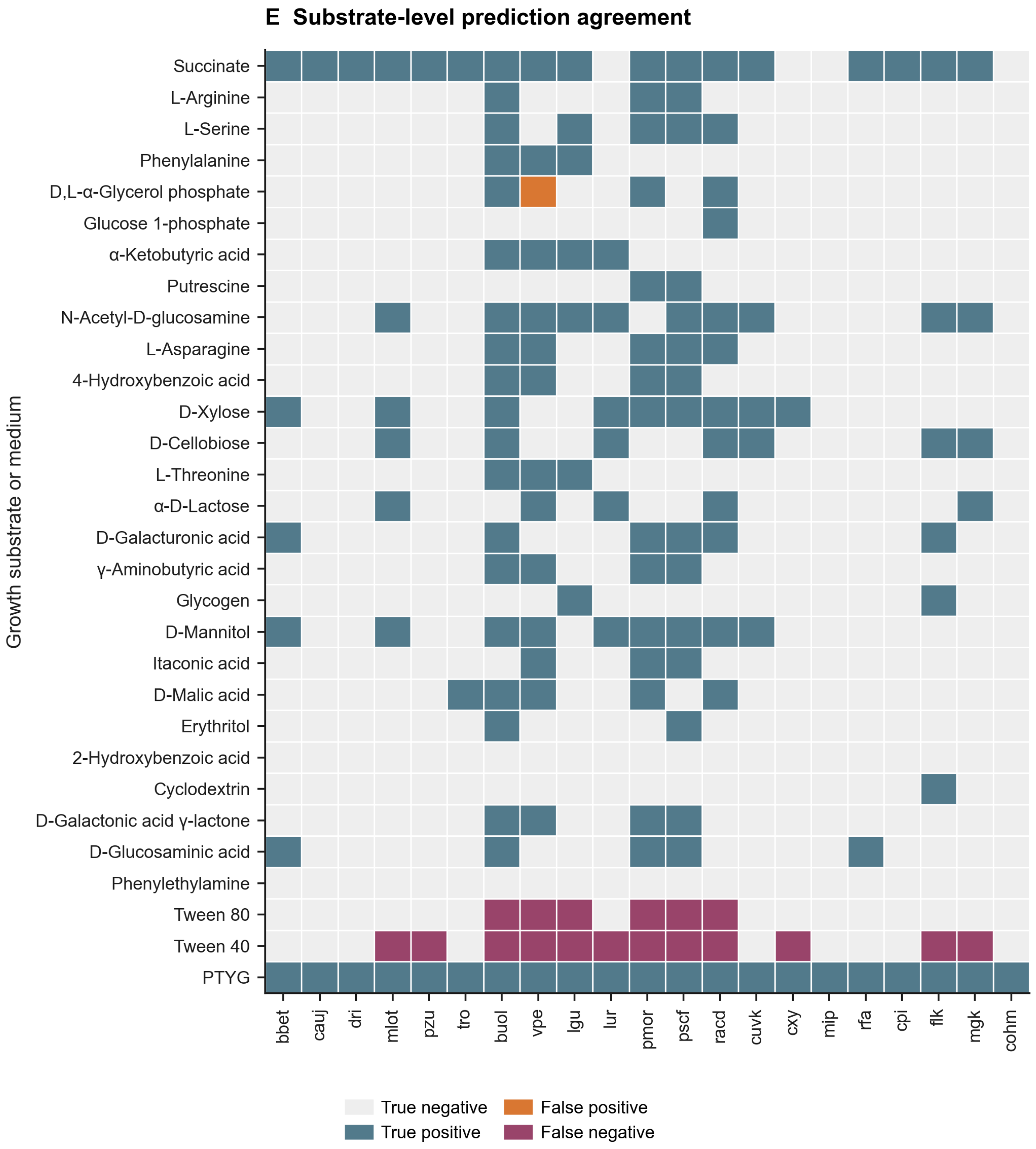


**Genome content and confusion metrics (continued)**. (E) Curated-model predictions compared with the growth records used during curation. Rows represent individual substrate conditions or peptone-tryptone-yeast extract-glucose (PTYG) medium; columns represent strains. Blue and light grey indicate correct predictions of growth and no growth, respectively. Orange marks predicted growth without recorded growth (false positives), and purple marks recorded growth that was not predicted (false negatives).

**Supplementary Figure S4**

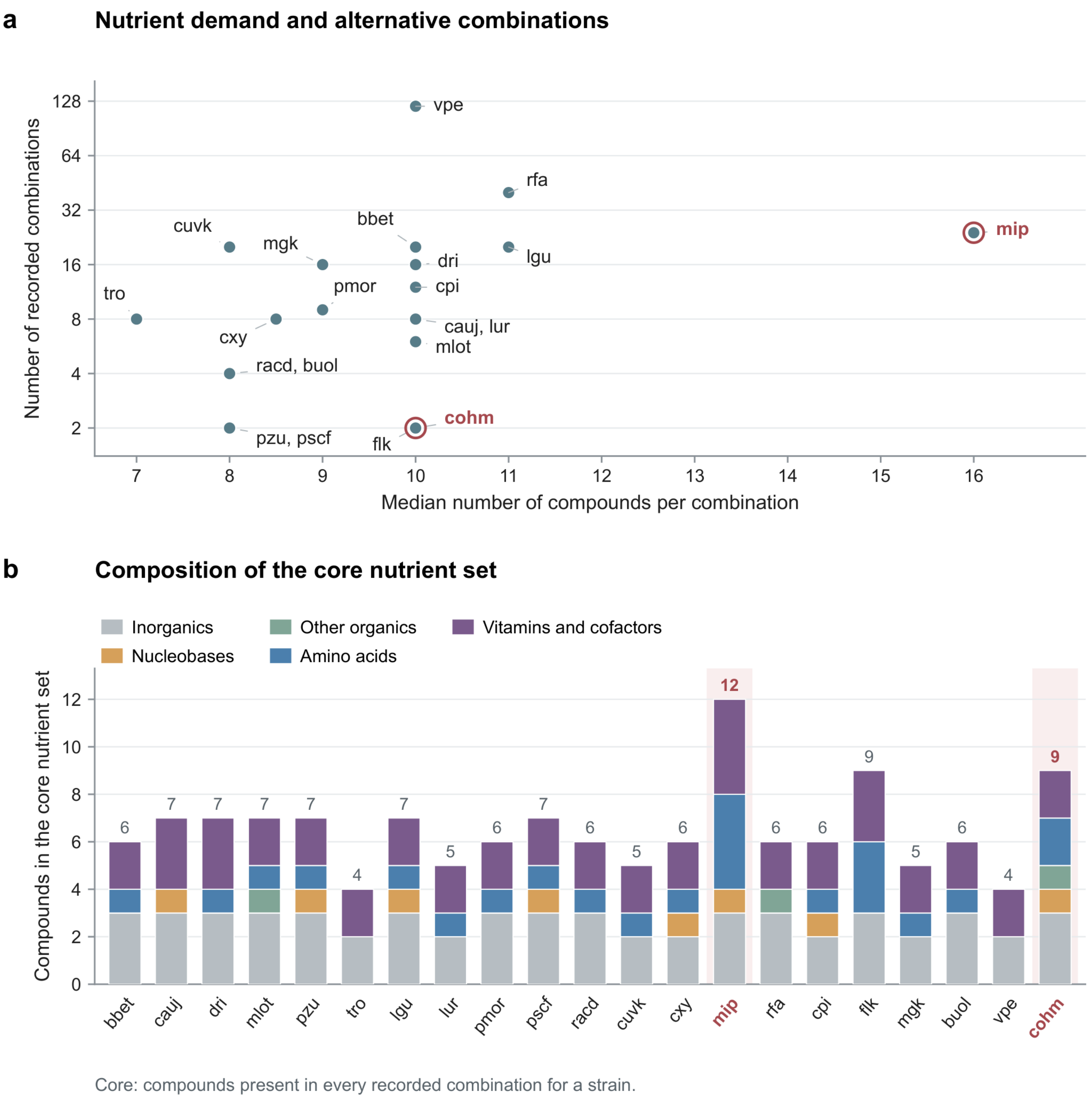


**Nutrient demand and core requirements across alternative in silico minimal media.** Recorded iMM predictions were compared across 21 strain models using the PTYG-derived pool of 69 compounds. (A) Relationship between the median number of supplied compounds per combination and the number of recorded combinations for each strain. The vertical axis is logarithmic. Labels at shared positions identify strains with identical values. The PTYG-restricted strains mip and cohm are highlighted in red. (B) Chemical composition of each strain's core nutrient set, defined as the compounds present in every recorded combination. Stacked bars distinguish amino acids, vitamins and cofactors, nucleobases, inorganic compounds and other organic compounds; numbers above bars give the total core size. The mip core contains four amino acids and four vitamins or cofactors, compared with two of each in cohm.

**Supplementary Figure S5**

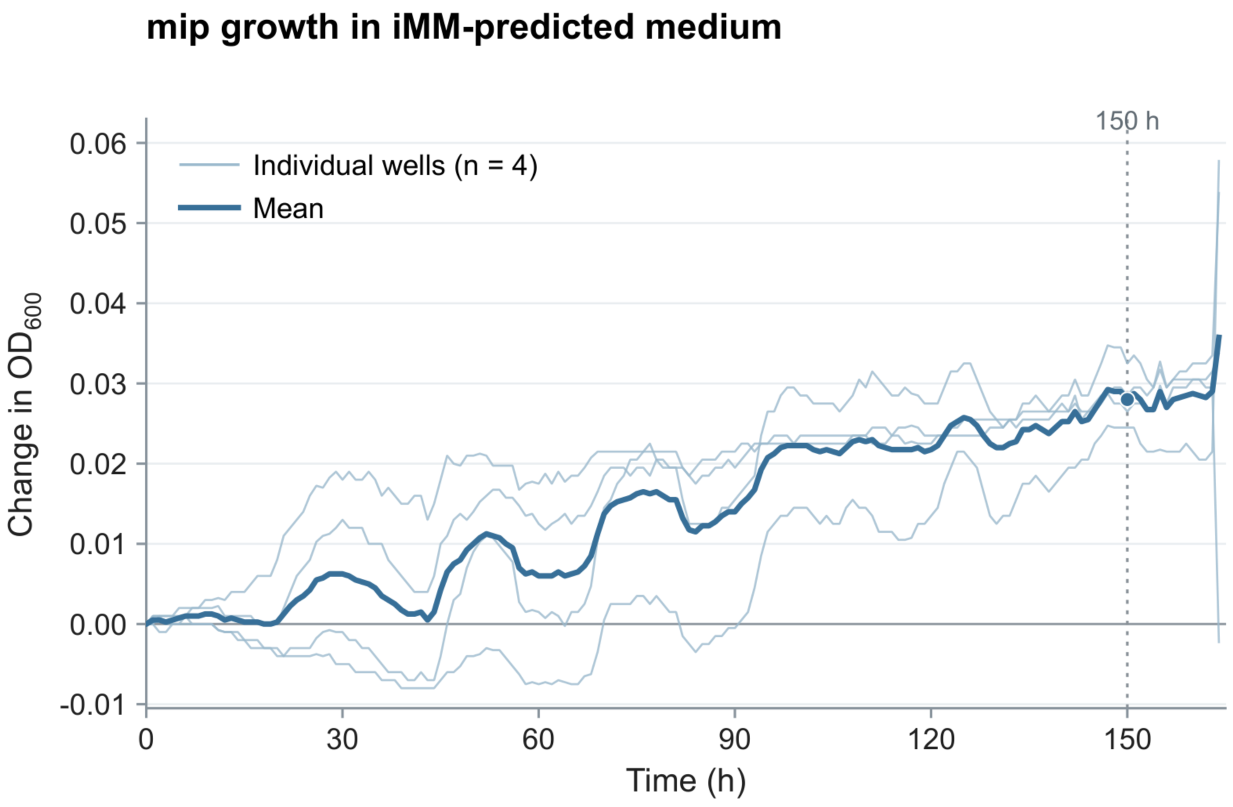


**Growth of *Microbacterium* (mip) in iMM-predicted medium.** Thin lines show the four individual inoculated wells, and the thick blue line shows their mean. Optical density at 600 nm ($OD_{600}$) was corrected at each time point by subtracting the mean of four matching uninoculated medium wells, then expressed as the change from each well's corrected initial reading. The dotted vertical line marks 150 h, when the mean increase in $OD_{600}$ was 0.0280 ± 0.0034 (mean ± SD; n = 4 wells).

**Supplementary Figure S6**

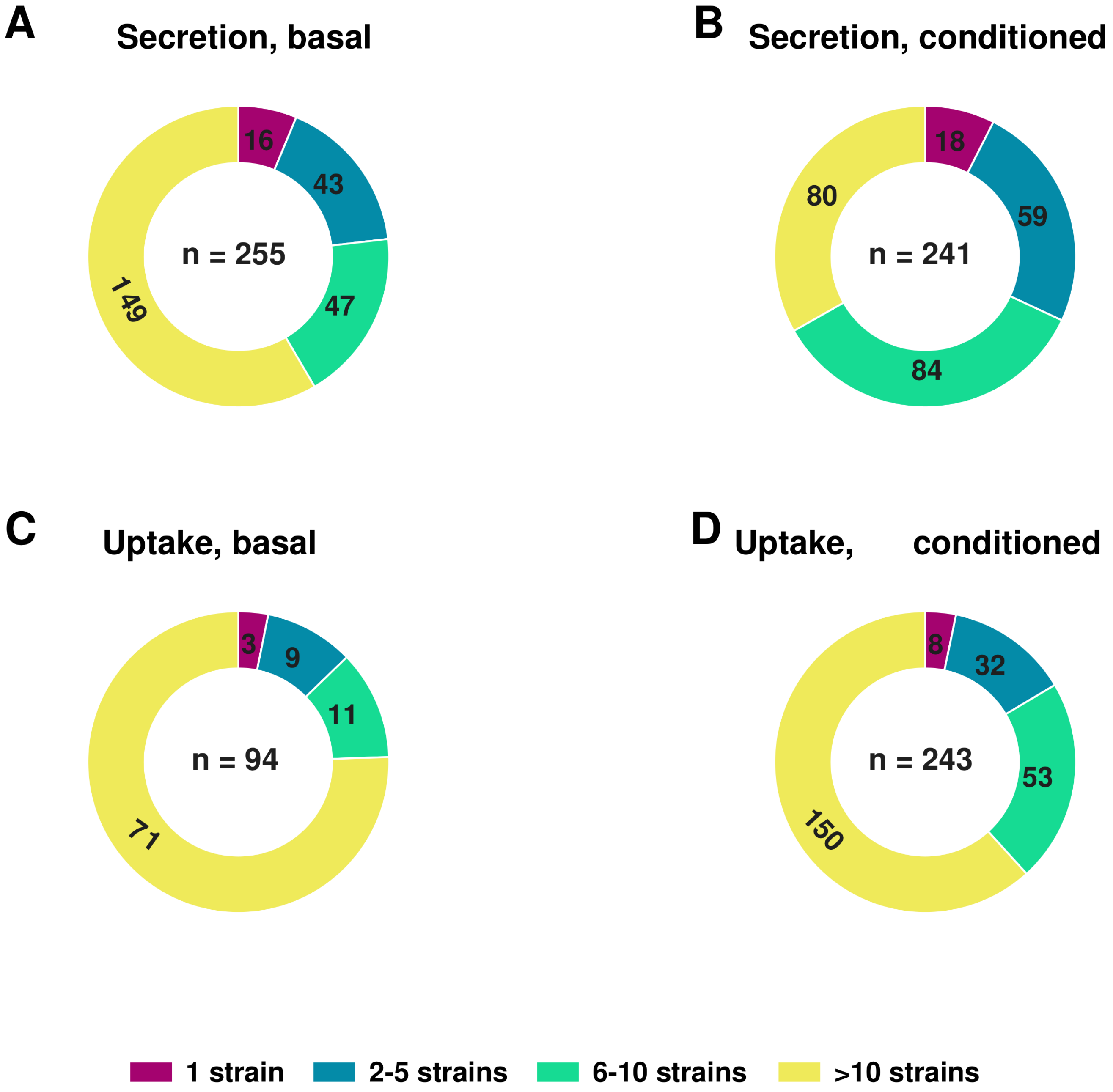


**Distribution of feasible metabolites across strains**. Donut charts group distinct metabolites by the number of strains predicted to secrete them in basal (A) and conditioned (B) environments, or take them up in basal (C) and conditioned (D) environments. Colors distinguish metabolites associated with one strain, 2-5 strains, 6-10 strains or more than 10 strains. Segment labels give metabolite counts; centre labels give the total number of distinct metabolites in each panel. Each metabolite is counted once per panel. Basal results cover 20 strains (*mip* fails to grow) and conditioned results cover 21. Profiles combine DiMEs (Decomposed in silico Minimal Exchanges) solutions across biomass yields of 10-100% of the optimum.

### Supplementary figure S7 Sheet 1 of 2

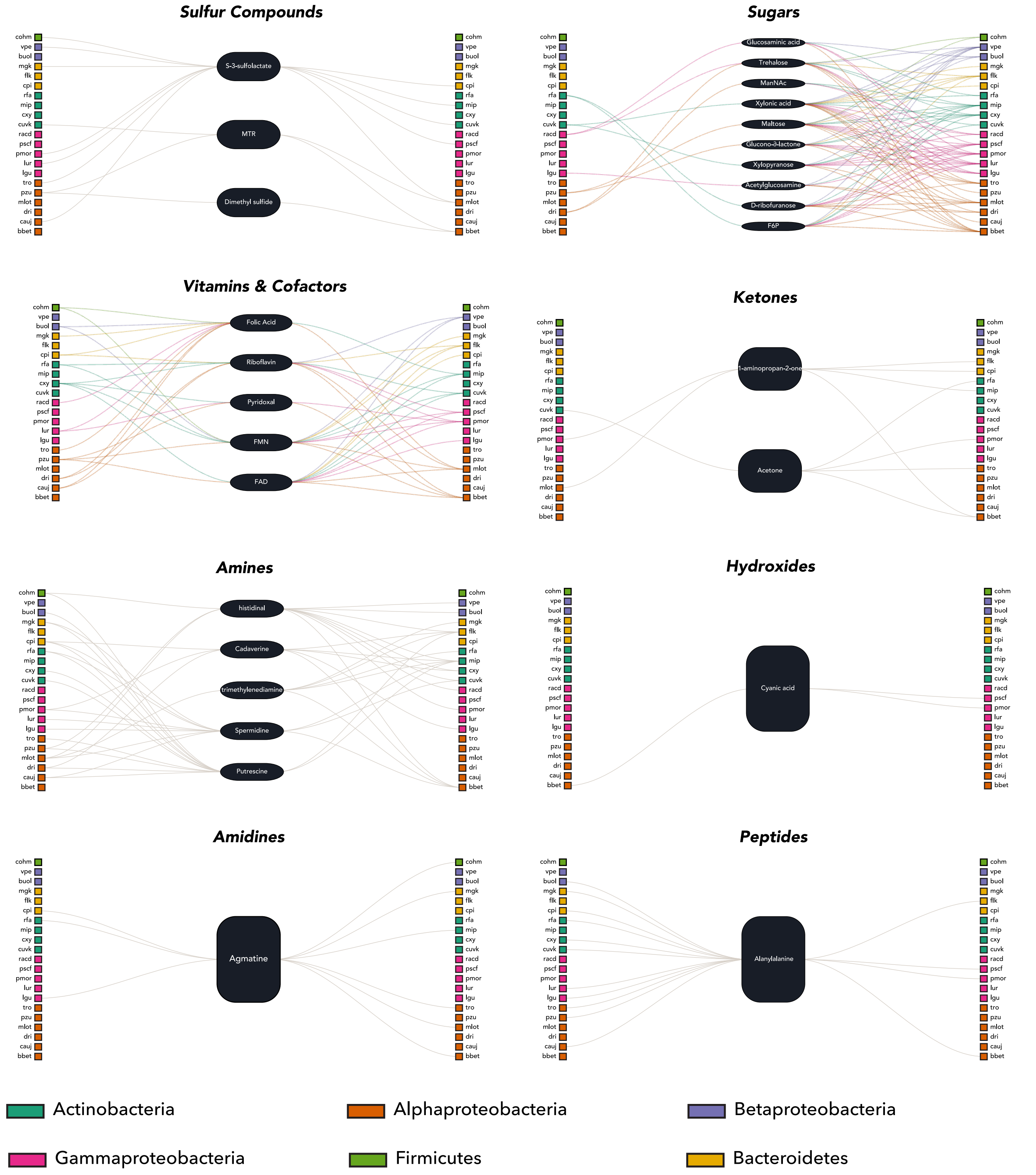


**Candidate metabolite exchange links across 15 metabolite classes.** The first eight class-resolved maps are shown here; the remaining seven and the full caption continue on the next page. Strains on the left are predicted producers, central nodes are metabolites, and strains on the right are potential consumers. Colors identify taxonomic groups.

### Supplementary figure S7 Sheet 2 of 2

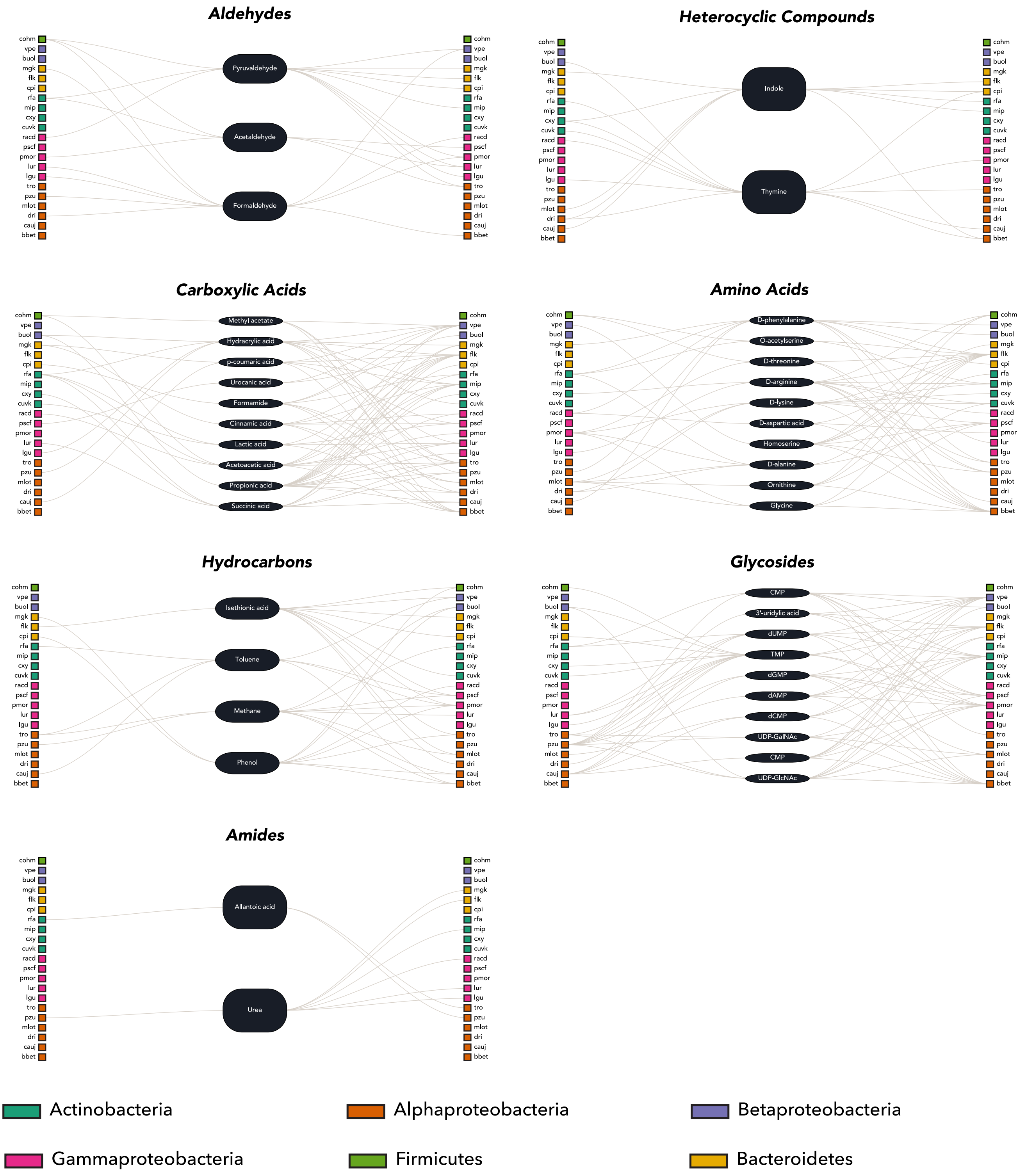


**Candidate metabolite exchange links across 15 metabolite classes (Continued).** Sankey diagrams connect predicted producers in the basal environment (left), metabolites (centre), and potential consumers after community conditioning (right), using pooled Decomposed in silico Minimal Exchanges (DiMEs) profiles. Links identify metabolites predicted to be secreted in the basal environment and to become newly feasible uptake substrates after the resource pool was expanded with predicted community secretions. For each metabolite, strains predicted to both produce and consume it were omitted from both sides. Up to ten metabolites per class are shown, prioritising compounds with few predicted producers and multiple potential consumers. Strain colors indicate taxonomic groups. Connection widths increase with the number of associated producer–consumer combinations and are scaled separately within each panel; they do not represent exchange flux. The figure includes sugars, carboxylic acids, and vitamins and cofactors shown in the main figure, together with twelve additional classes. These maps identify candidate exchanges rather than experimentally measured metabolite transfer.

### Supplementary Figure S8

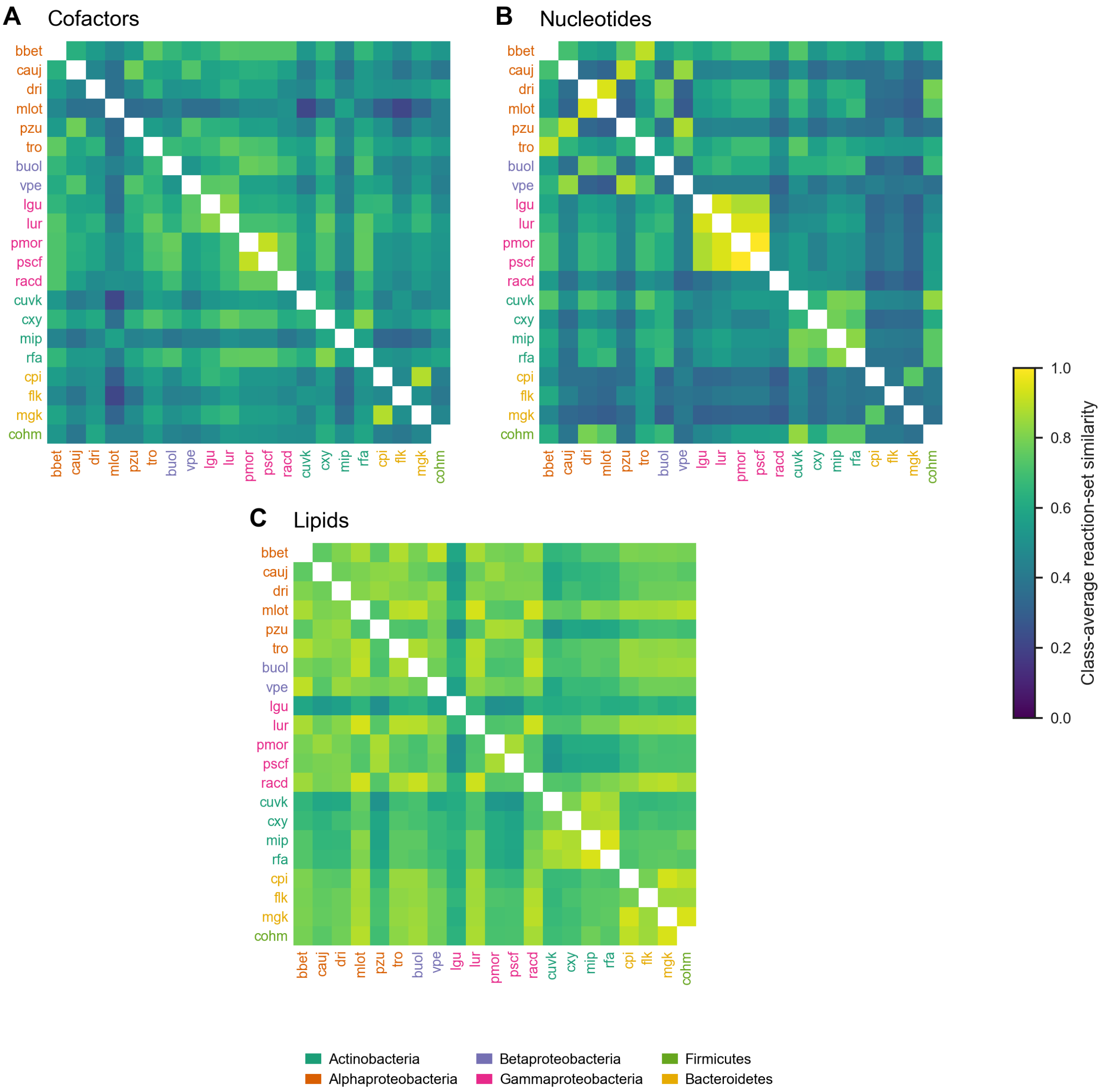


**Biosynthetic pathway similarity in PTYG**. Pairwise similarity of NIS-derived pathways for producing cofactors (A), nucleotides (B) and lipids (C) in peptone-tryptone-yeast extract-glucose (PTYG) medium for 21 strains. Rows and columns represent strains. For each biomass building block, reactions from all recovered routes were combined, and Jaccard similarities were averaged within each class. A value of 1 denotes identical recovered reaction sets across the compared components. Strain-label colors identify the taxonomic groups in the legend.

**Supplementary Figure S9 Sheet 1 of 3**

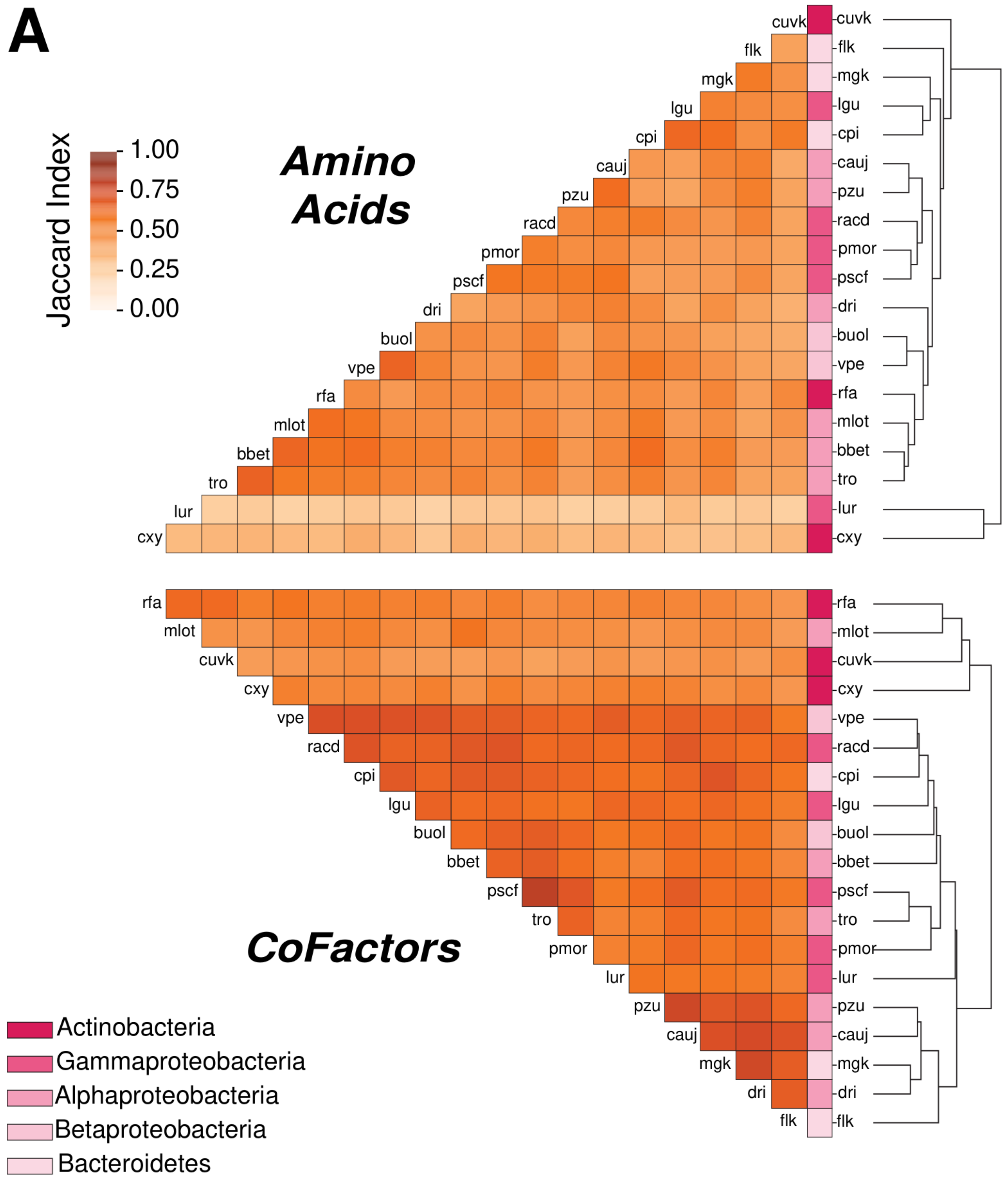


**Biosynthetic pathway similarity and gene requirements in minimal medium**. (A) Class-averaged Jaccard similarity of NIS-derived biosynthetic reaction sets in M9 with succinate for 19 strains; mip and cohm are excluded. This sheet shows amino acids and cofactors. For each biomass building block (BBB), reactions from all recovered routes were combined before pairwise comparison; similarities were then averaged within each BBB class. Missing routes contribute zero. Dendrograms show hierarchical clustering, and coloured strips identify the taxonomic groups in the legend.

**Supplementary Figure S9 Sheet 2 of 3**

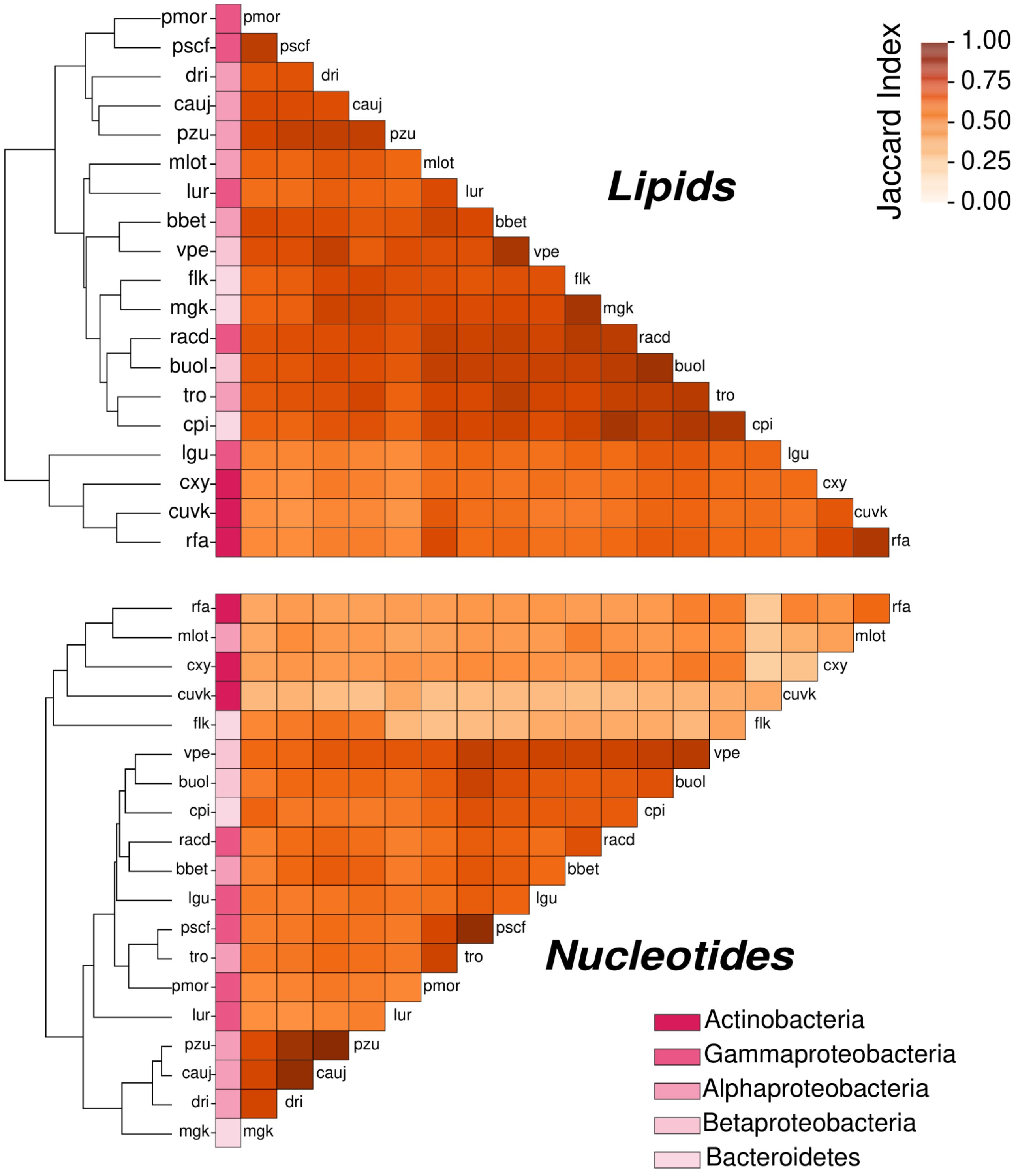


**Biosynthetic pathway similarity and gene requirements in minimal medium (continued)**. Panel (A) continues with lipid and nucleotide biosynthesis for the same 19 strains in M9 with succinate. Jaccard similarity is the fraction of reactions shared by two recovered reaction sets relative to their union, averaged across biomass building blocks within each class. Higher values indicate greater similarity. Missing routes contribute zero, so low values need not indicate dissimilarity between two recovered pathways. Dendrograms show hierarchical clustering; coloured strips identify the taxonomic groups in the legend.

**Supplementary Figure S9 Sheet 3 of 3**

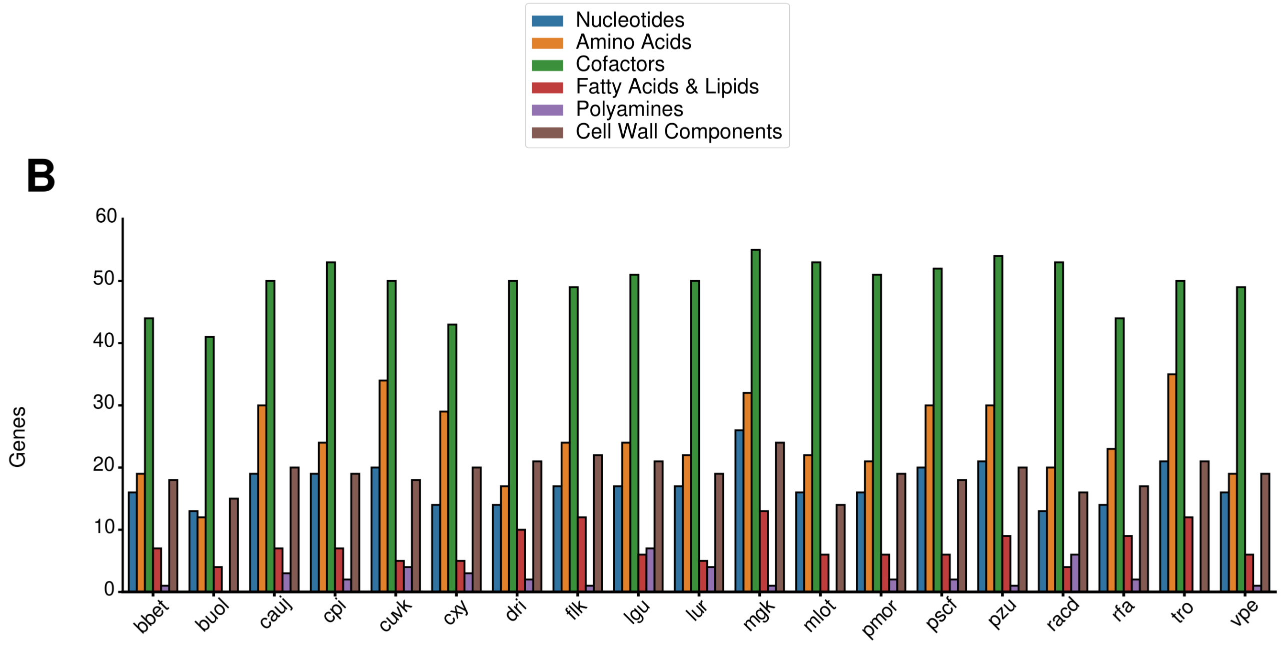


**Biosynthetic pathway similarity and gene requirements in minimal medium (continued)**. (B) Genes required for the production of biomass building blocks (BBBs) in M9 with succinate, grouped into six classes: nucleotides, amino acids, cofactors, fatty acids and lipids, polyamines, and cell-wall components. A gene is counted for a class when its deletion reduces predicted production of an assessed BBB in that class below 10% of the WT model's biomass flux. Bars show class-specific counts for 19 strains; mip and cohm are excluded.

### Supplementary Figure S10 Sheet 1 of 2

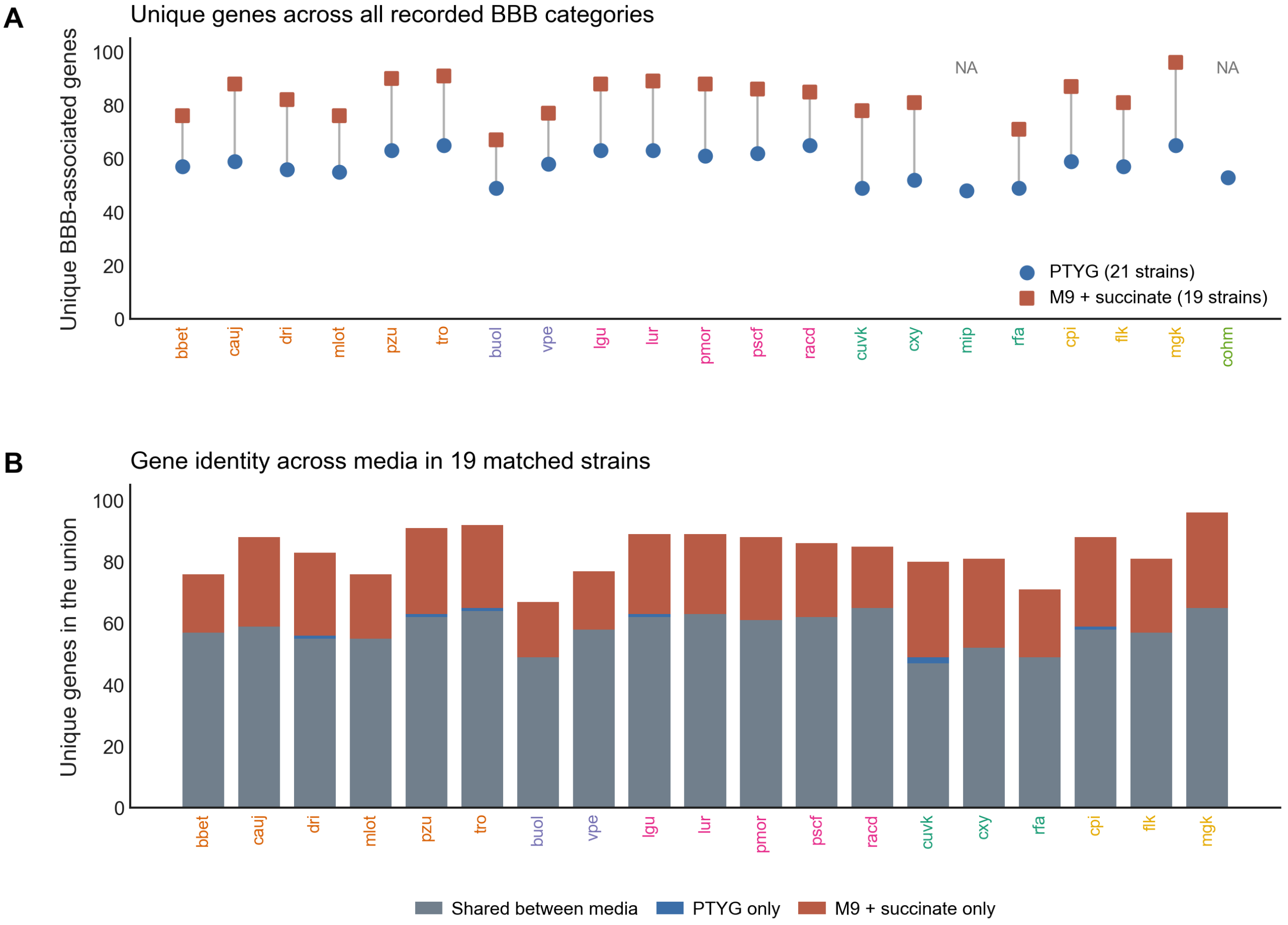


**Medium-dependent gene requirements for biomass-building-block production**. (A) Number of distinct genes whose deletion reduced predicted flux of at least one assessed biomass building block (BBB) below 10% of the undeleted model's value. Blue circles show peptone-tryptone-yeast extract-glucose (PTYG) results for 21 strains; red squares show M9 with succinate results for 19 strains. Lines connect the same strain. (B) Genes shared between the two media or recovered only in one medium for the 19 matched strains. Grey, blue and red bar segments indicate shared, PTYG-only and M9-with-succinate-only genes, respectively.

**Supplementary Figure S10 Sheet 2 of 2**

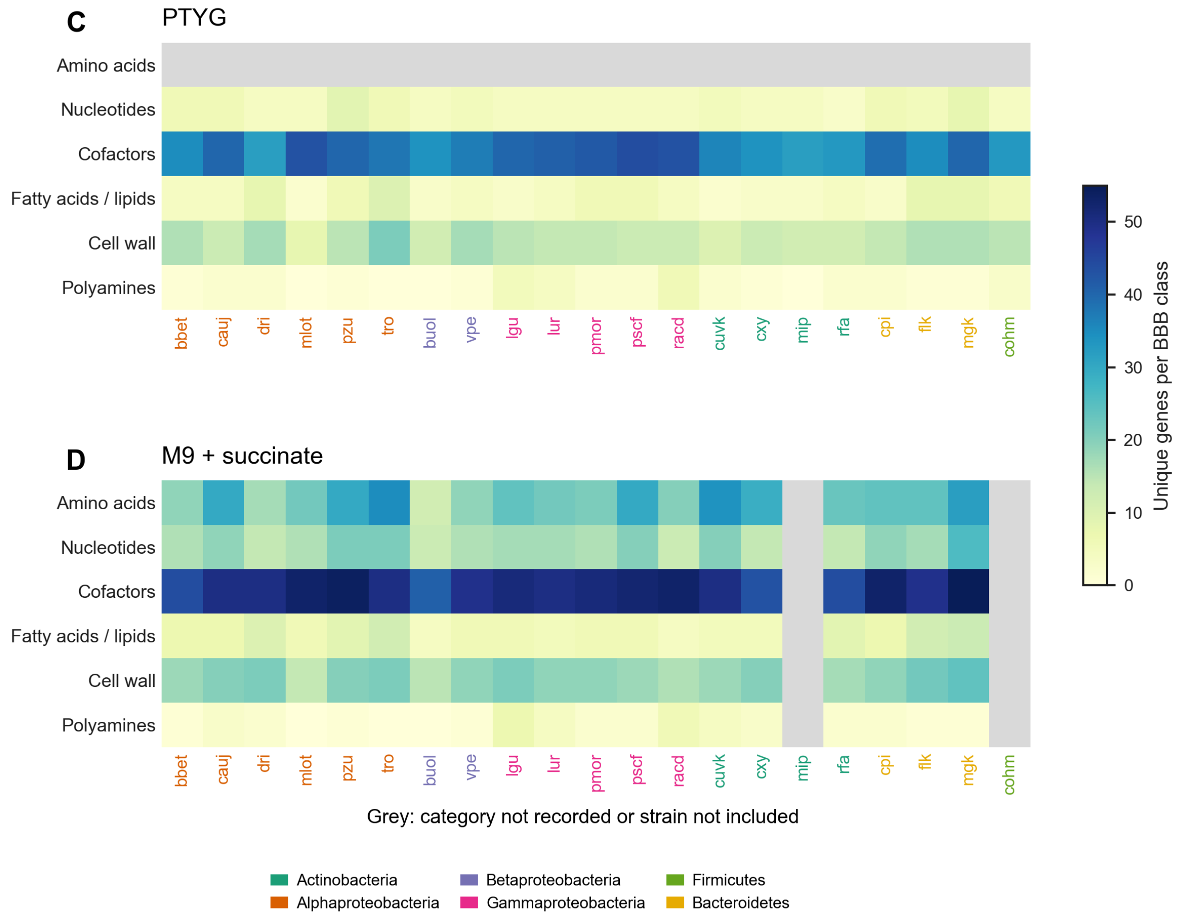


**Medium-dependent gene requirements for biomass-building-block production** (continued). Class-specific gene counts in PTYG (C) and M9 with succinate (D). A gene is counted when its deletion reduces production of an assessed biomass building block (BBB) in that class below 10% of the WT model's flux. Darker cells indicate more genes. Grey marks an unrecorded category or an unavailable strain-medium result, rather than zero. Genes can contribute to several classes, so class counts are not additive. Strain-label colors identify the taxonomic groups in the legend.

**Supplementary Figure S11**

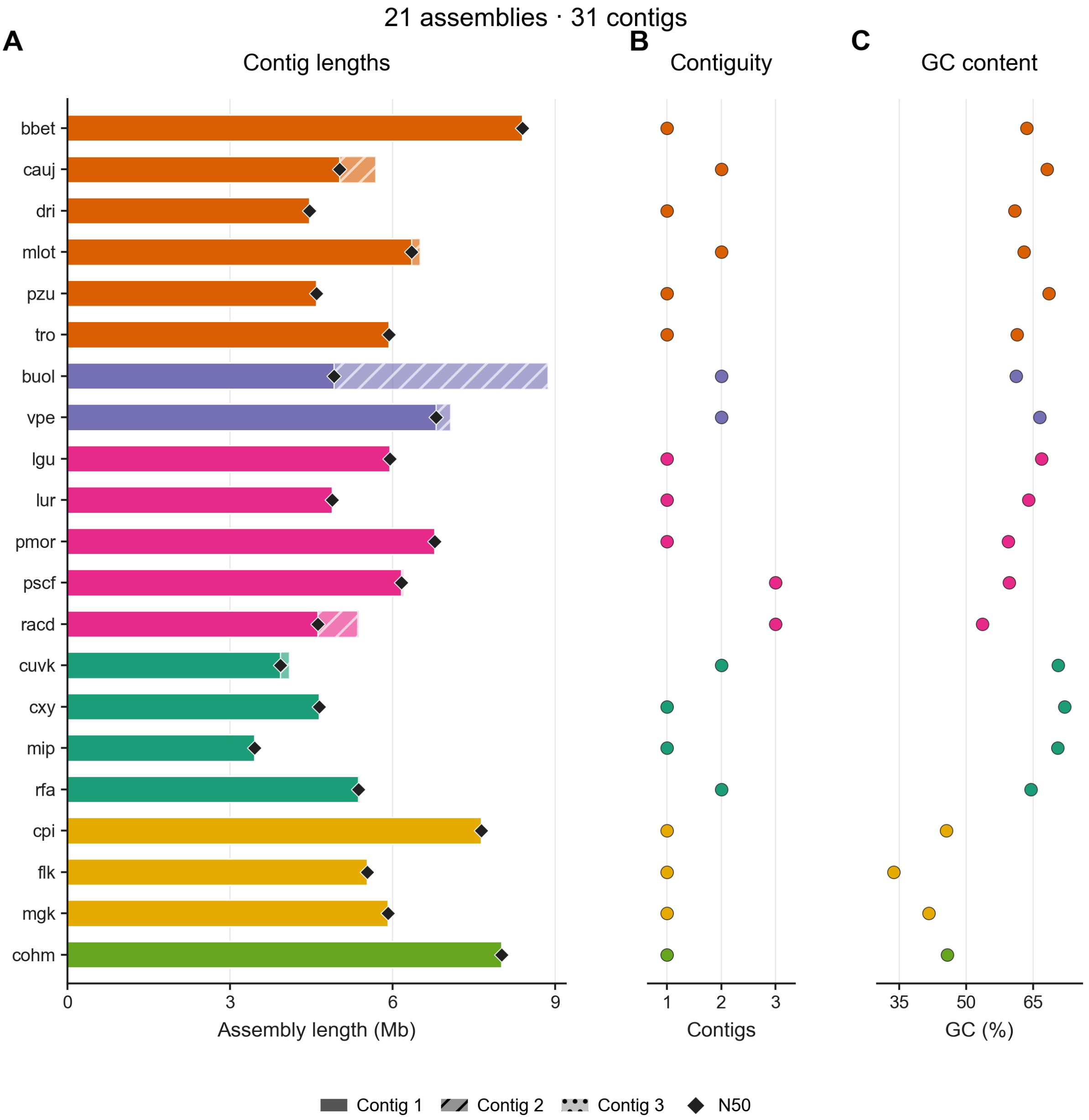


**Contiguity and composition of the genome assemblies used for model reconstruction**. (A) Total assembly length, partitioned into contigs ordered from longest to shortest. Black diamonds mark N50, the contig length at which the cumulative length of the ordered contigs reaches at least half of the total assembly length. (B) Number of contigs per assembly. (C) Guanine-cytosine (GC) content. The 21 assemblies contain 31 contigs: 13 assemblies have one contig, six have two and two have three. No N bases were recorded. Colors identify taxonomic plotting groups. Full values are provided in Supplementary Table S1.

### Supplementary Figure S12

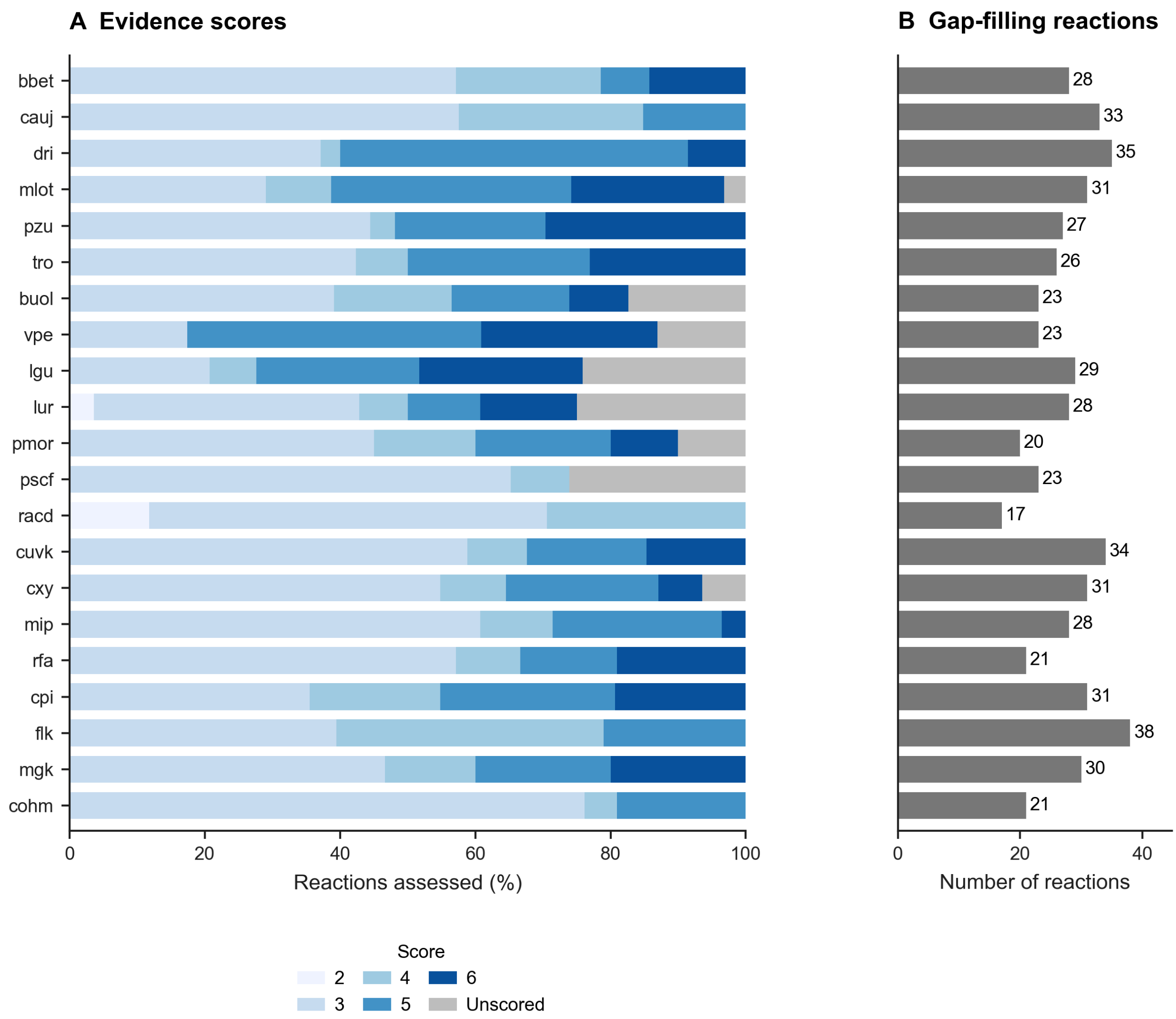


**Evidence supporting model gap filling**. (A) Distribution of evidence scores among reaction records assessed during gap filling, expressed as a percentage of the records for each strain. Scores follow the Enzyme Commission (EC) evidence rubric in Table 3. Shades of blue indicate recorded scores, and grey indicates no recorded score (not a score of zero). (B) Number of assessed reaction records per strain. The dataset contains 577 strain-reaction records: 545 have scores, and 32 do not.

### Supplementary Figure S13

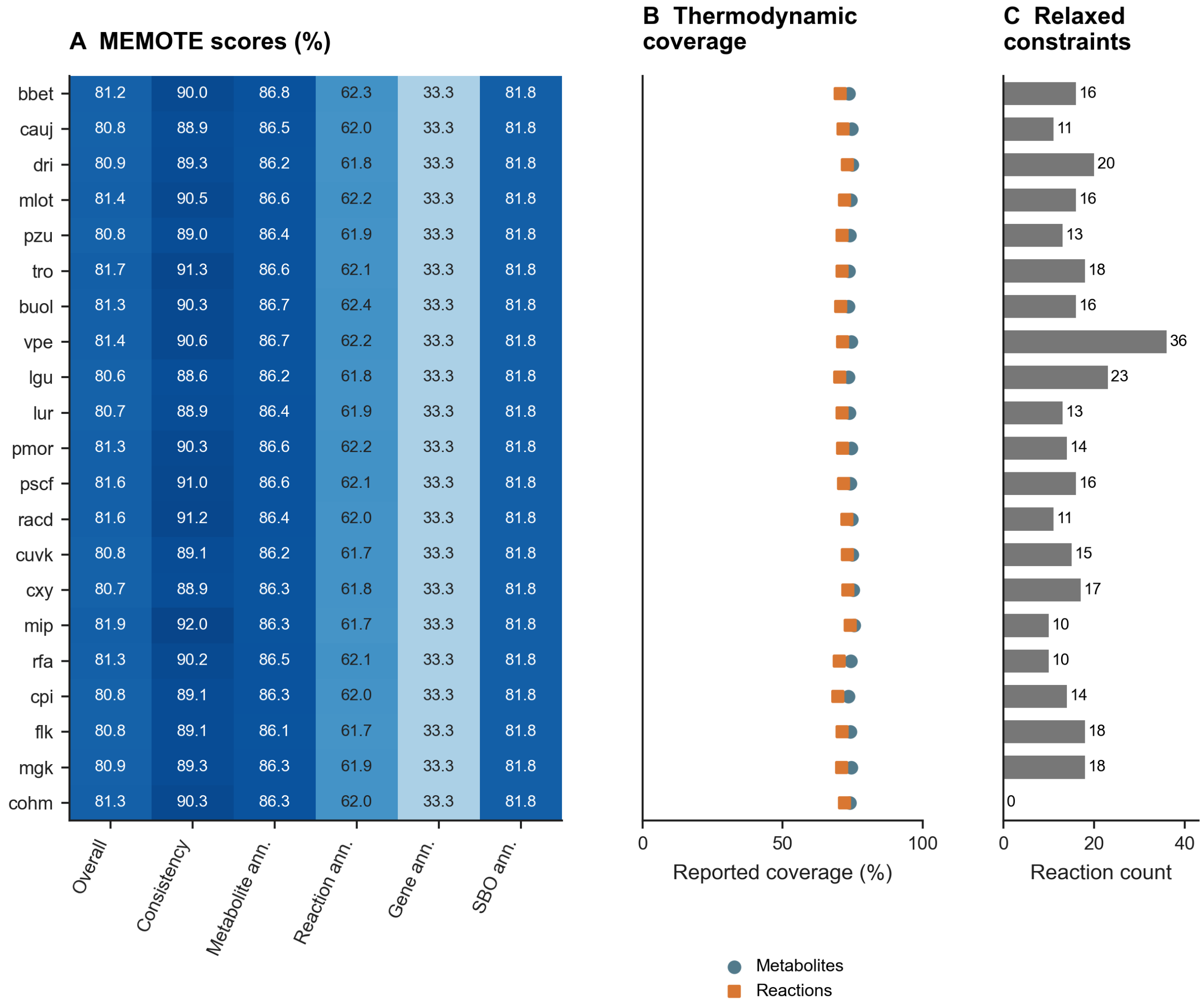


**Model quality scores and thermodynamic information**. (A) MEMOTE scores for the 21 genome-scale metabolic models. Columns show the overall score, model consistency, and metabolite, reaction, gene, and Systems Biology Ontology (SBO) annotation scores on a 0-100% scale. Cell labels give the scores, with darker blue indicating higher values. (B) Percentage of model metabolites (circles) and reactions (squares) with thermodynamic information. (C) Number of reaction constraints relaxed during thermodynamic curation of models in peptone-tryptone-yeast extract-glucose (PTYG) medium.

### Supplementary Figure S14 Sheet 1 of 2

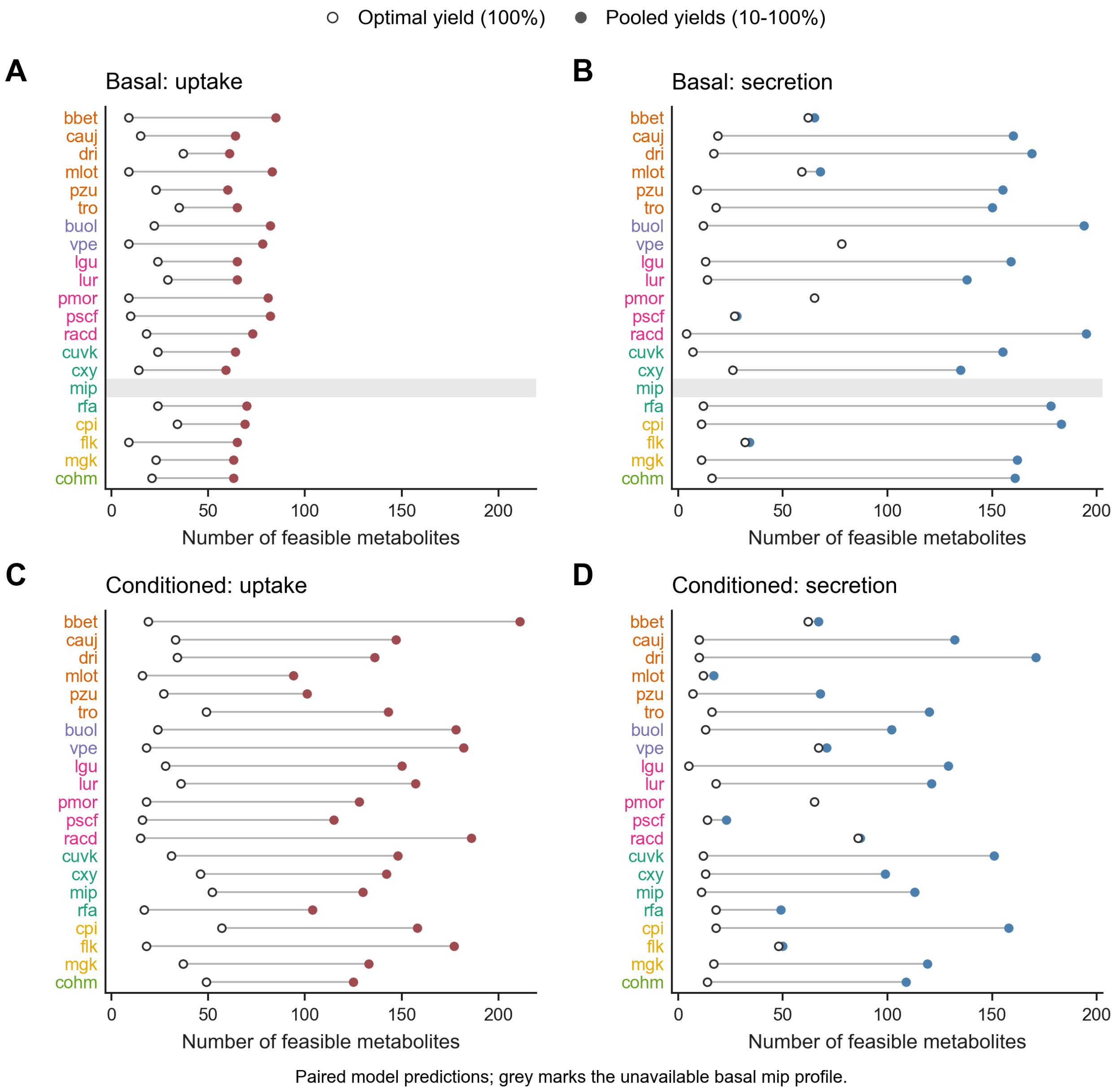


**Effect of biomass-yield aggregation on predicted exchange and niche overlap**. DiMEs (Decomposed in silico Minimal Exchanges) profiles at the optimal biomass yield (open circles) are compared with the union of profiles at yield levels from 10% to 100% of that optimum (filled circles). Biomass yield denotes biomass produced per amount of carbon taken up. Panels show basal uptake (A), basal secretion (B), conditioned uptake (C) and conditioned secretion (D). Lines connect the same strain; red and blue filled circles distinguish uptake and secretion, respectively. Grey marks the unavailable basal mip profile. The combined profiles represent alternative feasible exchanges, not exchanges that necessarily occur together. Strain-label colors identify taxonomic plotting groups.

### Supplementary Figure S14 Sheet 2 of 2

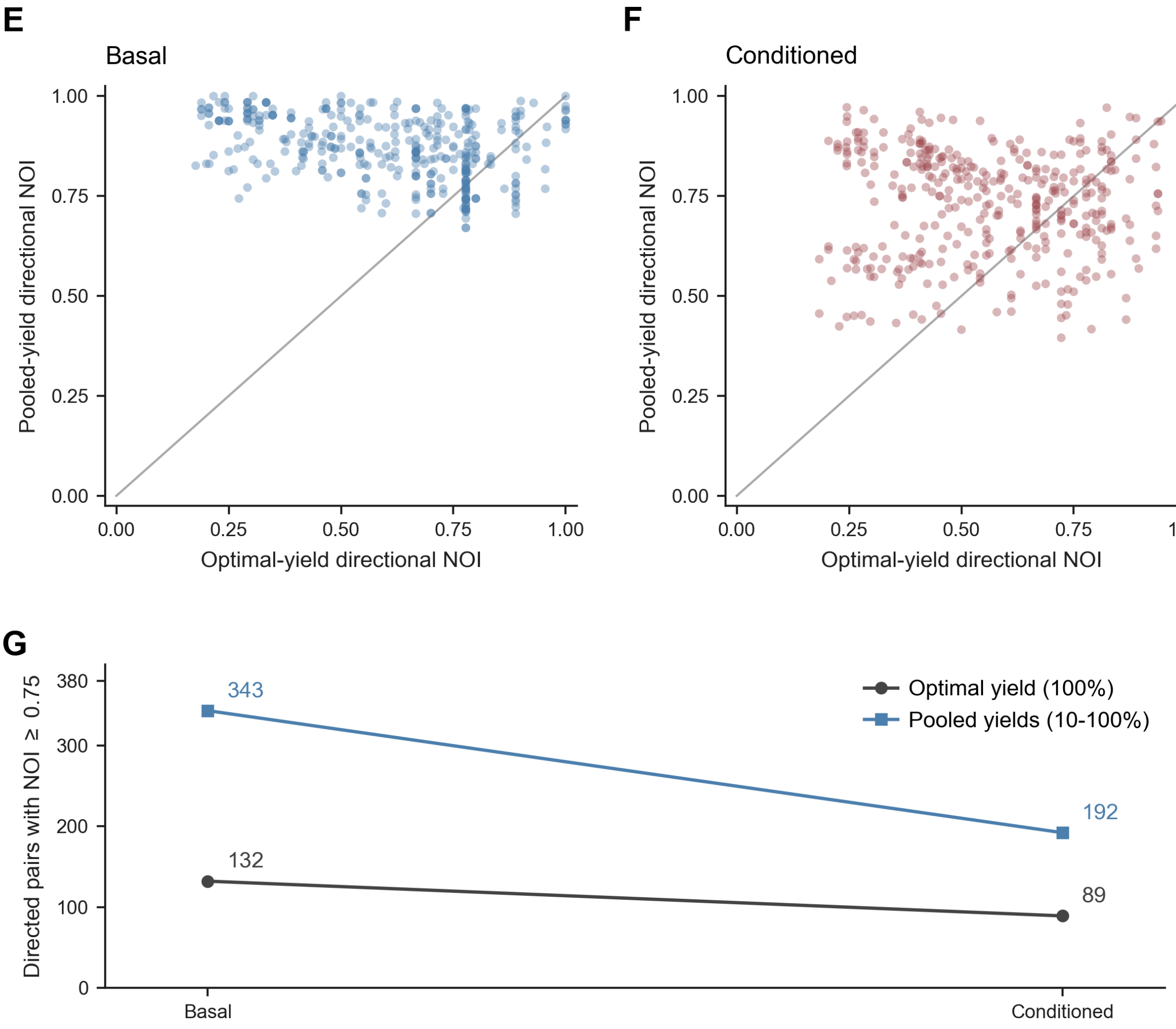


Matched 20 strains; 380 directed pairs in every comparison. Water is retained.

**Effect of biomass-yield aggregation on predicted exchange and niche overlap (continued)**. (E, F) Directional niche overlap in basal and conditioned environments, respectively, for the same 380 ordered pairs among 20 strains. The niche overlap index (NOI) is the fraction of the focal strain's feasible uptake set also present in the comparison strain's set. NOI = 1 therefore means complete coverage of the focal set, not necessarily identical uptake sets. Each point compares the optimal-yield NOI (x-axis) with the NOI calculated from profiles combined across 10-100% of optimal yield (y-axis); the diagonal marks equality. (G) Number of pairs with NOI ≥ 0.75. Conditioning changes this count from 132 to 89 at optimal yield and from 343 to 192 for the pooled yield.

**Supplementary Figure S15 Sheet 1 of 2**

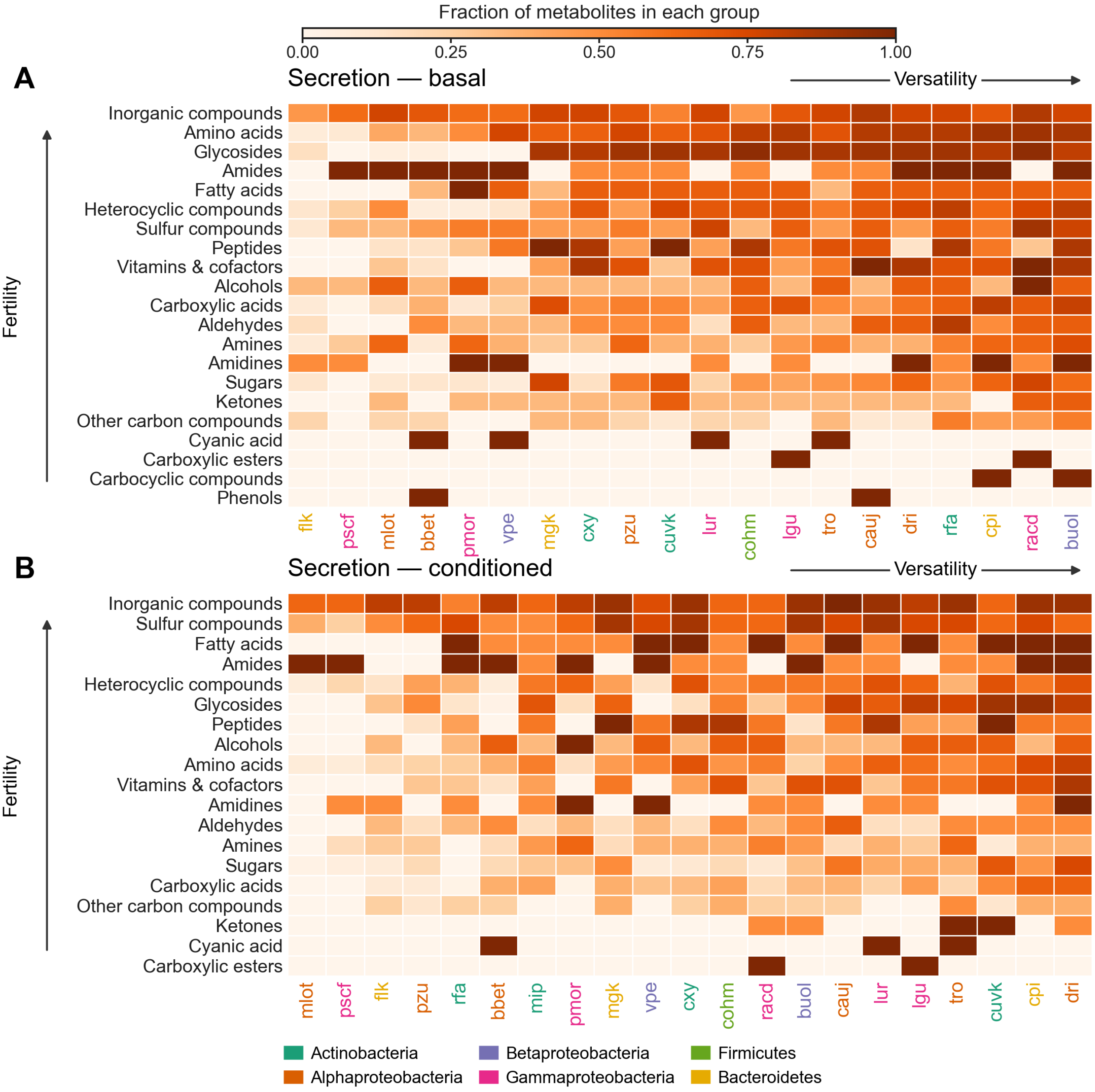


**Predicted exchange profiles**. Heatmaps show secretion in basal (A) and conditioned (B) environments; uptake is shown on the next sheet. The conditioned environment includes predicted community secretions. Rows represent metabolite classes, and columns represent strains. Cell color gives the fraction of compounds in each group that a strain can secrete, using as the denominator all compounds that at least one model can secrete in that environment. Arrows indicate increasing mean coverage across strains for each metabolite class (fertility, upwards) and across metabolite classes for each strain (versatility, rightwards). Basal profiles cover 20 strains (*mip* fails to grow), and conditioned profiles cover 21. Profiles combine solutions at 10-100% of optimal biomass yield. Strain-label colors identify the taxonomic groups in the legend.

**Supplementary Figure S15 Sheet 2 of 2**

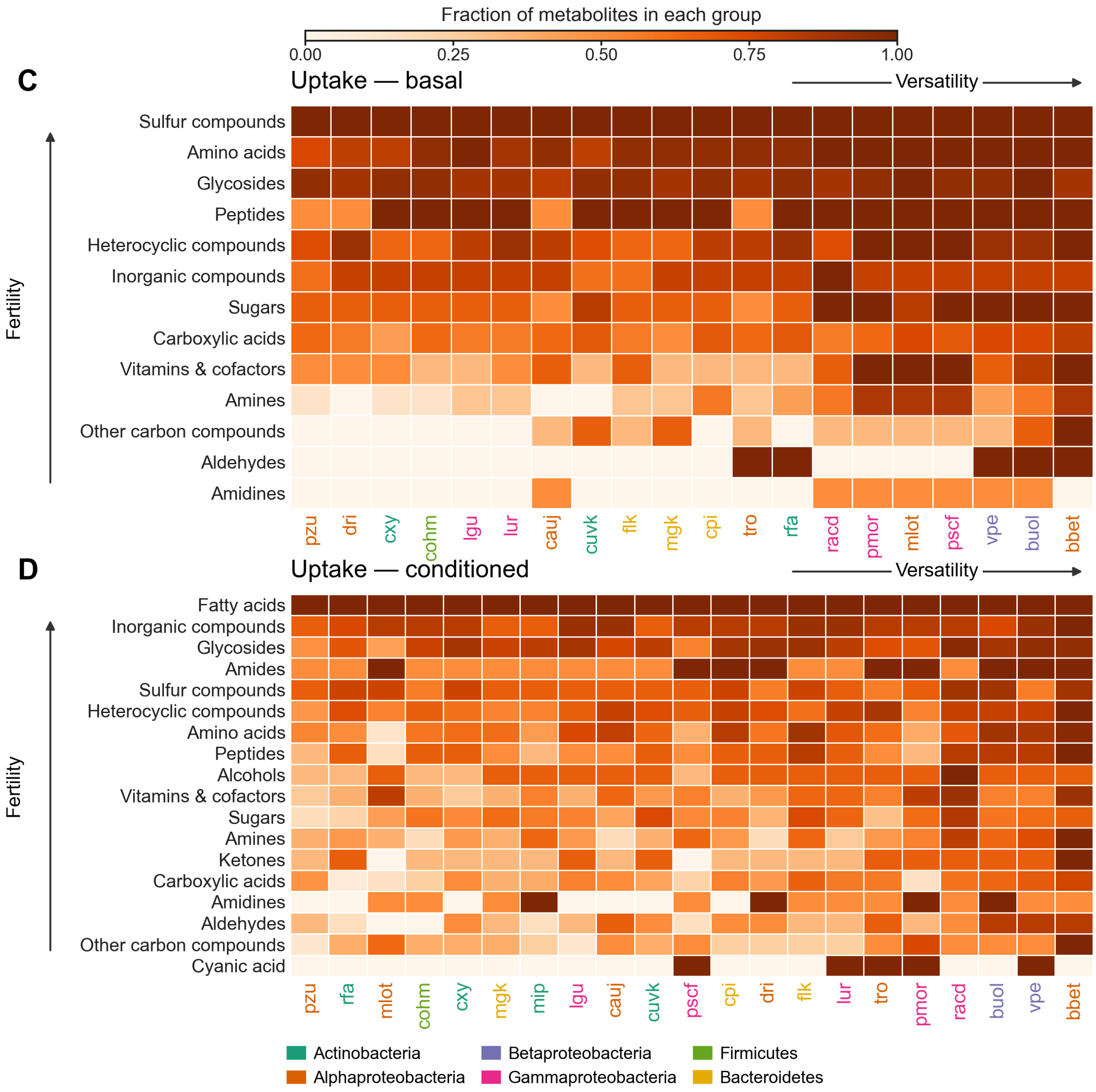


**Predicted exchange profiles (continued)**. Panels (C) and (D) show uptake in basal and conditioned environments, respectively. For each metabolite class, the denominator is the set of compounds that at least one model can take up in the corresponding environment. Darker cells indicate higher coverage on the 0-1 scale. Each panel has its own denominators and ordering. Fertility increases upwards and versatility rightwards, as defined on the preceding sheet. Basal profiles cover 20 strains (*mip* fails to grow), and conditioned profiles cover 21. Profiles combine solutions at 10-100% of optimal biomass yield. Strain-label colors identify the taxonomic groups in the legend.

### Supplementary Figure S16

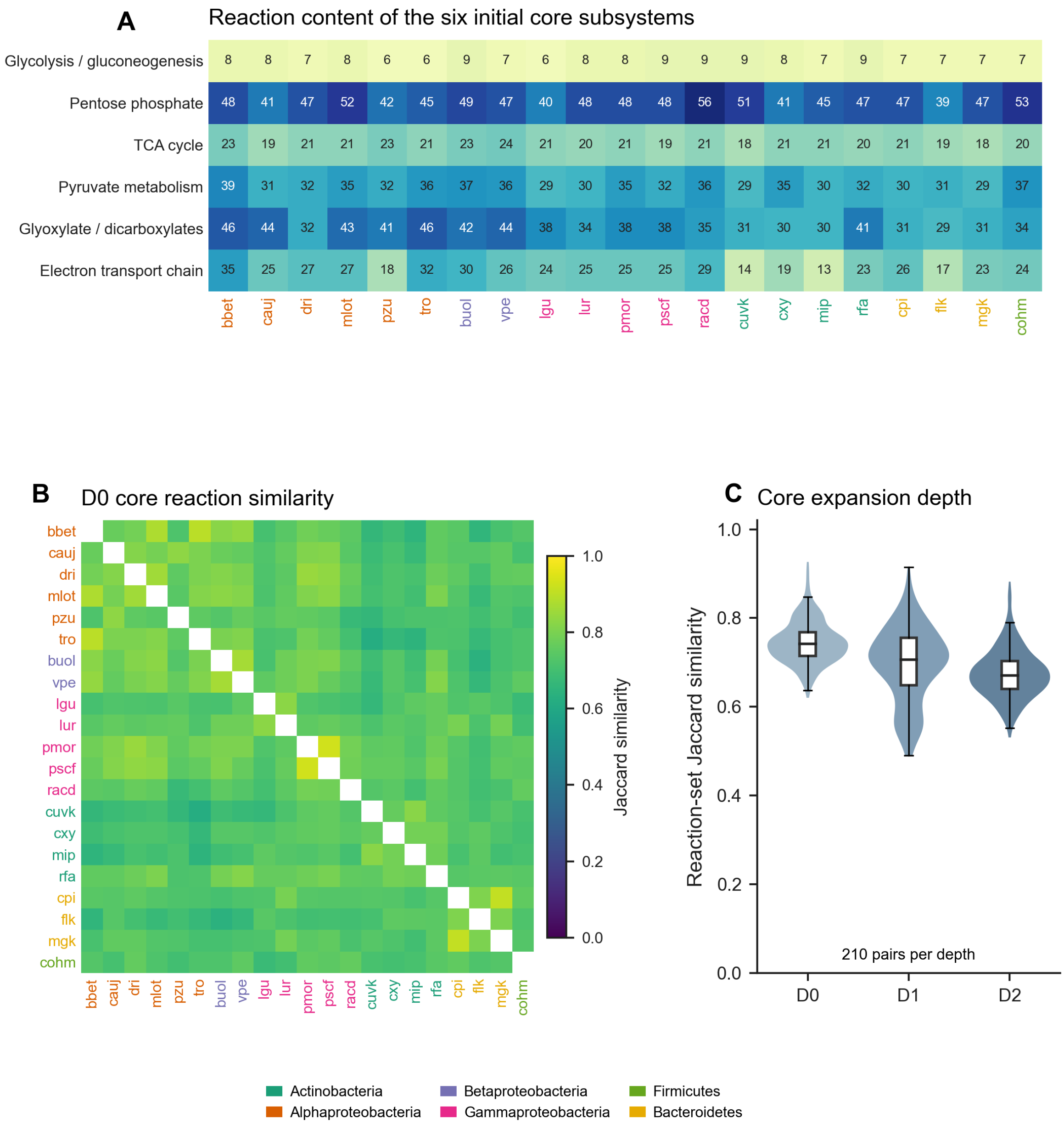


**Reaction content and similarity of central metabolic networks.** (A) Reaction counts in six initial core subsystems for each strain. (B) Pairwise Jaccard similarity of the initial core reaction sets (D0), defined as the number of shared reactions divided by the number of reactions present in either set. White diagonal cells omit self-comparisons. (C) Distributions of similarity for 210 strain pairs at the initial core (D0) and two successive expansions with connecting pathways (D1 and D2). Violin widths show the distributions; boxes span the interquartile range, centre lines mark medians, and whiskers extend to the most distant observations within 1.5 interquartile ranges. Taxonomic colors follow the legend.

**Supplementary Figure S17**

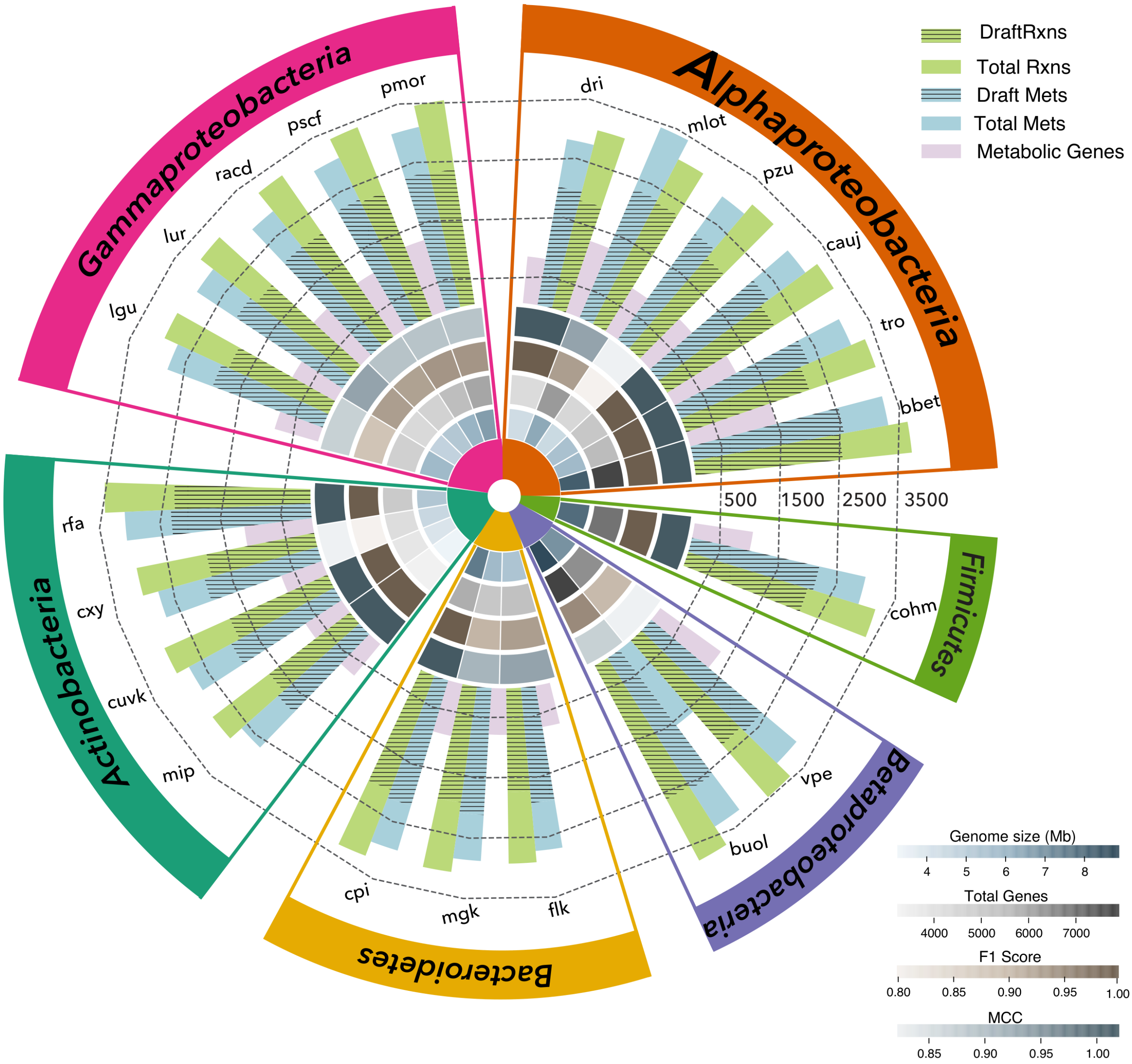


**Genome and metabolic-model characteristics of the 21 strains.** Each sector represents a strain, grouped by the taxonomic categories shown around the circle. Radial bars show reactions (green), metabolites (blue), and model-associated metabolic genes (purple). Hatched portions indicate draft-model reaction and metabolite counts; full bar lengths indicate curated-model totals. From the center outwards, shaded rings show genome assembly length in megabases (Mb), total annotated genes, F1 score, and Matthews correlation coefficient (MCC). Darker shading denotes larger values on the separate scales. F1 and MCC quantify agreement with the growth assays used for curation.

### Supplementary Figure S18 Sheet 1 of 2

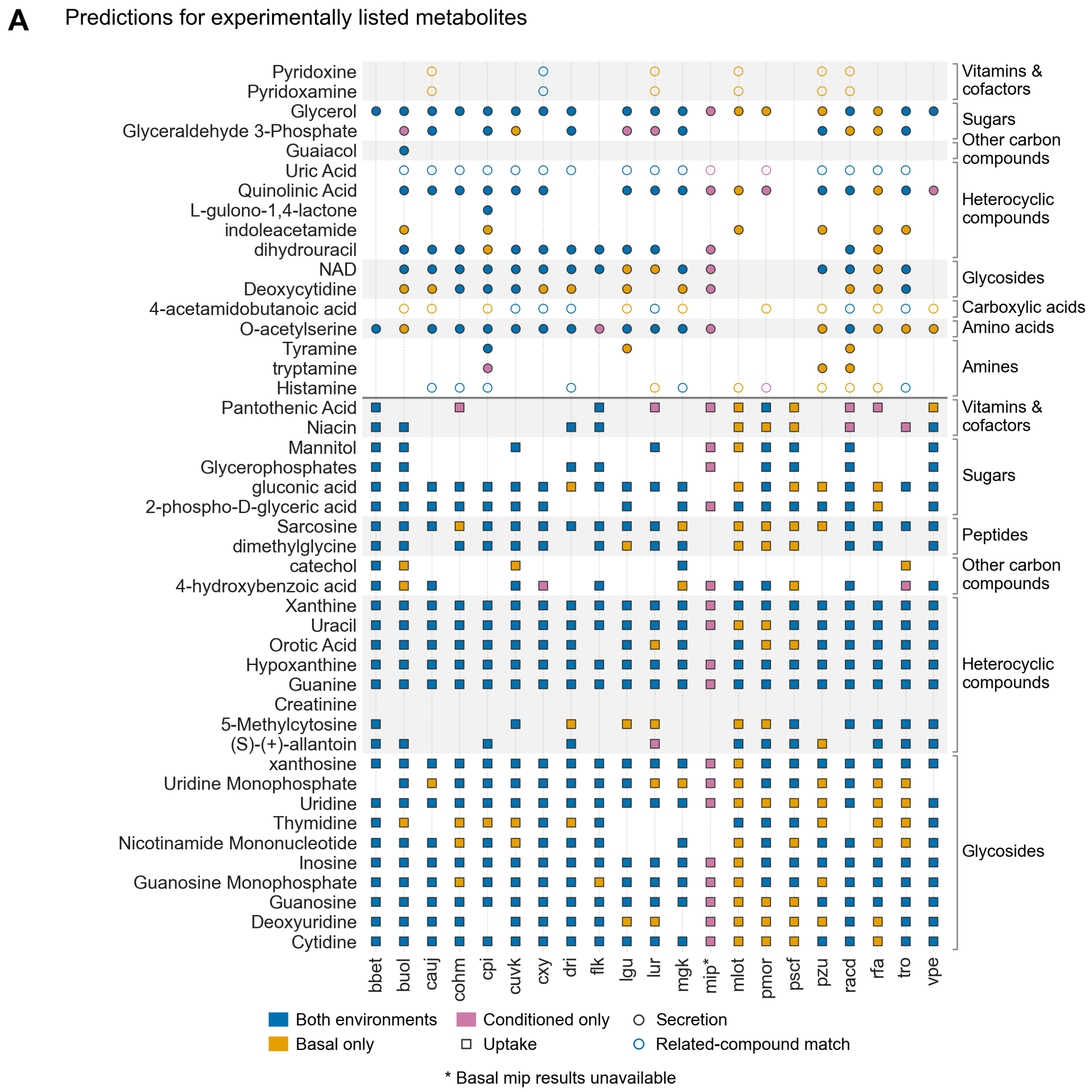


**Model-feasible exchange of metabolites listed in the experimental analysis**. The matrix spans panels (A) and (B) on successive sheets and compares model predictions with 73 metabolites listed for uptake and 17 listed for secretion in the experimental record. Columns represent strains and rows represent metabolites grouped by chemical class. Squares denote uptake and circles denote secretion. Blue, gold, and pink indicate feasibility in both environments, basal only, and conditioned only, respectively. Hollow circles indicate a related-compound match rather than an exact-metabolite match. Basal mip predictions are unavailable (*), so its conditioned markers do not establish a gain relative to basal. Predictions combine DiMEs (Decomposed in silico Minimal Exchanges) solutions across biomass yields of 10-100% of the optimum. The experimental lists do not identify individual producers or consumers. The strain-level assignments are model predictions.

### Supplementary Figure S18 Sheet 2 of 2

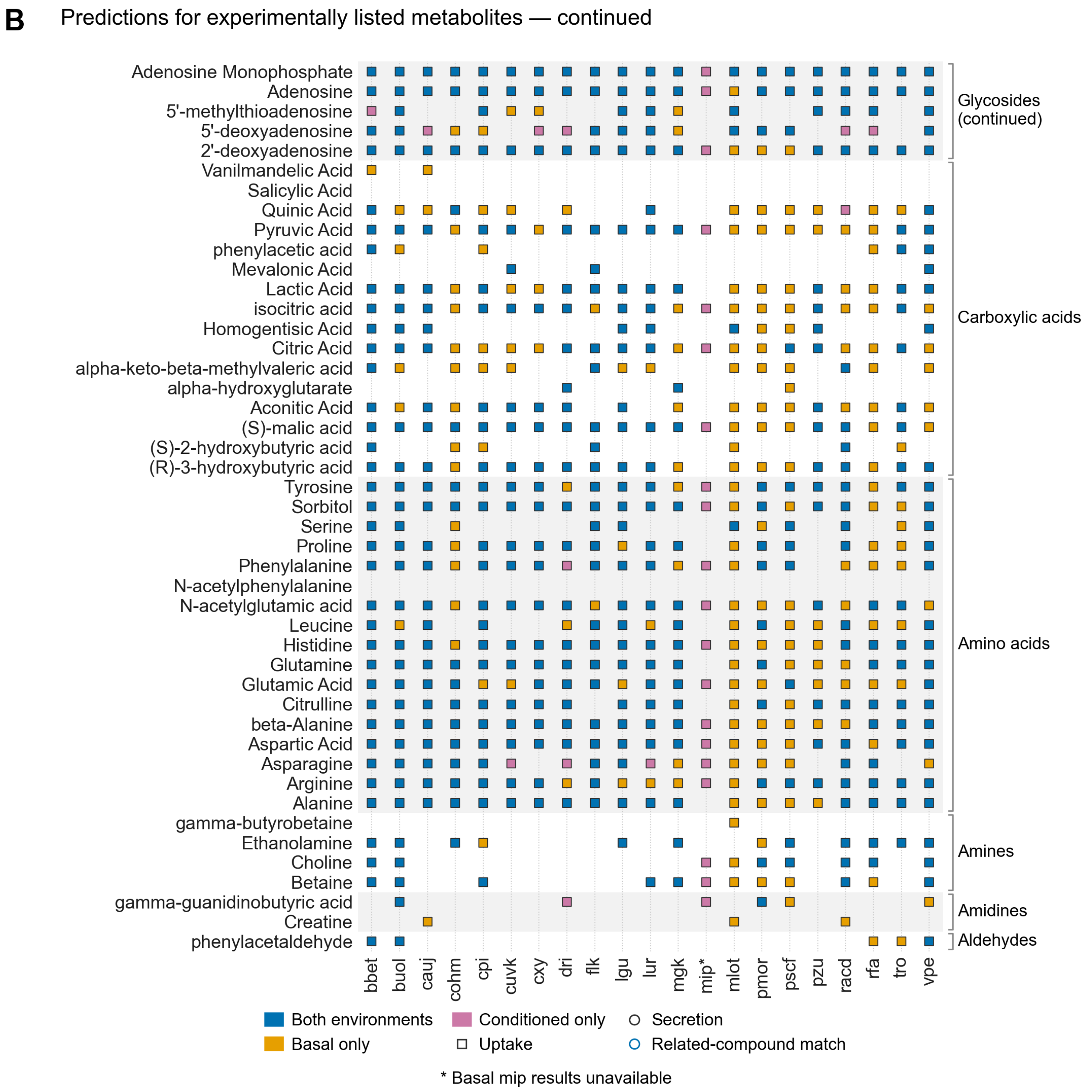


**Model-feasible exchange of metabolites listed in the experimental analysis (continued)**. Panel (B) continues the uptake matrix, beginning with the remaining glycosides. Squares indicate model-feasible uptake; blue, gold and pink denote feasibility in both environments, basal only and conditioned only, respectively. Blank cells indicate no model match, not an experimentally measured absence. Basal mip predictions are unavailable (*).

**Supplementary Table S1**

**PTYG medium composition**

| KEGG ID | Compound | KEGG ID | Compound |
|---|---|---|---|
| C00001 | H2O | C00159 | D-Mannose |
| C00007 | Oxygen | C00175 | Cobalt ion |
| C00009 | Orthophosphate | C00183 | L-Valine |
| C00011 | $CO_2$ | C00188 | L-Threonine |
| C00014 | Ammonia | C00238 | K+ |
| C00023 | Iron | C00242 | Guanine |
| C00025 | L-Glutamate | C00253 | Nicotinate |
| C00031 | D-Glucose | C00255 | Riboflavin |
| C00034 | Manganese | C00291 | Nickel |
| C00037 | Glycine | C00305 | Magnesium cation |
| C00038 | Zn2+ | C00314 | Pyridoxine |
| C00041 | L-Alanine | C00378 | Thiamine |
| C00047 | L-Lysine | C00380 | Cytosine |
| C00049 | L-Aspartate | C00407 | L-Isoleucine |
| C00059 | Sulfate | C00504 | Folate |
| C00062 | L-Arginine | C00698 | Cl- |
| C00064 | L-Glutamine | C00760 | Cellulose |
| C00065 | L-Serine | C00864 | Pantothenate |
| C00070 | Copper | C01330 | Na+ |
| C00072 | L-Ascorbic acid | C01413 | Cadmium |
| C00073 | L-Methionine | C01424 | Gallate |
| C00076 | Ca2+ | C01529 | Selenium |
| C00078 | L-Tryptophan | C05776 | Vitamin B12 |
| C00079 | L-Phenylalanine | C06267 | Vanadium |
| C00080 | H+ | C06268 | Chromium |
| C00082 | L-Tyrosine | C06696 | Lead |
| C00097 | L-Cysteine | C12505 | Magnesium sulfate |
| C00106 | Uracil | C13391 | Aluminum hydroxide |
| C00114 | Choline | C13409 | Titanium dioxide |
| C00120 | Biotin | C13881 | Ba2+ |
| C00123 | L-Leucine | C13884 | Sr2+ |
| C00135 | L-Histidine | C14818 | Fe2+ |
| C00147 | Adenine | C14819 | Fe3+ |
| C00148 | L-Proline | C18237 | Molybdoenzyme molybdenum cofactor |
| C00152 | L-Asparagine | | |

The 69 compounds included in the PTYG medium representation are listed with their KEGG compound identifiers. Read down the left pair of columns, followed by the right pair.

KEGG, Kyoto Encyclopedia of Genes and Genomes.

### Supplementary Table S2

### Biochemical composition of the basal and conditioned environments

| Biochemical group | Mapped from metabolomics | Basal additions | Basal total | Added by conditioning | Conditioned total |
|---|---|---|---|---|---|
| Amino acids and derivatives | 34 | 0 | 34 | 30 | 64 |
| Peptides | 0 | 0 | 0 | 5 | 5 |
| Sugars and derivatives | 8 | 0 | 8 | 35 | 43 |
| Other carboxylic acids and derivatives | 25 | 2 | 27 | 27 | 54 |
| Fatty acids and lipid-related compounds | 8 | 0 | 8 | 6 | 14 |
| Nucleobases, nucleosides and nucleotides | 26 | 0 | 26 | 24 | 50 |
| Vitamins, cofactors and related compounds | 8 | 3 | 11 | 7 | 18 |
| Inorganic compounds and ions | 0 | 20 | 20 | 10 | 30 |
| Other organic compounds | 11 | 0 | 11 | 26 | 37 |
| **Total** | **120** | **25** | **145** | **170** | **315** |

Counts summarise the input pools used for Decomposed in silico Minimal Exchanges (DiMEs). The basal pool comprises 120 compound identifiers mapped from sterile-soil metabolomics and 25 modelling additions. Conditioning adds 170 previously absent compounds from 255 unique predicted basal secretions; 85 secretions were already present in the basal pool. Each model compound identifier is counted once, including water, and assigned to one broad biochemical group for this table. These groups differ from the finer categories used in the exchange-profile figures. Compound-level assignments, grouping rules and source-name discrepancies are recorded in the accompanying Excel workbook. Counts describe model inputs, not uptake by every strain or independently confirmed experimental identifications.

**Supplementary Table S3**

**Statistics of the genome assemblies used for model reconstruction**

| Strain | Length (bp) | Contigs | N50 (bp) | GC (%) | Assembly accession |
|---|---|---|---|---|---|
| bbet | 8,393,826 | 1 | 8,393,826 | 63.59 | GCA_988226255 |
| cauj | 5,688,125 | 2 | 5,018,350 | 68.14 | GCA_988226285 |
| dri | 4,467,844 | 1 | 4,467,844 | 60.87 | GCA_988226505 |
| mlot | 6,503,638 | 2 | 6,354,096 | 62.98 | GCA_988226395 |
| pzu | 4,594,868 | 1 | 4,594,868 | 68.51 | GCA_988226365 |
| tro | 5,934,985 | 1 | 5,934,985 | 61.42 | GCA_988226465 |
| buol | 8,862,254 | 2 | 4,918,083 | 61.15 | GCA_988226305 |
| vpe | 7,064,762 | 2 | 6,806,102 | 66.48 | GCA_988226355 |
| lgu | 5,948,640 | 1 | 5,948,640 | 66.93 | GCA_988226385 |
| lur | 4,883,062 | 1 | 4,883,062 | 64.01 | GCA_988226445 |
| pmor | 6,779,898 | 1 | 6,779,898 | 59.44 | GCA_988226405 |
| pscf | 6,198,775 | 3 | 6,159,625 | 59.63 | GCA_988226325 |
| racd | 5,369,342 | 3 | 4,617,620 | 53.66 | GCA_988226485 |
| cuvk | 4,090,513 | 2 | 3,931,117 | 70.60 | GCA_988226495 |
| cxy | 4,643,243 | 1 | 4,643,243 | 72.07 | GCA_988226295 |
| mip | 3,452,946 | 1 | 3,452,946 | 70.48 | GCA_988226335 |
| rfa | 5,387,795 | 2 | 5,368,684 | 64.54 | GCA_988226545 |
| cpi | 7,634,451 | 1 | 7,634,451 | 45.52 | GCA_988226415 |
| flk | 5,531,431 | 1 | 5,531,431 | 33.78 | GCA_988226525 |
| mgk | 5,917,343 | 1 | 5,917,343 | 41.63 | GCA_988226455 |
| cohm | 8,008,983 | 1 | 8,008,983 | 45.77 | GCA_988226435 |

Metrics were calculated from the final FASTA files used for model reconstruction. Length is the sum of contig lengths. N50 is the contig length at which the cumulative length, after ordering contigs from longest to shortest, reaches at least half of the assembly length. GC content is the percentage of G and C bases among all A, C, G and T bases. None of the 21 assemblies contains N bases. Assembly accessions follow manuscript Table 1; bp, base pairs.
